# Synthetic inhibitors of pregnancy-associated plasma protein-A regulate IGF-1 signaling *in vitro* and *in vivo*

**DOI:** 10.64898/2026.09.30.755852

**Authors:** Anna Kertisová, Shang-Wu Shih, Nicholas S. Kirk, Jan Pícha, Agáta Rücklová, Jaroslav Kozák, Karolína Mrzílková, Marta Lubos, Eva Tloušťová, Miloš Buděšínský, Cunming Duan, Jiří Jiráček, Irena Selicharová

## Abstract

Dysregulation of insulin-like growth factor (IGF) signaling contributes to diverse human diseases. Therapeutic strategies have largely targeted the IGF-1 receptor, but the complexity of the IGF system has limited their clinical efficacy. Pregnancy-associated plasma protein-A (PAPP-A) is a secreted metalloproteinase that regulates local IGF signaling through proteolytic cleavage of IGF-binding proteins (IGFBPs) and represents an attractive therapeutic target. Here, we report the first synthetic PAPP-A inhibitors designed to target the enzyme active site using an IGFBP-5-derived anchor sequence. Replacing the scissile bond with phosphinate or phosphonate moieties yielded potent and selective PAPP-A inhibitors. Cryo-electron microscopy revealed the molecular basis of inhibition and key active-site interactions. The compounds inhibited cleavage of IGFBP-4 and IGFBP-5, two physiological PAPP-A substrates. They suppressed IGF-1 signaling in A549 lung cancer cells, with efficacy comparable to the endogenous PAPP-A inhibitor stanniocalcin-1. They also inhibited Papp-aa-dependent IGF signaling and cell proliferation in wild-type and *stc1a-/-* mutant zebrafish *in vivo* in a dose-dependent manner. Together, these findings establish the first class of synthetic, selective PAPP-A inhibitors capable of modulating IGF signaling. They provide new chemical tools for studying and modulating pathological IGF signaling and offer new directions for developing PAPP-A-targeted therapeutic strategies.

## Introduction

The insulin/insulin-like growth factor (IGF) signaling axis is an evolutionarily ancient and highly conserved hormonal system that regulates metabolism, somatic growth, tissue homeostasis, and lifespan across animal species^1^. In humans, this axis comprises insulin and two insulin-like growth factors (IGF-1 and IGF-2), which act through activation of tyrosine kinase receptors, including insulin receptor isoforms A and B (IR-A and IR-B), the IGF-1 receptor (IGF-1R), and their hybrid forms^2^. While insulin is secreted exclusively from pancreatic β-cells in response to elevated glucose levels and is rapidly cleared from circulation, IGFs are produced by multiple tissues in both endocrine and paracrine modes, and their circulating levels and bioavailability are tightly controlled by six IGF-binding proteins (IGFBP1–6)^3^. These IGFBPs prolong IGF’s half-life in the circulation, affect IGF transport to target tissues, and control IGF availability to the IGF-1R in target cells^4^. Proteolytic cleavage of IGFBPs is major mechanism for local IGFs activity^2,3^.

Among the best-characterized IGFBP proteases is pregnancy-associated plasma protein-A (PAPP-A). PAPP-A is a member of the metzincin metalloproteinase superfamily^5^. PAPP-A together with its paralog PAPP-A2 (Ref.^6^) forms the pappalysin family. To date, the confirmed substrates of PAPP-A are IGFBP-4 in complex with IGF-1 or IGF-2, IGFBP-5, and partially IGFBP-2, whereas PAPP-A2 cleaves IGFBP-5 and partially IGFBP-3 (Ref.^7^). Physiologically, PAPP-A activity is inhibited by several endogenous proteins, including stanniocalcins (STC1 and STC2) and the pro-form of eosinophil major basic protein (proMBP)^8–10^.

The highest PAPP-A expression has been detected in human placental trophoblasts^11^ and circulating PAPP-A levels have long been used as a biomarker in prenatal screening programs for early risk assessment^12^. Beyond pregnancy, PAPP-A functions postnatally and mediates the release of IGFs from the IGF/IGFBPs complexes to regulate somatic growth, tissue remodeling, and repair^13^. Genetic deletion of PAPP-A activity in mice has been linked to improved health span and lifespan extension^14,15^. On the other hand, elevated PAPP-A expression or activity has been associated with cancer progression^16^, cardiovascular disease^17^, kidney disease^18,19^ or fibrosis^20^. PAPP-A binds to the cell surface through its short consensus repeat (SCR) modules and was proposed to act as a local regulator of IGF signaling by cleaving IGFBP-4 at proximity of the IGF-1R^21^. Consequently, inhibition of PAPP-A has emerged as a promising strategy for tissue-selective modulation of local IGF activity. Targeting IGF-1R signaling itself has long been considered an attractive anti-cancer strategy; however, therapeutic approaches based on IGF-1R–neutralizing antibodies or tyrosine kinase inhibitors have shown limited clinical success due to insufficient efficacy, insulin receptor cross-reactivity and resistance mechanisms^22,23^. In this context, modulation of IGF bioavailability through inhibition of PAPP-A represents an attractive alternative further supported by its cell-specific secretory nature and extracellular localization in tumor microenvironments^24^, which may facilitate pharmacological targeting.

Despite the growing recognition of PAPP-A as a biologically and therapeutically important regulator of the IGF axis, there is currently a striking lack of pharmacological tools capable of selective modulation of PAPP-A activity *in vivo*. To date, studies investigating the role of PAPP-A in disease progression and aging have relied on genetic knockout models^14^ and neutralizing antibodies^8,25^.

Although these approaches have demonstrated the therapeutic potential of PAPP-A inhibition, they remain limited for mechanistic studies and translational development. In particular, the absence of low-molecular-weight PAPP-A inhibitors restricts the ability to investigate acute and tissue-specific modulation of PAPP-A activity. Therefore, the development of selective small-molecule PAPP-A inhibitors is critically needed both as chemical biology tools and as potential therapeutic agents targeting IGF-related diseases and aging-associated pathologies.

Here we present for the first time synthetic active-site-directed PAPP-A inhibitors with *K*_i_ in nanomolar range capable of inhibiting cleavage of both IGFBP-4 and IGFBP-5. The inhibitors were based on the anchor peptide sequence of IGFBP-5 (Ref.^26^) where the scissile bond was replaced by non-cleavable pseudopeptide moiety. We used phosphinic or phosphonic groups serving as tetrahedral transition-state analogs that mimic the structure of the water-attacked scissile peptide bond during hydrolysis by Zn^2+^ metalloproteases^27–29^. We demonstrated potency of our inhibitors to inhibit PAPP-A activity *in vitro* and *in vivo* and we also solved the structure of inhibitor-bound PAPP-A revealing the structural basis of inhibition. These inhibitors provide valuable tools for further investigation of PAPP-A biology and may support future development of PAPP-A-targeted therapies.

## Results

### Synthesis of peptides

All peptides used in the study were synthesized by standard procedures of solid-phase peptide synthesis. List of the peptides and synthesis details are presented in **Supporting Information, Scheme 1 and Figures S1, S4-S6**. We synthesized a new fluorogenic substrate (Q-BP5) for a quenched fluorescence assay to measure PAPP-A activity. The Q-BP5 was based on the anchor peptide, IGFBP-5 (Ref.^139–163^) (numbering according to the UniProt P24593 including the signal 20 amino-acid peptide) from the structure (PDB 7UFG) (Ref.^26^) extended of eight residues following the scissile bond. The peptide was modified according to a previously established quenched fluorescence assay^30^. Next, we synthesized a fluorogenic peptide (FS-6)^31^ that is cleaved by several matrix metalloproteinases (MMPs), to test specificity and selectivity of our inhibitors.

The peptide IGFBP-5^139-171+Y^ **Figure 1** (peptide **1**) was used as a control substrate and as a template for synthesis of the inhibitors. C-terminal tyrosine residue was added to enable precise concentration measurements using absorbance at 280 nm.

**Figure 1:**
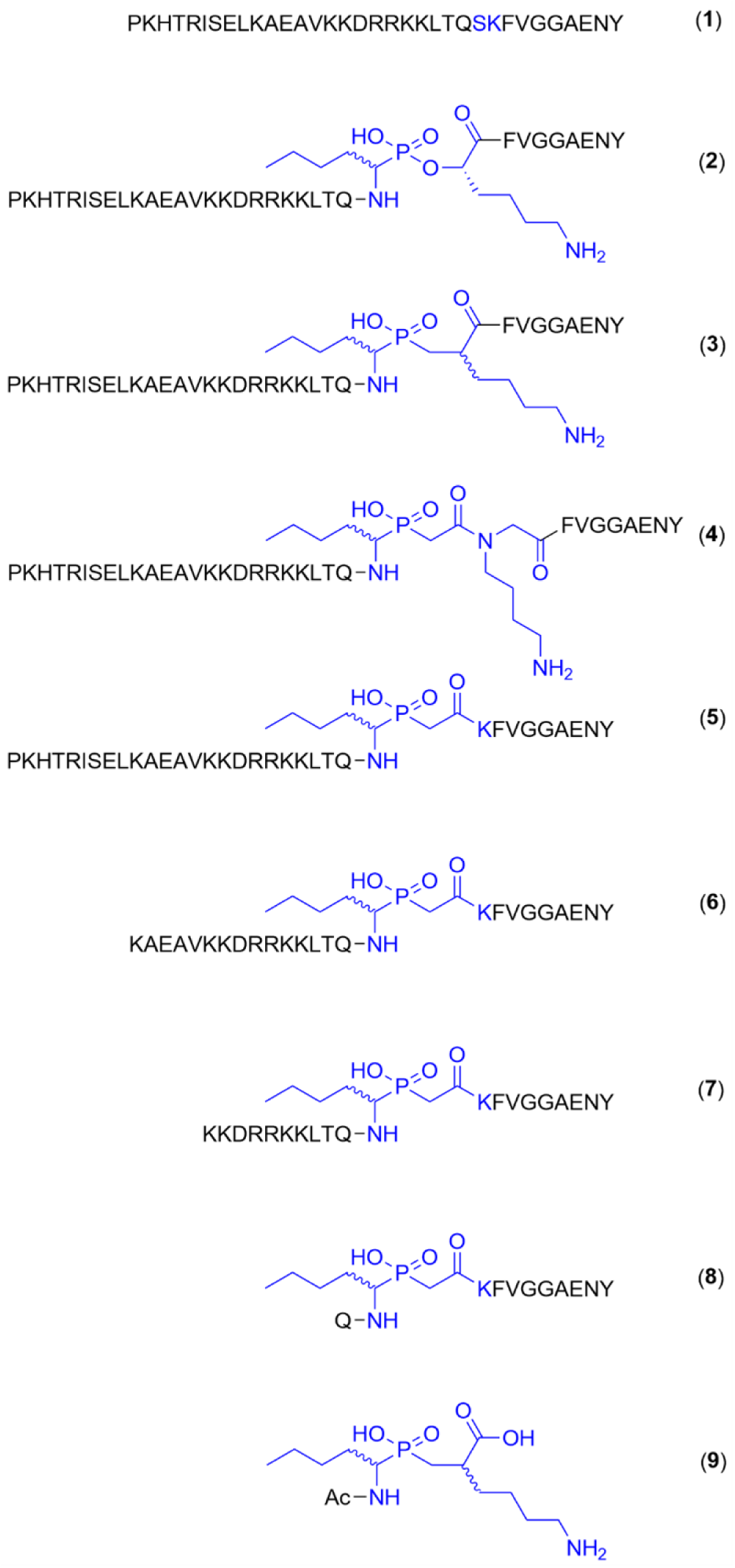
Structures of the synthesized PAPP-A substrate peptide **1** which corresponds to the IGFBP-5^139-171+Y^ sequence and inhibitor pseudopeptides **2**-**9** derived from peptide **1**. The native scissile Ser– Lys dipeptide and the pseudopeptide moieties replacing it are shown in blue.

### Synthesis of inhibitors

We synthesized protected pseudo-dipeptide precursors **I**–**IV** designed to mimic the tetrahedral transition-state intermediate of the scissile peptide bond between norleucine and lysine residues. The pseudopeptide precursors were incorporated into peptide **1** and its truncated variants at the PAPP-A cleavage site, replacing IGFBP-5 residue S^163^ or the S^163^–K^164^ dipeptide (**Figure 1**). Precursor **I** contained a phosphonate motif, -PO(OH)-O-, and was used for the synthesis of inhibitor **2**. Precursors **II**–**IV** contained a phosphinate motif, -PO(OH)-CH_2_-, and were used for the synthesis of inhibitors **3**– **9** (**Supporting Information, Schemes 2–7** and **Figures S2** and **S3**). Some of the reactions employed for syntheses of intermediate and final compounds were not stereospecific. Separation of the resulting stereoisomers was not feasible under the conditions employed. Consequently, the pseudopeptides were tested as unresolved mixtures of diastereoisomers, comprising four diastereoisomers for inhibitors **3** and **9** and two for inhibitors **2** and **4**–**8** (**Figure 1**).

### Inhibition of PAPP-A activity in quenched fluorescence assay

Active PAPP-A was isolated from the medium of transfected HEK293T cells (**Figure S7 A**). Cleavage characteristics and inhibitory potencies of the tested compounds were determined using the quenched fluorescence assay^30^ with Q-BP5 as a substrate. The measured IC_50_ and calculated *K*_i_ of the inhibitors are shown in **Table 1**. The Q-BP5 was cleaved with *K*_m_ = 4.65 ± 0.28 μM (**Figure 2A**). The non-modified control peptide **1** competed in the assay with IC_50_ = 452 ± 24 nM, equivalent to a *K*_i_ of about 144 nM after recalculation using Cheng-Prusoff equation. The pseudopeptides **2** and **5** were more than twofold stronger and the pseudopeptides **3** and **4** almost 20-fold more potent than peptide **1**. This means that introducing a phosphinate group -PO(OH)-CH_2_-at the cleavage site substantially increased the affinity of the peptide, and that the affinity depended on the type of bond between the phosphinate and lysine side chain. The representative inhibition curves are shown in **Figure 2B**. Inhibitory potencies of all compounds at 0.1 μM and 100 μM are shown in (**Figure S7 B**).

**Figure 2:**
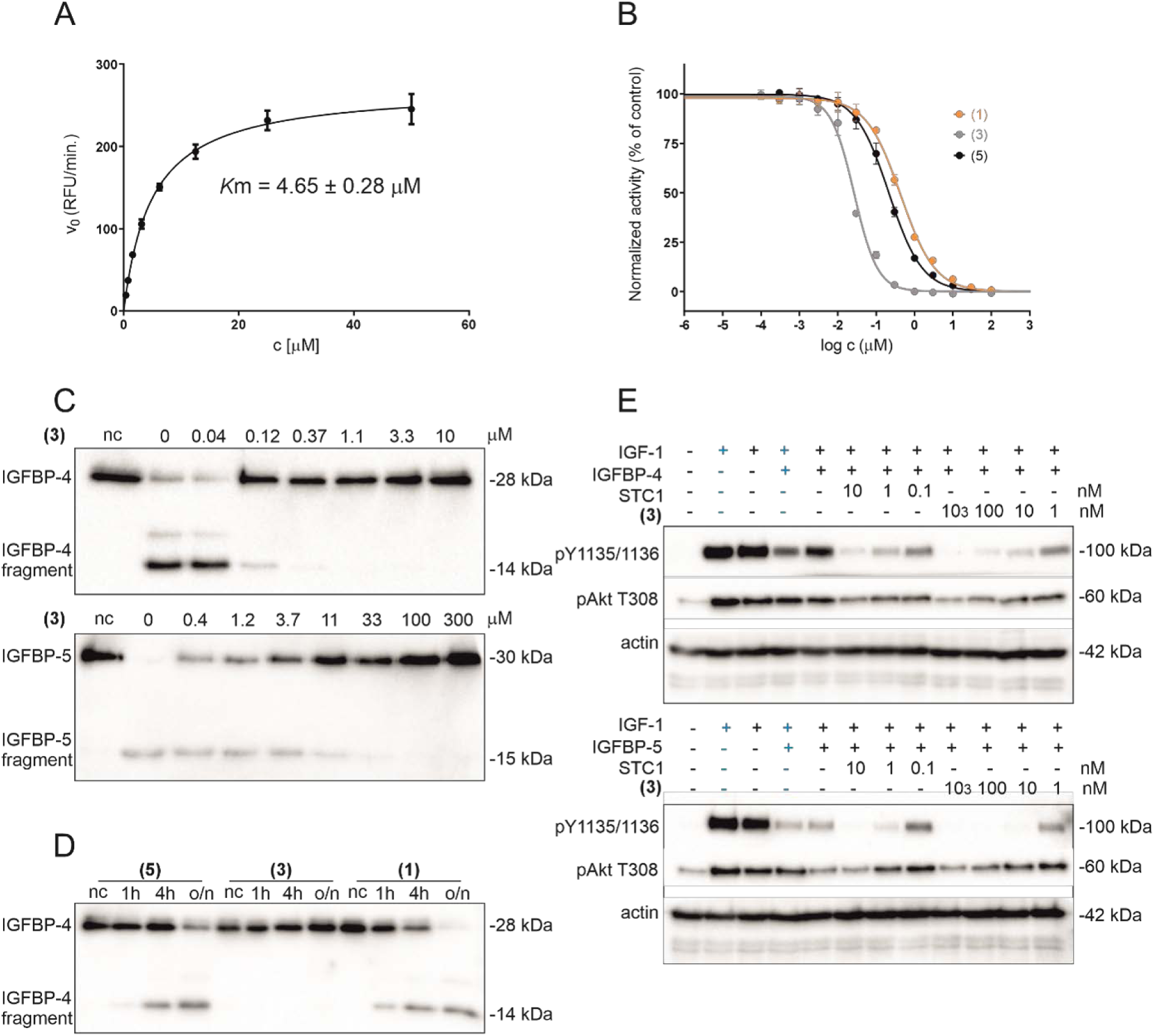
Inhibition of PAPP-A activity. (**A**) Determination of PAPP-A *K*_m_ using the quenched fluorescence assay. (**B**) Inhibition of PAPP-A activity in the quenched fluorescence assay in competition with peptide **1 (1)**, pseudopeptide **3 (3)** or pseudopeptide **5 (5)**. Data are shown as mean and SD (n=3). (**C**) Inhibition of cleavage of IGFBP-4 and IGFBP-5 with pseudopeptide **3 (3)**. The proteins and their cleaved fragments were detected using western blot with anti-IGFBP-4 antibody and anti-6 × His tag primary antibody for IGFBP-5 (nc – negative control without enzyme) (**D**) Time course effects of IGFBP-4 cleavage by peptide **1(1)** and pseudopeptides **3 (3),** and **5 (5)** (3.3 μM), (o/n – overnight incubation). (**E**) Inhibition of IGF-1R signaling in A549 cell line with pseudopeptide **3 (3)**. Cells were pre-treated with the pseudopeptide **3** or STC1 of indicated concentrations for 1 h. IGF-1 (10 nM) was preincubated with IGFBP-4 or IGFBP-5 (20 nM) for 30 min and added to the cells for two hours (black signs) or 20 min (blue signs). (-) means that the compound was not present, (+) means treatment with IGF-1 alone or in complex with IGFBPs. Stimulation of IGF-1R signaling was analyzed using standard western blotting. Details are described in methodology section.

**Table 1:** Inhibition constants of compounds determined using quenched fluorescence assay. Sequences of the peptide **1** and pseudopeptides **2-9** are shown in **Figure 1**. Measured values of IC_50_ were recalculated to *K*_i_ using Cheng-Prusoff equation (*K*_i_ = IC_50_/(1+ [S]/*K*_m_), where [S] = 10 μM and *K*_m_ = 4.65 μM).

| Compound | Source peptide | IC<sub>50</sub> [μM] | K<sub>i</sub> [μM] |
| --- | --- | --- | --- |
| **1** | IGFBP-5<sup>139-171+Y</sup> | 0.452 ± 0.024 (3) | 0.144 ± 0.008 |
| **2** | IGFBP-5<sup>139-171+Y</sup> | 0.123 ± 0.010 (3) | 0.039 ± 0.003 |
| **3** | IGFBP-5<sup>139-171+Y</sup> | 0.027 ± 0.001 (3) | 0.009 ± 0.0003 |
| **4** | IGFBP-5<sup>139-171+Y</sup> | 0.022 ± 0.002 (3) | 0.007 ± 0.0007 |
| **5** | IGFBP-5<sup>139-171+Y</sup> | 0.193 ± 0.021 (3) | 0.061 ± 0.007 |
| **6** | IGFBP-5<sup>148-171+Y</sup> | 32.7 ± 0.8 (3) | 10.4 ± 0.3 |
| **7** | IGFBP-5<sup>153-171+Y</sup> | 231 ± 30(3) | 73.3 ± 9.6 |
| **8** | IGFBP-5<sup>162-171+Y</sup> | 92.7 ± 17.2 (3) | 29.4 ± 5.5 |
| **9** | IGFBP-5<sup>163-164</sup> | 1039 ± 114 (3) | 330 ± 36 |

We also tested the pseudo-dipeptide block itself (compound **9**), STC1, a natural PAPP-A inhibitor, and batimastat, a matrix metalloproteases (MMP) non-specific inhibitor. The pseudo-dipeptide block **9** was not active. STC1 was tested only at 0.1 μM concentration and inhibited cleavage of the peptide Q-BP5 by lowering, but not eliminating, the PAPP-A activity to 52% in agreement with data from previous studies^32^. Batimastat was tested at 100 μM and showed limited potency when lowering the PAPP-A activity only to 62% (**Figure S7 B**).

It was known from previous studies that amino-acids distant from the scissile bond are necessary for substrate binding. We synthesized shortened versions of the pseudopeptide **5** covering IGFBP-5 sequences 148-171+Y (**6**), 153-171+Y (**7**) and 162-171+Y (**8**), (**Figure 1**). All of the peptides were less potent than the full-length pseudopeptide **5** (**Table 1**). Surprisingly, the pseudopeptide **6** starting with the crucial residue K^148^ (Ref.^26^) had two orders of magnitude reduced activity compared to the full-length pseudopeptide **5** suggesting that the amino acids N-terminal from K^148^ substantially contribute to anchor peptide binding. We performed molecular dynamics (MD) modeling to gain deeper mechanistic insight into substrate binding (**Supporting Information and Table S1**). Basic residues at positions R^143^, K^148^, K^158^ and sequence motif KDR^154-156^, were confirmed as main anchoring points within the enzyme-substrate interface in agreement with previous studies^26,33^ and the cryo-EM structure solved in this work (see below). Residues surrounding the cleavage site Q^162^ and K^164^ were suggested to play a role in orienting the cleaved peptide bond into a catalytically competent position through bending the peptide. IGFBP-5 is also a substrate of PAPP-A2. PAPP-A2 (Sino Biological) was able to cleave the Q-BP5 substrate (**Figure S8 A**). However, a readable effect in the quenched fluorescence assay was detected only at 100 nM concentration of the enzyme which was not feasible for routine testing. PAPP-A2 was previously shown to cleave IGFBP-5 about 38-fold less efficiently than PAPP-A^34^. In **Figure S8 B**, we show cleavage of IGFBP-5 by PAPP-A2 using western blot. We tested pseudopeptide **3** in the western blot assay with PAPP-A2 but could not show clear inhibitory effect (**Figure S8 C**).

### Inhibition of PAPP-A cleavage of IGFBP-4 and IGFBP-5

The pseudopeptides **2** and **5** had very similar potency. The same applied for the pair of pseudopeptides **3** and **4**, but at markedly higher level (**Table 1**). We therefore tested pseudopeptides **3** and **5** as representative compounds in subsequent assays. Their effects in inhibiting the cleavage of full length IGFBP-4 and IGFBP-5 was assessed. Representative western blot of the cleavage of IGFBP-4 in complex with IGF-1 and cleavage of IGFBP-5 by pseudopeptide **3** is shown in **Figure 2C** and pseudopeptide **5** is shown in **Figure S9 A**. Clear inhibition of cleavage of IGFBP-4 in complex with IGF-1 by pseudopeptide **3** was observed at 0.12 μM while similar extent of inhibition of IGFBP-5 cleavage was achieved at 3.7 μM.

The non-modified peptide **1** competed with the substrates in the assays but during the prolonged overnight treatment we observed sustained inhibition of cleavage of IGFBP-4 by pseudopeptide **3** while the cleavable peptide **1** was consumed (**Figure 2D**).

### Expression of STC-PAPP-A-IGFBP axis genes in the A549 cell line

We tested the inhibitors in a cell-based assay and also *in vivo* to reveal their biological potency. The inhibition of naturally secreted PAPP-A was tested using the lung epithelial cell line A549 that was shown to produce PAPP-A in relatively large quantity^25^.

We analyzed mRNA expression of *PAPP-A*, *PAPP-A2*, *IGF*s, *IGFBP*s and *STC*s in the A549 cell line to discern possible interference of the components produced by the cells after incubation in serum-free medium (starved) and treatment with IGF-1 (**Figure S10**). The starved cells significantly increased their expression of *PAPP-A* and decreased expression of *STC1* and *STC2* compared to the control cells signifying that proportion of active PAPP-A was increased under starvation stress. The trend was partially reversed by IGF-1 treatment. The cells expressed *IGFBP-2* and *IGFBP-4* while expression of *IGFBP-5* was negligible. The expressions of *IGFBP-2* and *IGFBP-4* were not affected and thus were not expected to interfere with the signaling assay. *IGF-1* expression was diminished under the IGF-1 treatment. The *IGF-2* and *PAPP-A2* expression was negligible. The increase in PAPP-A expression in starving conditions was confirmed at the protein level in cell lysates using western blot (**Figure S10 G**).

### Inhibition of IGF-1 signaling in the A549 cell line

To study the effect of PAPP-A inhibition on IGF-1 signaling, A549 cells were serum starved overnight. Naturally secreted PAPP-A accumulated in the medium. The cells were pre-treated with the inhibitors or STC1 and then treated with IGF-1 pre-incubated with IGFBP-4 or IGFBP-5 at 1:2 molar ratio.

Activation of the IGF-1 signaling pathway, i.e. autophosphorylation of the IGF-1R and phosphorylation of Akt, was followed by western blot (**Figure 2E** shows effect of pseudopeptide **3**, pseudopeptide **5** is shown in **Figure S9 B**, and direct comparison of pseudopeptides **3** and **5** is in **Figure S9 C,D**). The inhibitors were able to inhibit IGF-1 signaling in a concentration dependent manner, the pseudopeptide **5** was markedly weaker than pseudopeptide **3** in the assay (**Figure S9 C,D**) in agreement with the quenched fluorescence assay. The effect of pseudopeptide **3** was comparable with STC1.

### Inhibition of IGF signaling in zebrafish *in vivo*

We next investigated the effects of the pseudopeptides **3** and **5** on PAPP-A activity *in vivo* using the Tg(igfbp5a:GFP) reporter zebrafish line. In this *in vivo* model, a population of epithelial cells known as Na^+^/K^+^-ATPase-rich ionocytes, or NaR cells, is labeled with green fluorescent protein (GFP)^35^.

Previous studies have shown that quiescent NaR cells can be induced to re-enter the active cell cycle by transferring fish into an induction embryo medium containing low levels of Ca²⁺. This quiescence-to-proliferation switch is mediated by the activation of local IGF signaling through Papp-aa-mediated Igfbp5a degradation (the zebrafish orthologs of human PAPP-A and IGFBP-5)^36–38^. Under normal Ca²⁺ conditions, Papp-aa proteinase activity is suppressed by high levels of endogenous Stc1a (the zebrafish ortholog of human STC1), which sequesters IGFs in the IGF/Igfbp5a complex and prevents NaR cell proliferation. Under low-Ca²⁺ conditions, *Stc1a* expression is diminished and Papp-aa-mediated Igfbp5a degradation is increased. This increases IGF bioavailability and IGF signaling, thereby promoting NaR cell proliferation^36–38^. Thus, this system provides an excellent whole-animal platform to test whether the pseudopeptides can inhibit Papp-aa activity *in vivo*.

As shown in **Figure 3A,B** and **S11**, compared with Tg(*igfbp5a*:GFP) fish maintained in control medium, fish transferred to induction medium had significantly more NaR cells. Consistent with previous reports, this increase in NaR cell proliferation was abolished by treatment with BMS-754807, an IGF-1R inhibitor. The addition of pseudopeptides **3** and **5** decreased NaR cell proliferation in a dose-dependent manner (**Figure 3A,B** and **S11**). At 10 and 20 µM, pseudopeptide **3** completely abolished NaR cell proliferation. Pseudopeptide **5** at 10 µM or higher also had a significant inhibitory effect (**Figure 3A,B** and **S11**).

**Figure 3.**
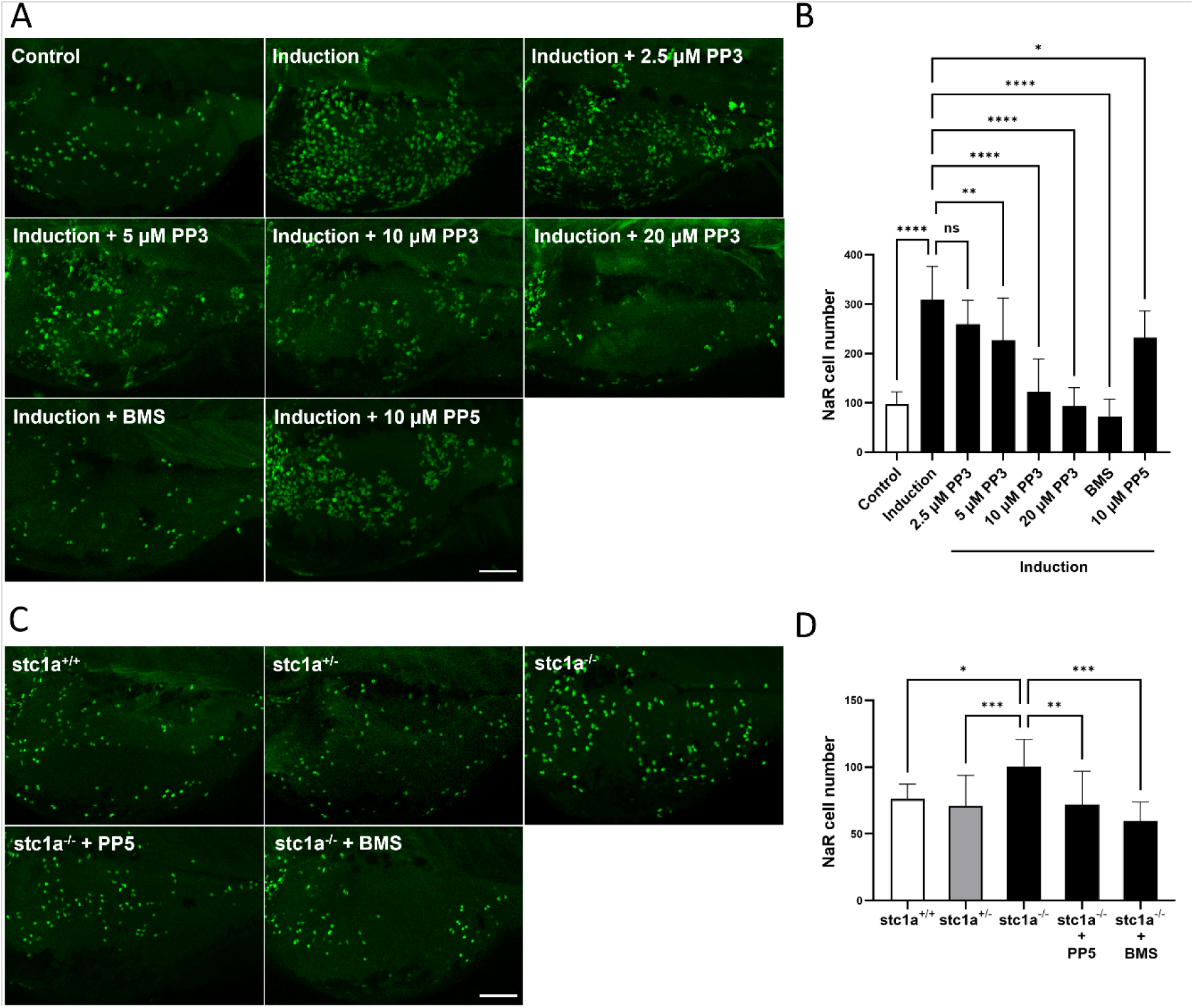
Inhibition of Papp-aa activity *in vivo*. (**A–B**) Pseudopeptides **3** and **5** inhibit IGF signaling-mediated NaR cell proliferation *in vivo*. Tg(igfbp5a:GFP) fish were raised to 3 days post-fertilization (dpf) and then transferred to control or induction medium. Pseudopeptide **3** (PP3), pseudopeptide **5** (PP5), or BMS-754807 (BMS) was added at the indicated concentrations, and treatment solutions were changed daily. Two days later, GFP expressing NaR cell numbers in the yolk-sac region of each fish were quantified. Representative images of the yolk-sac region are shown in **A**, and quantified results are shown in **B**. Images are lateral views with anterior to the left and dorsal up. Scale bar = 100 μm. n = 9–13 fish/group. (**C–D**) Pseudopeptide **5** (PP5) abolishes cell proliferation induced by the loss of Stc1a. Progeny from *stc1a⁺/⁻* intercrosses were raised to 3 dpf and then treated with 10 μm PP5 or 0.5 μm BMS. After 2 days of treatment, NaR cell numbers in the yolk-sac region in each fish were quantified, and fish were subsequently genotyped individually. Representative images are shown in **C**, and quantified results are shown in **D**. n = 9–24 fish/group. Images are lateral views with anterior to the left and dorsal up. Scale bar = 100 μm. Data in **B** and **D** are presented as mean ± SD. ns, not significant; \**p* < 0.05, \*\**p* < 0.01, \*\*\**p* < 0.001, \*\*\*\**p* < 0.0001.

We next used *stc1a⁻/⁻* null mutant fish to further test the inhibitors. Previous studies have shown that Papp-aa activity and NaR cell proliferation are constitutively elevated in *stc1a⁻/⁻* null mutant fish^38^. If these pseudopeptides inhibit Papp-aa activity and IGF signaling *in vivo*, they should reduce NaR cell proliferation in *stc1a⁻/⁻* fish. To test this, progeny from *stc1a⁺/⁻* intercrosses in the Tg(igfbp5a:GFP) background were treated with pseudopeptide **5**. BMS-754807 was used as a positive control. The results showed that pseudopeptide **5** reduced the number of NaR cells in *stc1a⁻/⁻* mutant fish to a level comparable to that in the wild-type control fish (**Figure 3C,D**). Collectively, these data suggest that these inhibitors suppress PAPP-A/Papp-aa activity and IGF signaling-stimulated cell proliferation *in vivo*.

### Cryo-EM structure of PAPP-A in complex with inhibitor 3

To understand the structural basis of inhibition by the pseudopeptides we determined a 2.6 Å cryo-EM reconstruction of pseudopeptide **3** bound to near-full length, active PAPP-A (Sino Biological, cat: 17290-H44H), resolving the laminin G-like domain (LG), metalloprotease domain (MP) and central domain (M1/M2) from P^92^ to S^1191^ (**Figure 4A** and **S12, Table S2**). Pseudopeptide **3** was resolved from K^140^ to G^167^ (IGFBP5 numbering), with residues to T^161^ remaining essentially identical to the previously published IGFBP5 bound inactive PAPP-A structure (**Figure 4C** and **S143A**)^26^. The pseudo-dipeptide block and residues F^165^ and V^166^ were completely resolved in our structure, forming a pseudo-beta sheet along the PAPP-A E^444^ - A^449^ strand, with backbone hydrogen bonds from the pseudo-lysine carbonyl to A^446^ and V^166^ amide nitrogen to E^444^, coordinating the F^165^ and V^166^ side chains into hydrophobic pockets containing L^445^ and F^470^ within the MP domain, respectively (**Figure 4B** and **S 13C**). While the remaining C-terminal residues were too flexible to be modelled unambiguously, density was observed tracing the C-terminal tyrosine to a pocket formed by P^527^ and P^537^ within the MP domain.

**Figure 4:**
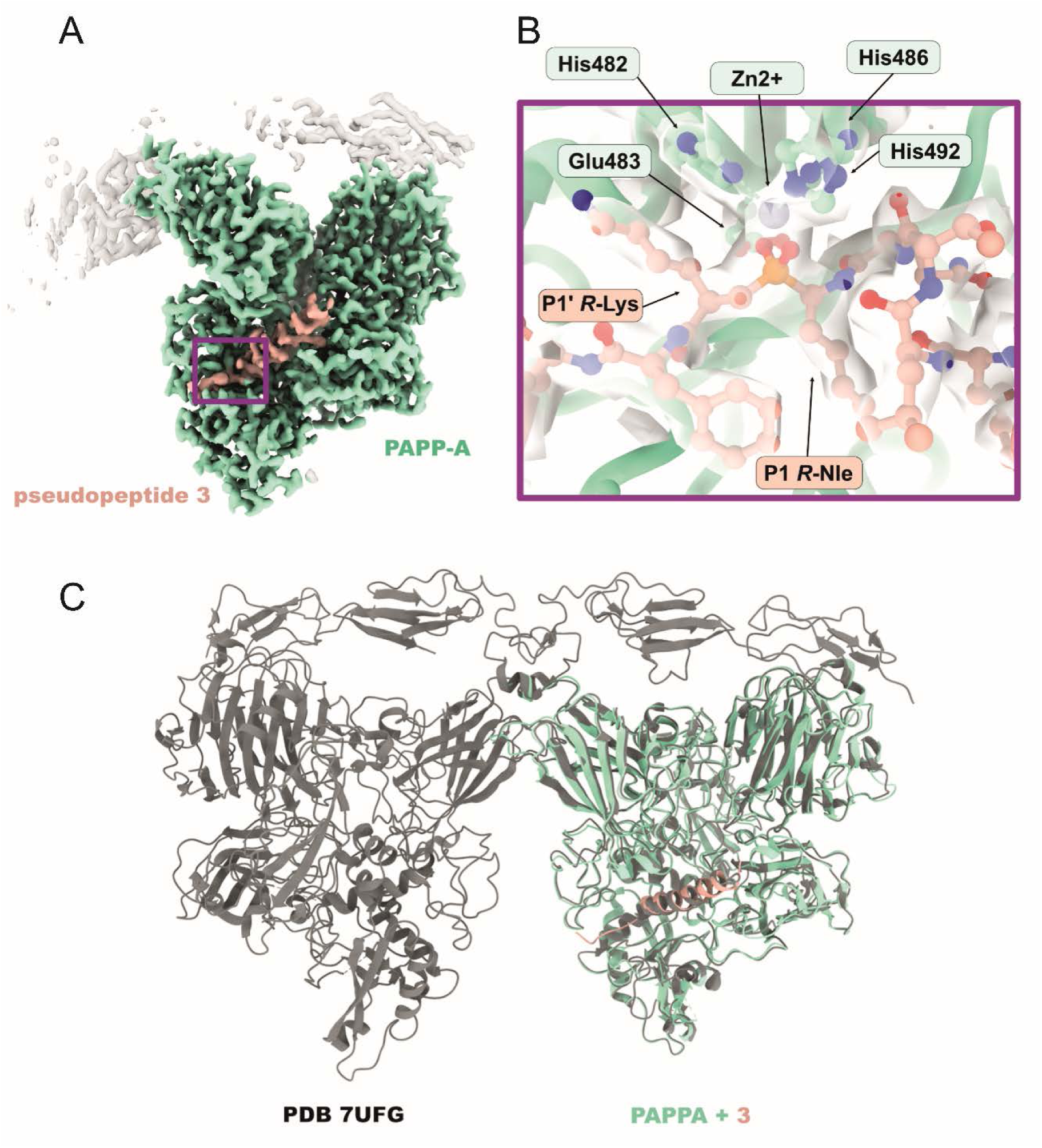
Cryo-EM structure of pseudopeptide 3 bound to PAPP-A. (**A**) Cryo-EM map of pseudopeptide **3** (salmon) bound to PAPP-A (turquoise). Map was sharpened using EMReady (46). (**B**) Inlay showing catalytic site of PAPP-A with key residues in atomic detail with associated cryo-EM density. (C) Overlay of cryo-EM structure with PDB 7UFG.

The expected tetrahedral phosphinate arrangement was observed at the catalytic site, with the Zn^2+^ ion coordinated by the phosphinate oxygens, the norleucine side chain occupying the S1 pocket and the lysine side chain as expected in the S1’ pocket. Despite being stereochemically impure, only the (*R*,*R*)-diastereoisomer of the pseudo-dipeptide phosphinate moiety of pseudopeptide **3** was observed in the structure.

### Cytotoxicity, specificity and stability of the inhibitors in plasma

We evaluated cytotoxicity of the pseudopeptides **3**, **5**, the shortened pseudopeptides **6**-**8** and the pseudo-dipeptide block **9** using six various cell lines. None of the cell lines showed significant decline in cell viability after treatment with the compounds at 100 μM concentrations for 3 days (**Figure S14**). Next, we tested cross-reactivity with other metzincin metalloproteases, namely matrix proteases MMP-1 and MMP-7. We synthesized a fluorogenic substrate FS-6 (Ref.^31^) and measured *K*_m_ for MMP-1 and MMP-7 (**Figure S15 A,D**). The PAPP-A inhibitors or their fragments at 100 μM did not show any inhibition of cleavage of FS-6 by MMP-1 or MMP-7 (**Figure S15 C,F**) and *vice versa*, PAPP-A did not cleave the FS-6 (**Figure S15 G**). Batimastat effectively inhibited both MMP-1 and MMP-7 (**Figure S15 B,E**) while it was almost inactive against PAPP-A at the dose used (**Figure S7 B**).

The model pseudopeptide **5** was tested for stability in human and mouse plasma. The pseudopeptide signal was present with minimally changed intensity after one hour and with decreased intensity after two hours incubation at 37°C (**Figure S16)**.

## Discussion

In this work, we developed the first low-molecular-weight PAPP-A inhibitors. Our strategy combined a peptide sequence derived from IGFBP-5, an endogenous PAPP-A substrate, with the replacement of the scissile bond by a pseudopeptide moiety. As a pseudopeptide bond, we employed several phosphinate and phosphonate variants. These non-hydrolysable isosteres were expected to mimic the tetrahedral intermediate formed during peptide bond hydrolysis and to bind strongly to the catalytic Zn²⁺ ion in the active site, thereby blocking substrate cleavage and inhibiting the enzyme.

Unlike other members of the metzincin family, such as matrixins, adamalysins and astacins, which are known for their low selectivity^39^, pappalysins exhibit very narrow substrate specificity. This specificity is largely governed by residues and structural constraints distant from the cleavage site^5,33,40^ while it has minimal restrictions on the amino acids directly adjacent to the scissile bond. Bearing this in mind, we selected the IGFBP-5 anchor peptide^26^ extended by eight C-terminal residues (peptide **1**) as the template for inhibitor design. The importance of the full-length anchor peptide sequence was demonstrated by the dramatic loss of affinity upon N-terminal truncation (**Table 1**), despite preservation of K^148^ (in pseudopeptide **6**), previously identified as a major contributor to binding^26,33^.

As expected, replacement of the scissile bond with a phosphinate markedly increased affinity, resulting in a 16-fold improvement for pseudopeptide **3** compared with peptide **1**. The C-terminal extension also increased the affinity of peptide **1** (IGFBP-5^141–171^, *K*_i_ = 144 nM) compared to the original anchor peptide (IGFBP-5^141-163^, *K*_d_ = 380 nM) (26). The pseudopeptide **3** demonstrated stereoselectivity with regards to the P1 residue norleucine (equivalent to IGFBP5 S^163^) and P1’ lysine, with only pseudo (*R*)-norleucine and pseudo (*R*)-lysine side chains observed in the cryo-EM structure despite the sample being prepared as an unresolved mixture of four diastereoisomers (**Figure 4B**). The requirement for (*R*)-amino acid in the P1’ is due to the pseudopeptide phosphinate having a different geometry to an amide of natural peptide bond where both (*S*, *L*)-lysine and (*S*, *L*)-norleucine would fit to the same pocket (**Figure S17**). Pseudo (*R*)-lysine occupies the S1’ pocket, which varies among the metzincin family members and has been the main selectivity determinant targeted for active-site directed inhibitors design^39^. An (*S*)-lysine of pseudopeptide side chain would sterically clash with the catalytic E^483^ and V^446^ in the observed pose, which permitted favorable backbone hydrogen bonds between A^446^ and A^449^ in PAPP-A and the P1’ and P2 residues of **3** (pseudo (*R*)-lysine, Q^162^), explaining the 4-fold increase in affinity compared to **2**, which includes only pseudo (*S*)-lysine at the P1’ position. Pseudo (*S*)-norleucine would similarly clash with E^483^, explaining its absence in the structure. The S1’ pocket was observed to have space for a larger side chain than lysine and have access to solvent towards its C-terminus, which explained pseudopeptide **4**’s similar potency to **3** despite the significant difference in their pseudo-dipeptide blocks. The lower affinity of **5** can be attributed to the greater separation between the lysine side chain and the phosphinate moiety.

Altogether, these observations suggest that an even more potent pseudopeptide inhibitors could be synthesized using (*R*)-norleucine side chain at the P1 position, a larger (*R*)-isomeric residue at the P1’ position and optimization of the C-terminal residues to improve or add to the hydrophobic interactions of F^165^ and V^166^ while retaining the favorable backbone geometry of pseudopeptide **3**.

These findings lay the groundwork for the rational development of future inhibitors. However, at the same time, our results highlight an important limitation of PAPP-A inhibitors. High-affinity binding to PAPP-A requires a relatively long peptide sequence, making the discovery of substantially smaller, purely chemical inhibitors or peptidomimetics challenging. This requirement together with striking stereospecificity inevitably increases both, the complexity of inhibitor design and the synthetic effort needed to generate potent compounds.

We have not observed any activity of our inhibitors against two tested matrix metalloproteinases (**Figure S15**), which probably reflects the high target specificity conferred by the extended IGFBP-5-derived peptide sequence. While the phosphinate/phosphonate group is expected to coordinate the catalytic Zn²⁺ ion, the surrounding peptide residues likely favor recognition by PAPP-A over MMPs. We should stress that our structure was determined using active PAPP-A and allowed detailed view of the catalytic site while the previously determined structures used active-site inactivated PAPP-A mutant^11,26,32^. The apparent looseness of the S and S’ pockets surrounding the catalytic site poses a question about possible existence of other, more feasible substrates that would better accommodate to the cleavage site. Evolutional studies have shown PAPP-A in invertebrates where IGFBPs do not exist^41^ and suggested that PAPP-A might have functions outside the IGF axis.

The model inhibitor was relatively stable in plasma (**Figure S16**) and the tested inhibitors possessed no detectable cytotoxicity (**Figure S14**). They even did not decrease the viability of the A549 cell line, which expresses PAPP-A and might be expected to rely on it. However, the assay was performed under standard growth conditions with abundant growth factors in the medium. We presume that the function of PAPP-A becomes more apparent under stress conditions and is related to the inducibility of its expression, as suggested by zebrafish studies^36,42^ and documented by our observation of significant changes in PAPP-A and STCs mRNA expression under stress induced by cultivation in serum-free medium (**Figure S10**).

IGFBP-5 is also a substrate of PAPP-A2. The physiological role of PAPP-A2 remains relatively unclear. Clinical data suggest an important role in the regulation of circulating IGFs^43^ but *in vitro* characterization of PAPP-A2 activity is still limited. Cleavage of IGFBP-5 by PAPP-A2 has been reported to be markedly less efficient (approximately 38-fold lower) than by PAPP-A^34^. In a structural study, the interaction of IGFBP-5 with PAPP-A2 was suggested to be similar to the interaction with PAPP-A but the binding of the anchor peptide could not be directly captured. Instead, it was inferred using AlphaFold prediction. In our experiments, we observed measurable PAPP-A2 activity only at high enzyme concentrations (100 nM). Q-BP5 was cleaved by this enzyme (**Figure S8 A**) and we might assume that our inhibitors are likely capable of interacting with PAPP-A2. But, their affinity might be so low that the observed inhibition of PAPP-A2 was negligible at the concentrations tested (**Figure S8 C**).

ProMBP, STC1 and STC2 are natural protein inhibitors of PAPP-A^9,11,32^. Potent inhibitory antibodies were also developed^25,44^. However, all these protein inhibitors act by binding to an exosite critical for IGFBP-4 recognition. ProMBP and STC2 form covalent complexes through cystine bond. All these proteins only partially restrict access to the active site, as evidenced by the retained ability to cleave peptide substrates^11,32^ and antibodies inactivity towards IGFBP-5 cleavage^44^.

In contrast, our inhibitors directly target and block the active site of PAPP-A and represent a new class of PAPP-A inhibitors. Our most potent compounds exhibited low nanomolar *K*_i_ values and effectively suppressed IGF-1 signaling in cell-based assay to a degree comparable to STC1 (**Figures 2E and S9**). Importantly, these compounds function as potent inhibitors of Papp-aa-regulated IGF signaling and cell proliferation *in vivo*. Both pseudopeptide **3** and pseudopeptide **5** inhibited Papp-aa-regulated NaR cell proliferation in Tg(igfbp5a:GFP) fish and pseudopeptide **5** was able to rescue the *stc1a-/-*phenotype (**Figures 3** and **S11**). *In vivo*, pseudopeptide **3** was more potent than pseudopeptide **5** consistent with the protease activity and cell-based assay results. At high doses, these inhibitors completely abolished the cell proliferation induced by IGF signaling. These results are particularly encouraging because the compounds were effective in a whole-organism context, in which compound uptake, tissue penetration, and accessibility to target cells can all influence efficacies. Together, these findings provide strong evidence supporting the feasibility of targeting PAPP-A/Papp-aa *in vivo* through a substrate-mimetic active-site blockade mechanism.

Beyond providing new chemical tools for studying PAPP-A biology, our findings may have broader therapeutic implications. Previous attempts to suppress the IGF axis through direct inhibition of IGF-1R have achieved only limited clinical success. In contrast, PAPP-A acts as a local regulator of IGF bioavailability and is expressed in a tissue- and context-dependent manner^13,21^. Therefore, pharmacological inhibition of PAPP-A may represent a strategy to attenuate pathological IGF signaling selectively within disease-associated microenvironments while preserving systemic IGF functions. Although this concept remains to be validated in disease models, the inhibitors described here provide a starting point for exploring this therapeutic avenue.

In summary, we have developed novel and potent low-molecular active-site directed PAPP-A inhibitors. We believe that these inhibitors may be applicable for the treatment of conditions that would benefit from reduced IGF availability and IGF signaling.

## Methods

Detailed methodology can be found in Supplementary information available online.

### Chemical syntheses

The non-standard amino acids for synthesis of quenched fluorescent peptide Q-BP5 were synthesized in house (**Supporting information Scheme 1, Figure S1**). 7-Methoxycoumarin-4-acetic acid (MCA) and 2,4-dinitrophenyl (DNP) used for preparation of fluorogenic substrate FS-6 were purchased from Iris Biotech GmBH. Syntheses of pseudo-dipeptide precursors and all intermediate compounds are described in Supporting information (**Schemes 2-7, Figures S2, S3**). Peptides were synthesized by solid phase synthesis using standard procedures in an automated peptide synthesizer. Pseudo-dipeptide blocks and modified amino acids were coupled manually. All peptide-based compounds were purified by reversed-phase high-performance liquid chromatography (RP-HPLC) and their identity was confirmed by high-resolution mass spectrometry (HR-MS), **Figures S4-S6**. Detailed methods including analyses can be found in Supporting information.

### Recombinant PAPP-A preparation

Recombinant PAPP-A was expressed in HEK293T cells (ATCC; CRL-3216) and isolated from conditioned media using standard procedures as described in Supporting methods. WT PAPP-A in pcDNA3.1-C-(k)DYK vector (Genscript (cloneID: OHu20096) was used for transfection. The conditioned media was filtered and concentrated using 100 kDa MWCO concentrator columns (Millipore UFC9100). PAPP-A was purified by FLAG affinity chromatography (Genscript Anti-DYKDDDDK G1 Affinity resin) according to manufacturer’s standard procedure. The protein was aliquoted and stored in -80 °C for further use. Concentration of the PAPP-A dimer was estimated from absorbance at 280 nm with ε = 468170 cm^-1^ x M^-1^ based on amino acid sequence. The effectiveness of the transfection was checked by western Blotting, using monoclonal anti-FLAG antibody conjugated with HRP (Sigma-Aldrich, A8592).

### Quenched fluorescence assay

The inhibitory potency of the compounds was measured by quenched fluorescence assay as described in Supporting methods. We designed and synthesized the fluorogenic substrate Q-BP5 to measure activity of recombinant PAPP-A. The measurements with PAPP-A were performed in reaction buffer 50 mM Tris-HCl pH 8.0 with 1 mM CaCl_2_ and 0.01% Triton X-100. We tested also commercial PAPP-A2 (Sino Biological, cat.no. 10528-H08H).

We synthesized an established substrate FS-6 (Ref.^31^) to measure activity of commercial MMP-1 (Sino Biological, cat.no. 10532-H08H) and MMP-7 (a kind gift from Prof. Sagi, WIS, Israel). The measurements with activated MMP-1 and MMP-7 were performed in reaction buffer 100 mM Tris-HCl pH 7.5, 100 mM NaCl, 10 mM CaCl_2_ with 0.01% Triton and 1% DMSO.

The substrates and inhibitors were serially diluted using BRAVO pipetting station (Agilent Technologies, USA) in black 384-well low volume U-bottom plates (Corning, USA). Pipetting was performed either manually, or compounds were pre-spotted using an ECHO 650 liquid handler (Beckman Coulter, USA) and CERTUS FLEX liquid dispenser added enzyme and substrate mixes. Reactions were started by the enzyme addition. The fluorescence was measured using Tecan Spark plate reader (Tecan, Switzerland) with excitation at 310 nm and emission at 420 nm for Q-BP5 and 325 nm and 400 nm for FS-6, in kinetic mode for 1 h at room temperature.

All samples were run as triplicates. The kinetic slope data were analyzed using GraphPad Prism 8 (GraphPad, USA), applying Michaelis-Menten non-linear regression model for *K*_m_ measurement. IC_50_ values were assessed by non-linear fitting of log(inhibitor) *vs* response variable slope. To estimate the inhibition constant (*K*_i_), we recalculated the IC_50_ data using Cheng-Prussoff equation.

### Cryo-EM

Pseudopeptide **3** and PAPP-A (Sino Biological, cat.no. 17290-H44H) were reconstituted in DPBS (final conc. 0.5 mg/mL of PAPP-A), applied to UltrAuFoil grids, blotted and vitrified in liquid ethane using a VitroBot MkIV (Thermo Fisher Scientific). 13,693 movies were collected on a Titan Krios G4 equipped with a Gatan K3 camera and Quantum-GIF energy filter. The movies were processed in cryoSPARC v4.7.1, resulting in a 2.6 Å final map (GSFSC 0.143). The model was built starting from PDB: 7UFG, which was refined against our map in ChimeraX v1.10, omitting the pseudo-dipeptide block. The pseudo-dipeptide block was then built manually in ChimeraX, restraints were generated using eLBOW within phenix v2.1 and the pose refined in coot v0.9.8.6. Final refinement statistics were generated in phenix. Detailed description of the methodology can be found in Supporting information.

### Assessment of cleavage of IGFBP-4 and IGFBP-5 by PAPP-A using western blot

The IGFBP-4 (Sino Biological, 10967-H08H) and IGFBP-5 (Sino Biological, 10206-H08B) cleavage by PAPP-A was followed using Western blot as described in Supporting information. The cleavage reactions were performed in reaction buffer 50 mM Tris-HCl pH 8,0; 1 mM CaCl_2_ for 1 h at 37 °C in absence or presence of serially diluted inhibitors (0.04-300 μM). IGFBP-4 was pre-incubated with IGF-1 (ratio 1:3) and IGFBP-5 was used without IGF-1 addition. The IGFBP-4 and IGFBP-5 cleavage was visualized using western blot analysis with an anti-IGFBP-4 primary antibody (Abcam, cat. no. AB205581; dilution 1:1000) and anti-6 × His tag primary antibody (Invitrogen, cat. no. MA1-135; dilution 1:1000) for IGFBP-5.

### RT-qPCR for mRNA expression along STC-PAPP-A-IGFBP-IGF axis

The PAPP-A, PAPP-A2, STC1/2, IGFBP-2/4/5 and IGF-1/2 mRNA expressions in the cell line A549 were assessed using standard RT-qPCR procedures as in Ref.^45^ and in Supporting methods. The expression of targeted genes was normalized by GAPDH mRNA levels. Used primers are listed in **Table S3**.

### Inhibition of IGF-1 signaling in a cell-based assay

We used A549 (ATCC; CCL-185) cell line to assess the ability of inhibitors to attenuate IGF-1 signaling through inhibition of its liberation from IGFBPs as described in Supporting methods. We showed PAPP-A expression in the A549 cell lysates on protein level using western blot using anti-PAPP-A antibody (Abcam, cat. no.: EPR12897) and anti-GAPDH antibody (Cell Signaling, cat. no.: 97166S) used as a loading control. Confluent cells were incubated overnight in serum-free medium. The cells were then pre-treated with the inhibitors for 1 h (pseudopeptides **3** and **5** (1 nM – 1 μM) or STC1 (0.1 – 10 nM) (Sino Biological, 15411-H08H)). IGF-1 (10 nM) was preincubated with IGFBP-4 or IGFBP-5 for 30 min in a ratio 1:2 and added to the cells for 20 min or 2 hours. The samples were analyzed using standard western blotting. Autophosphorylation of the receptor was detected using phospho-IGF-1Rβ (Tyr1135/1136)/IRβ (Tyr1150/1151) (19H7), Cell Signaling, and phosphorylation of Akt using phospho-Akt (Thr308) (C31E5E), Cell Signaling. Anti-actin^20–33^, Sigma-Aldrich, was used as a loading control.

### Inhibition of Papp-aa-Igfbp5a-IGF signaling *in vivo* in zebrafish

All zebrafish experiments were conducted in accordance with protocols approved by the University of Michigan Institutional Animal Care and Use Committee. Zebrafish were raised, maintained, crossed, and staged according to standard zebrafish husbandry guidelines. Embryos and larvae were maintained at approximately 28°C in standard E3 embryo medium.

The Tg(igfbp5a:GFP) line and *stc1a*(+17)⁺/⁻;Tg(igfbp5a:GFP) fish were generated in previous studies^35,38^. Fish embryos were raised in standard E3 medium until 3 days post-fertilization (dpf), transferred to 6-well plates, and washed three times with either control medium containing 0.2 mM Ca²⁺ or induction medium containing 0.001 mM Ca²⁺ before drug treatment. Pseudopeptides **3** and **5** were dissolved directly in induction medium. BMS-754807, purchased from MedChemExpress, Monmouth Junction, NJ, USA, was first dissolved in DMSO and then diluted in induction medium. Drug solutions were changed daily. After 2 days of drug treatment, fish were imaged, and NaR cells in the yolk-sac region were quantified as previously described^35^. Individual fish were subsequently genotyped as previously reported^38^.

### Statistical analysis

Statistical tests were carried out using GraphPad Prism 8 software (GraphPad Software, Inc., San Diego, CA). Values are shown as means ± SD. Statistical significance between experimental groups was performed using unpaired two-tailed t test, Chi-square test, long-rank test and one-way ANOVA followed by Tukey’s multiple comparison test. Statistical significances were accepted at *P* < 0.05 or greater.

### Cytotoxicity assay

Pseudopeptides **3**, **5**, **6**, **7**, **8** and **9** were tested for cytotoxicity by standard procedure employing CellTiter-Glo® 2.0 assay (Promega, Madison, USA) as described in Supporting methods. Four cancer cell lines (CCRF-CEM, HepG2, Hela S3, HL-60) and non-tumor human dermal fibroblasts (NHDF) were used. The A549 cell line was tested with pseudopeptides **3**, **5** and **8**. Compounds were spotted to 384-well white plates (Thermo Fisher Scientific, Waltham, USA) by ECHO550 Acoustic Liquid Handler (Beckman). Luminescence was measured by Tecan Spark plate reader (Tecan, Switzerland). The signal of the compound-treated cells (after 72 hours) was related to the value of untreated control which was arbitrarily set to 100% viability. IC_50_ values were calculated from dose-response curves using non-linear regression method using GraphPad Prism 8 software.

### Stability in plasma

Pseudopeptide **5** was incubated in human (Biowest) or mouse (Innovative Research) plasma for 0, 15, 60 and 120 min at 37°C. The proteins were precipitated with 90% acetonitrile, 0.1% TFA, 1 : 1 (v/v) for 1 hour and separated by centrifugation. Compounds contained in supernatant were after lyophilization resuspended in deionized water and analyzed using MALDI-TOF. The MALDI-TOF spectra were measured on rapifleX MALDI-TOF/TOF mass spectrometer (Bruker Daltonics, Germany) with Smartbeam 3D™ laser with up to 10 kHz repetitions. Internal standard (Mw 3765) was added to enable quantitation. The experiment was repeated twice. To monitor plasma enzymatic activity, propantheline (Sigma Aldrich) was used as a positive control. Detailed description can be found in Supporting information.

## Data availability

The data underlying this study are available in the article and its Supporting Information. Atomic model and map from cryo-EM have been deposited at the PDB and EMDB with the accession codes 45DF and EMD-83190, respectively. Raw data including all measurements, uncropped blots and AlphaFold3 computations are available *via* the Open Science Framework repository: 10.17605/OSF.IO/7CZ2B.

## Author Contributions

Conceptualization: I.S. and J.J. Methodology: A.K., S.-W.S., N.S.K., J.P., A.R., J.K., K.M., M.L., E.T. and M.B., Investigation: A.K., S.-W.S., N.S.K. K.M., M.B., C.D., J.J. and I.S. Formal analysis: A.K., S.-W.S., N.S.K., M.B., C.D. and I.S. Funding acquisition: J.J., N.S.K., and C.D. Supervision: I.S., J.J. and C.D. Writing—original draft and Writing—review and editing: I.S., J.J. N.S.K. and C.D.

## Additional information

The online version contains supplementary material available at.

Correspondence and requests for materials should be addressed to Irena Selicharová, Institute of Organic Chemistry and Biochemistry, Czech Academy of Sciences, Flemingovo nám. 2, Prague 6, 16610, Czech Republic, Phone: +420 220 183 236.

## Acknowledgements

We acknowledge the Biological pharmacology and Mass spectrometry service groups at IOCB.

We acknowledge use of the electron microscopy facilities at the Ian Holmes Imaging Centre, Bio21 Institute, Parkville, Australia.

## References

1. Chrudinová, M. et al., Evolution of insulin, insulin-like growth ffactors, and their cognate receptors in vertebrates, invertebrates, and viruses. Mol. Biol. Evol. 43, msaf319 (2026).

2. Choi, E., Duan, C. & Bai, X. C. Regulation and function of insulin and insulin-like growth factor receptor signalling. Nat. Rev. Mol. Cell. Biol. 26, 558–580 (2025).

3. Forbes, B. E., McCarthy, P., & Norton, R. S. Insulin-like growth factor binding proteins: a structural perspective. Front. Endocrinol. 3, 38 (2012).

4. Firth, S. M., & Baxter, R. C. Cellular actions of the insulin-like growth factor binding proteins. Endocr. Rev. 23, 824–854 (2002).

5. Boldt, H. B. et al., Mutational analysis of the proteolytic domain of pregnancy-associated plasma protein-A (PAPP-A): classification as a metzincin. Biochem. J. 358, 359–367 (2001).

6. Overgaard, M. T. et al., Pregnancy-associated plasma protein-A2 (PAPP-A2), a novel insulin-like growth factor-binding protein-5 proteinase. J. Biol. Chem. 276, 21849–21853 (2001).

7. Oxvig, C., Conover, C. A. The stanniocalcin-PAPP-A-IGFBP-IGF axis. J. Clin. Endocrinol. Metab. 108, 1624–1633 (2023).

8. Jepsen, M. R. et al., Stanniocalcin-2 inhibits mammalian growth by proteolytic inhibition of the insulin-like growth factor axis. J. Biol. Chem. 290, 3430–3439 (2015).

9. Kløverpris, S. et al., Stanniocalcin-1 potently inhibits the proteolytic activity of the metalloproteinase pregnancy-associated plasma protein-A. J. Biol. Chem. 290, 21915–21924 (2015).

10. Overgaard, M. T. et al., Expression of recombinant human pregnancy-associated plasma protein-A and identification of the proform of eosinophil major basic protein as its physiological inhibitor. J. Biol. Chem. 275, 31128–31133 (2000).

11. Zhong, Q. et al., Structural insights into the covalent regulation of PAPP-A activity by proMBP and STC2. Cell Discov. 8, 137 (2022).

12. Pillai, R. N., Konje, J. C., Tincello, D. G. & N. Potdar, Role of serum biomarkers in the prediction of outcome in women with threatened miscarriage: a systematic review and diagnostic accuracy meta-analysis. Hum. Reprod. Update 22, 228–239 (2016).

13. Conover, C. A. & Oxvig, C. The pregnancy-associated plasma protein-A (PAPP-A) story. Endocr. Rev. 44, 1012–1028 (2023).

14. Bale, L. K., West, S. A. & Conover, C. A. Inducible knockdown of pregnancy-associated plasma protein-A gene expression in adult female mice extends life span. Aging Cell 16, 895–897 (2017).

15. Conover, C. A. & Oxvig, C. PAPP-A: a promising therapeutic target for healthy longevity. Aging Cell 16, 205–209 (2017).

16. Conover, C. A. & Oxvig, C. PAPP-A and cancer. J. Mol. Endocrinol. 61, T1–T10 (2018).

17. Conover, C. A. Pregnancy-associated plasma protein-A (PAPP-A) and cardiovascular disease. Growth Horm. IGF Res. 79, 101625 (2024).

18. Kashyap, S. et al., Metalloproteinase PAPP-A regulation of IGF-1 contributes to polycystic kidney disease pathogenesis. JCI Insight 5, e135700 (2020).

19. Jepsen, M. R. et al., Increased activity of the metalloproteinase PAPP-A promotes diabetes-induced glomerular hypertrophy. Metabolism 132, 155218 (2022).

20. Bale, L. K. et al., Genetic and pharmacological inhibition of PAPP-A reduces bleomycin-induced pulmonary fibrosis in aged mice via reduced IGF signaling. Aging Biol. 2, e20240023 (2024).

21. Oxvig, C. The role of PAPP-A in the IGF system: location, location, location. J. Cell Commun. Signal. 9, 177–187 (2015).

22. Wang, P., Mak, V. C. Y., & Cheung, L. W. T. Drugging IGF-1R in cancer: New insights and emerging opportunities. Genes Dis. 10, 199–211 (2023).

23. Baserga, R. The decline and fall of the IGF-I receptor. J. Cell Physiol. 228, 675–679 (2013).

24. Frystyk, J., Teran, E., Gude, M. F., Bjerre, M. & Hjortebjerg, R. Pregnancy-associated plasma proteins and stanniocalcin-2 – novel players controlling IGF-I physiology. Growth Horm. IGF Res. 53-54, 101330 (2020).

25. Mikkelsen, J. H. et al., Indirect targeting of IGF receptor signaling in vivo by substrate-selective inhibition of PAPP-A proteolytic activity. Oncotarget 5, 1014–1025 (2014).

26. Judge, R. A. et al., Structure of the PAPP-A(BP5) complex reveals mechanism of substrate recognition. Nat. Commun. 13, 5500 (2022).

27. Yiallouros, I. et al., Phosphinic peptides, the first potent inhibitors of astacin, behave as extremely slow-binding inhibitors. *Biochem*, J, 331 **( Pt** **2**), 375–379 (1998).

28. Dive, V. et al., Phosphinic peptides as zinc metalloproteinase inhibitors. Cell Mol. Life Sci. 61, 2010–2019 (2004).

29. Collinsová, M. & Jiracek, J. Phosphinic acid compounds in biochemistry, biology and medicine. Curr Med. Chem. 7, 629–647 (2000).

30. Gyrup, C., Christiansen, M. & Oxvig, C. Quantification of proteolytically active pregnancy-associated plasma protein-A with an assay based on quenched fluorescence. Clin. Chem. 53, 947–954 (2007).

31. Neumann, U., Kubota, H., Frei, K., Ganu, V. & Leppert, D. Characterization of Mca-Lys-Pro-Leu-Gly-Leu-Dpa-Ala-Arg-NH2, a fluorogenic substrate with increased specificity constants for collagenases and tumor necrosis factor converting enzyme. Anal. Biochem. 328, 166–173 (2004).

32. Kobberø, S. D. et al., Structure of the proteolytic enzyme PAPP-A with the endogenous inhibitor stanniocalcin-2 reveals its inhibitory mechanism. Nat. Commun. 13, 6084 (2022).

33. Laursen, L. S. et al., Substrate specificity of the metalloproteinase pregnancy-associated plasma protein-A (PAPP-A) assessed by mutagenesis and analysis of synthetic peptides: substrate residues distant from the scissile bond are critical for proteolysis. Biochem. J. 367, 31–40 (2002).

34. Sridar, J. et al., Cryo-EM structure of human PAPP-A2 and mechanism of substrate recognition. Commun. Chem. 6, 234 (2023).

35. Liu, C. et al., Development of a whole organism platform for phenotype-based analysis of IGF1R-PI3K-Akt-Tor Action. Sci. Rep. 7, 1994 (2017).

36. Liu, C. et al., The metalloproteinase Papp-aa controls epithelial cell quiescence-proliferation transition. eLife 9 (2020).

37. Liu, C. et al., Ca(2+) concentration-dependent premature death of igfbp5a(-/-) fish reveals a critical role of IGF signaling in adaptive epithelial growth. Sci. Signal. 11 (2018).

38. Li, S. et al., Calcium state-dependent regulation of epithelial cell quiescence by stanniocalcin 1a. Front. Cell Dev. Biol. 9, 662915 (2021).

39. Georgiadis, D. & Yiotakis, A. Specific targeting of metzincin family members with small-molecule inhibitors: progress toward a multifarious challenge. Bioorg. Med. Chem. 16, 8781–8794 (2008).

40. Weyer, K. et al., A substrate specificity-determining unit of three Lin12-Notch repeat modules is formed in trans within the pappalysin-1 dimer and requires a sequence stretch C-terminal to the third module. J. Biol. Chem. 282, 10988–10999 (2007).

41. Kjeldsen, C. M. N. & C. Oxvig, The proteinase PAPP-A has deep evolutionary roots outside of the IGF System. Genome Biol. Evol. 17, evaf042 (2025).

42. Li, S. Li, H. Wang, Z. & Duan, C. Stanniocalcin 1a regulates organismal calcium balance and survival by suppressing Trpv6 expression and inhibiting IGF signaling in zebrafish. Front. Endocrinol. 14, 1276348 (2023).

43. Nimptsch, K. et al., Pregnancy associated plasma protein-A2 (PAPP-A2) and stanniocalcin-2 (STC2) but not PAPP-A are associated with circulating total IGF-1 in a human adult population. Sci. Rep. 14, 1770 (2024).

44. Mohrin, M., et al., Inhibition of longevity regulator PAPP-A modulates tissue homeostasis via restraint of mesenchymal stromal cells. Aging Cell 20, e13313 (2021).

45. Dzianová, P. et al., The efficiency of insulin production and its content in insulin-expressing model β-cells correlate with their Zn(2+) levels. Open Biol. 10, 200137 (2020).

46. He, J., Li, T., & Huang, S.-Y. Improvement of cryo-EM maps by simultaneous local and non-local deep learning. Nat. Commun. 14, 3217 (2023).

