## Supplementry Information for "Synthetic inhibitors of pregnancy-associated plasma protein-A regulate IGF-1 signaling *in vitro* and *in vivo*"


**This PDF file includes:**

Schemes 1 to 7
Figures S1 to S17
Tables S1 to S3
Supporting text
SI References

**Other supporting materials for this manuscript include the following:**

Raw data are available via the Open Science Framework repository:
10.17605/OSF.IO/7CZ2B

**Content:** **Page**


**Synthetic schemes**

List of Abbreviations 4
Scheme 1: Synthesis of protected nonstandard amino acids (**18** and **20**) 5
Scheme 2: Synthesis of derivatized phosphinic acid (**28**) 6
Scheme 3: Synthesis of pseudo-dipeptide precursor **I** 7
Scheme 4: Synthesis of pseudo-dipeptide precursor **II** 8
Scheme 5: Synthesis of pseudo-dipeptide precursor **III** 9
Scheme 6: Synthesis of precursor **IV** 10
Scheme 7: Synthesis of pseudo-dipeptide **9** 11

**Figures**

Figure S1: Analytical RP-HPLC of nonstandard amino acids 12
Figure S2: Analytical RP-HPLC of key synthetic intermediates 13
Figure S3: Analytical RP-HPLC of pseudo-dipeptide precursors **I-IV** 14
Figure S4: HR-MS and analytical RP-HPLC of fluorescent substrates 15
Figure S5: HR-MS and analytical RP-HPLC of purified peptide **1**, pseudopeptides **2-4** and pseudo-dipeptide **9** 16
Figure S6: HR-MS and analytical RP-HPLC of purified pseudopeptide **5** and its shortened versions **6-8** 18
Figure S7: PAPP-A purification and inhibition of activity 20
Figure S8: PAPP-A2 activity determination 21
Figure S9: Inhibition of PAPP-A activity with pseudopeptide **5** and **3** 22
Figure S10: Expression of STC-PAPP-A-IGFBP axis genes 23
Figure S11: Inhibition of zebrafish Papp-aa activity *in vivo* 24
Figure S12: Cryo-EM processing workflow 25
Figure S13: Supplementary modelling 26
Figure S14: Cytotoxicity 27
Figure S15: Inhibition of MMP-1 and MMP-7 28
Figure S16: Stability in plasma 30
Figure S17: AlphaFold 3 modeled active site of native peptide 31

**Tables**

Table S1: Averaged RMSD for trajectories in MD simulation 32
Table S2: Refinement statistics for cryo-EM structure 33
Table S3: Primers for qRT-PCR 34

**Supporting Methods**

1. Chemical syntheses 35
1.1: Synthesis of modified amino acids 35
1.1.1: Synthetic protocols for compounds in Scheme 1 36
1.2: Synthesis of pseudo-dipeptide precursors **I-IV** 42
1.2.1: Synthesis of Cbz-Nleψ[PO(OH)H] 42
1.2.1.1: Synthetic protocols for compounds in Scheme 2 43
1.2.2.: Synthesis of precursor **I** 47
1.2.2.1: Synthetic protocols for compounds in Scheme 3 47
1.2.3: Synthesis of precursor **II** 52
1.2.3.1: Synthetic protocols for compounds in Scheme 4 53
1.2.4: Synthesis of precursor **III** 59
1.2.4.1: Synthetic protocols for compounds in Scheme 5 59
1.2.5: Synthesis of precursor **IV** 63

1.2.5.1: Synthetic protocols for compounds in Scheme 6 64
1.3: Synthesis of peptides and pseudopeptides 67
1.3.1: Fluorescent substrates Q-BP5 and FS-6, peptide **1** and pseudopeptides **2-8** 67
1.3.2: Pseudopeptide **9** 68
1.3.2.1: Synthetic protocols for compounds in Scheme 7 68
2. Cell culture 71
3. Recombinant PAPP-A preparation 71
4. Quenched fluorescence assay 72
5. Assessment of cleavage of IGFBP-4 and IGFBP-5 by PAPP-A using Western Blot 73
6. RT-qPCR for mRNA expression along STC-PAPP-A-IGFBP-IGF axis 74
7. Inhibition of IGF-1 signaling in a cell-based assay 74
8. Inhibition of IGF signaling in zebrafish *in vivo* 75
9. Cytotoxicity assay 75
10. Stability in plasma 76
11. Molecular modeling 77
12. Cryo-EM 79
12.1: Cryo-EM sample preparation and data collection 79
12.2: Cryo-EM refinement 79
12.3: Model building 80

SI References 80

**Synthetic schemes**

**List of Abbreviations**

| | |
|---|---|
| Ad | Adamantan-1-yl |
| Bn | Benzyl |
| Boc | *tert*-butoxycarbonyl |
| $Boc_2O$ | di-*tert*.-butyldicarbonate |
| Boc-GABA-OH | 4-(*tert*-butoxycarbonylamino)butanoic acid |
| BOP | (Benzotriazol-1-yloxy)tris(dimethylamino)phosphonium hexafluorophosphate |
| Cbz | Benzyloxycarbonyl |
| Cbz-OSu | *N*-(Benzyloxycarbonyloxy)succinimide |
| DCM | Dichloromethane |
| DIAD | Diisopropyl azodicarboxylate |
| DIC | *N*,*N*´-diisopropylcarbodiimide |
| DIU | *N*,*N*´-diisopropylurea |
| DMAP | 4-(dimethylamino)pyridine |
| DMF | Dimethylformamide |
| DPPA | Diphenylphophoryl azide |
| EDC.HCl | 1-ethyl-3-(3-dimethylaminopropyl)carbodiimide hydrochloride |
| Eschenmoser´s salt | *N*,*N*-Dimethylmethyleneamonium iodide |
| Fmoc | 9-Fluorenylmethoxycarbonyl |
| Fmoc-OSu | *N*-(9-fluorenylmethoxycarbonyloxy)succinimide |
| DIPEA | Diisopropylethylamin |
| HATU | *N*-[(dimethylamino)-3-oxo-1H-1, 2, 3-triazolo[4,5-b]pyridin-1-yl-methylene]-*N*-metylmethanaminium hexafluorophosphate |
| HBTU | *O*-(benzotriazol-1-yl)-*N,N,N´,N´*-tetramethyluronium hexafluorophosphate |
| HCTU | *O*-(1H-6-Chlorobenzotriazole-1-yl)-1, 1, 3, 3, -tetramethyluronium hexafluorophosphate |
| HMDS | Hexamethyldisilazane |
| HOBt | Hydroxybenzotriazole |
| Meldrum´s acid | 2,2-Dimethyl-1,3-dioxan-4,6-dione |
| NHS | *N*-hydroxysuccinimide |
| PyBroP | Bromotripyrrolidinophosphonium hexafluorophosphate |
| TEA | Triethylamine |
| THF | Tetrahydrofuran |
| TFA | Trifluoroacetic acid |
| TIS | Triisopropylsilane |

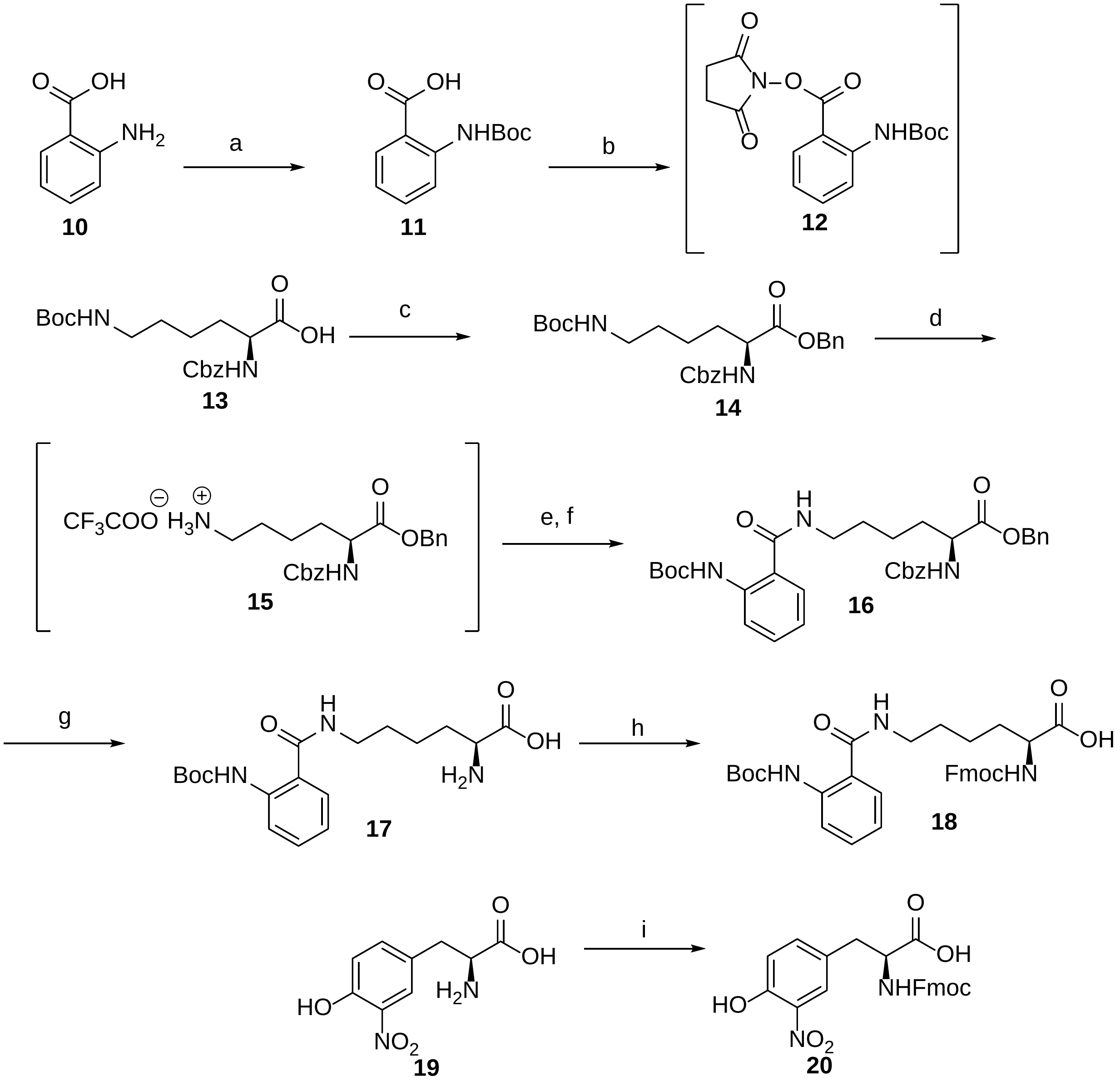


**Scheme 1. Synthesis of protected nonstandard amino acids 18 and 20**

Reagents, conditions, and yields: (a) $Boc_2O$, $NaHCO_3$, $H_2O$/dioxane, rt, 72 h, 70%; (b) NHS, DIC, THF, 0 °C for 1 h, then rt overnight; (c) BnBr, $K_2CO_3$, DMF, rt overnight, 84%; (d) TFA, DCM, 0 °C for 1 h, then rt for 1 h; (e) **12**, TEA, DCM, rt overnight, 89%; (f) **11**, PyBroP, DIPEA, DMF, rt overnight, 51%; (g) $H_2$, 10% Pd/C, MeOH, rt, 15 psi, 18 h, 80%; (h) Fmoc-OSu, $NaHCO_3$, $H_2O$/dioxane, 0 °C for 1 h, then rt overnight, 48%; (i) Fmoc-OSu, $NaHCO_3$, $H_2O$/dioxane, 0 °C for 1 h, then rt overnight, 93%.

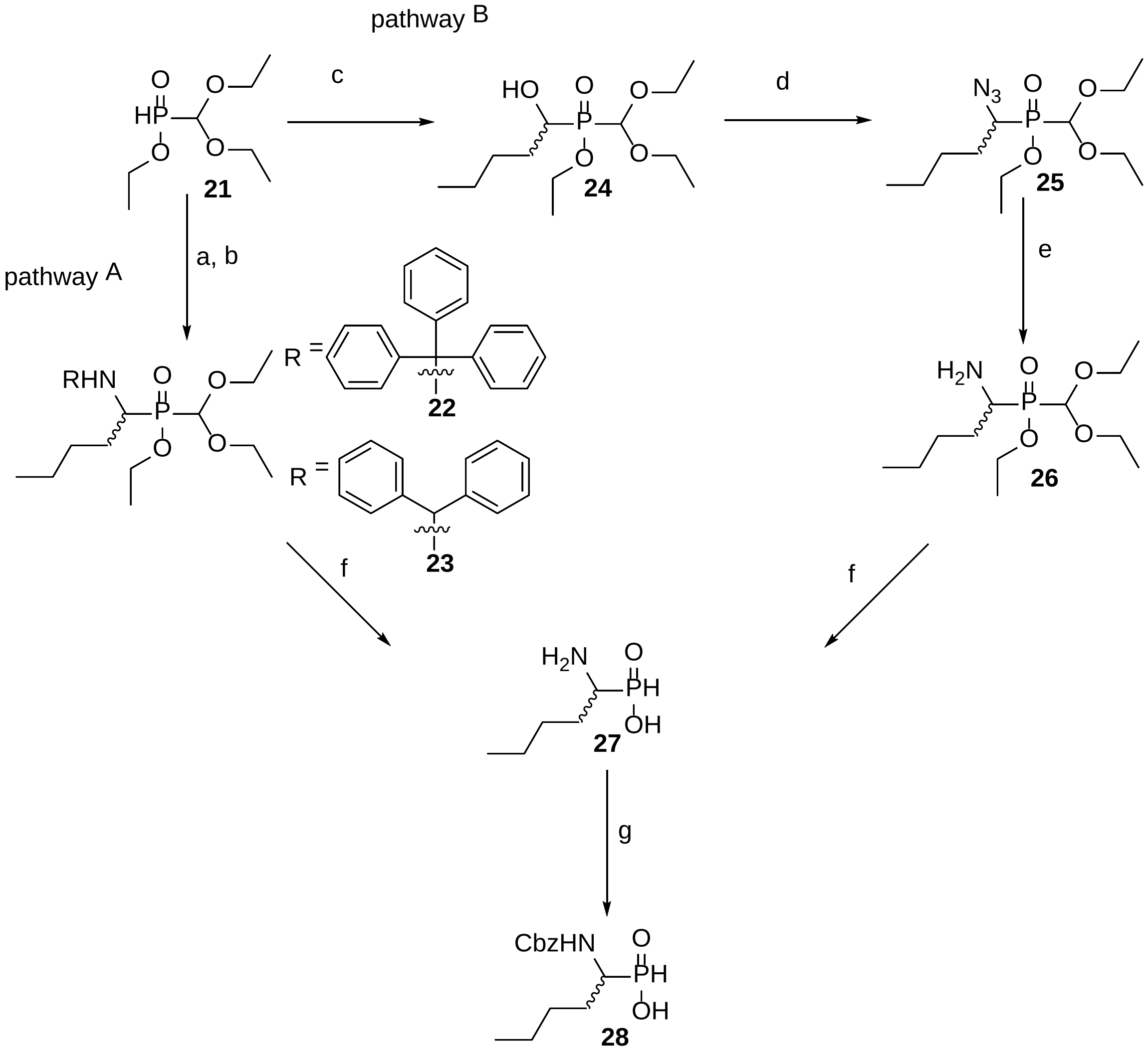


**Scheme 2**: **Synthesis of derivatized phosphinic acid 28**

Reagents, conditions, and yields: (a) $TrtNH_2$, valeraldehyde, benzene, reflux, 59%; (b) diphenylmethylamine, valeraldehyde, benzene, reflux, 69%; (c) valeraldehyde, DIPEA, 95 °C, 48 h, 60%; (d) $PPh_3$, DIAD, DPPA, THF, 0 °C for 1 h, then rt overnight, 87%; (e) $H_2$, 10% Pd/C, MeOH, rt, 15 psi, 12 h, 87%; (f) 35% HCl, reflux overnight, then 1,2-epoxypropane, EtOH, 77% (from 22), 71% (from 23), and 77% (from 26); (g) Cbz-OSu, $Na_2CO_3$, $H_2O$/dioxane, 0 °C for 1 h, then rt overnight, 82%.

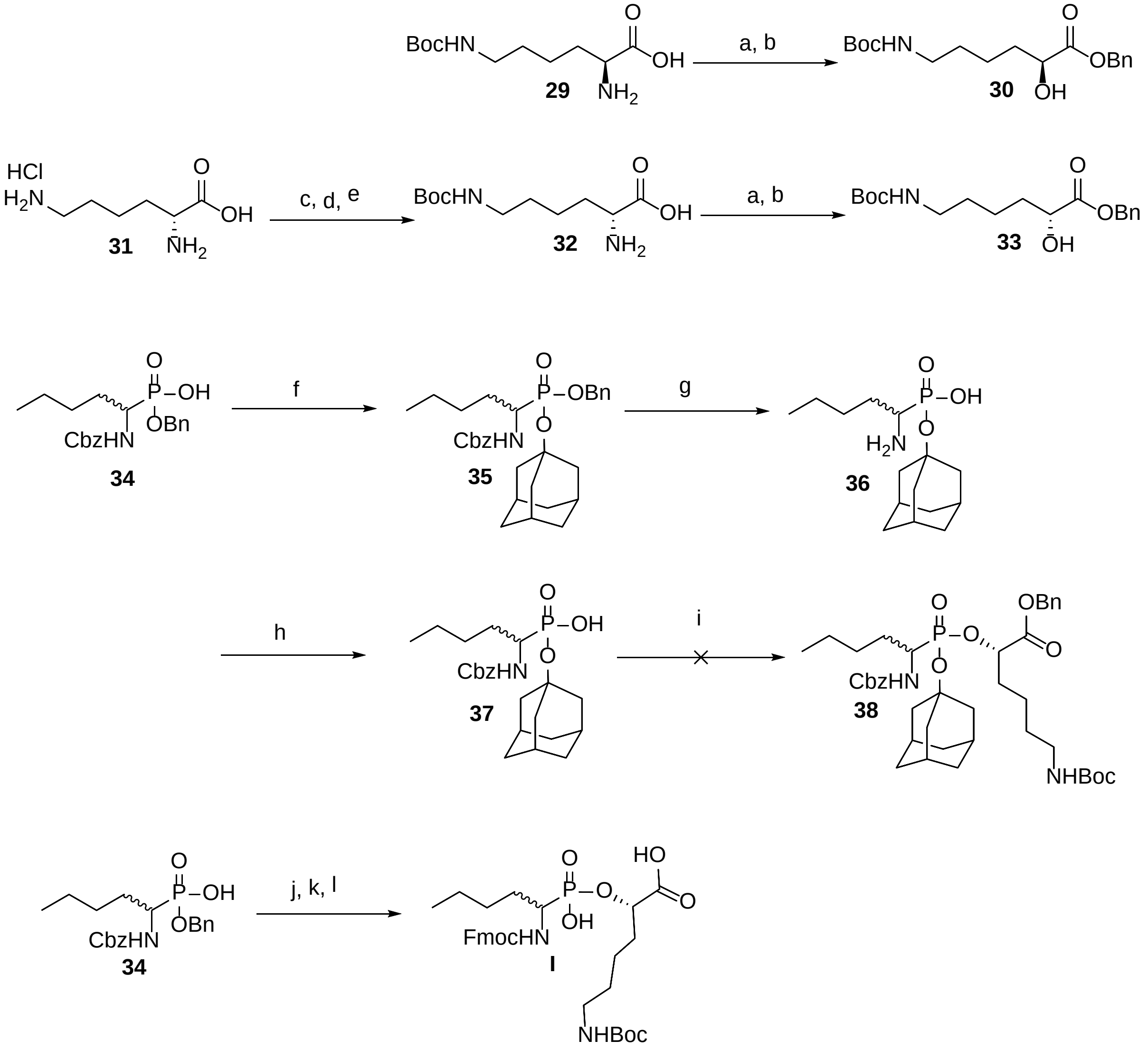


**Scheme 3: Synthesis of pseudo-dipeptide precursor I**

Reagents, conditions, and yields: (a) $NaNO_2$, $H_2O$, AcOH, 0 °C, 1 h, then rt, 4 h; (b) benzyl bromide, $K_2CO_3$, DMF, rt, overnight (35% over two steps for 30, 38% over two steps for 33); (c) $Cu(OAc)_2 \cdot H_2O$, NaOH, $H_2O$, rt, 1 h; (d) $Boc_2O$, acetone, overnight; (e) 8-hydroxyquinoline, $H_2O$, rt, 4 h (93% over three steps); (f) $Ag_2O$, 1-bromoadamantane, $CHCl_3$, reflux, overnight (93%); (g) $H_2$, 10% Pd/C, MeOH, rt, 15 psi, 18 h (80%); (h) Cbz-OSu, $Na_2CO_3$, $H_2O$, dioxane, 0 °C, 1 h, then rt, overnight (74%); (i) either 30, BOP, PyBroP, HCTU, HATU, DIPEA, DMF, rt, overnight, or 33, $PPh_3$, DIAD, DPPA, THF, rt, overnight (0%); (j) 33, $PPh_3$, DIAD, DPPA, THF, rt, overnight; (k) $H_2$, 10% Pd/C, MeOH, rt, 15 psi, 12 h; (l) Fmoc-OSu, $NaHCO_3$, $H_2O$, dioxane, 0 °C, 1 h, then rt, overnight (47% over three steps).

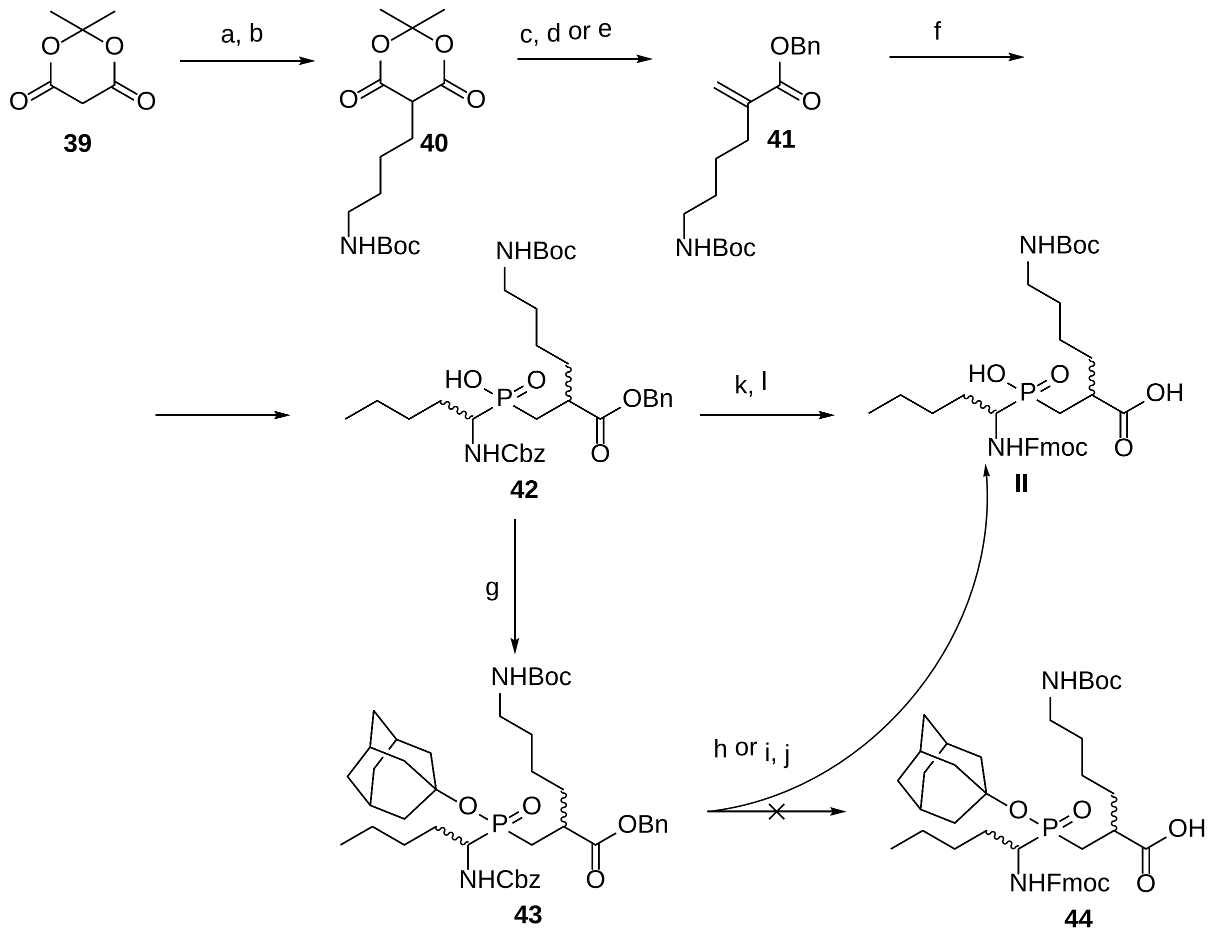


**Scheme 4: Synthesis of pseudo-dipeptide precursor II**

Reagents, conditions, and yields: (a) Boc-GABA-OH, DMAP, DIC, DCM, 0 °C for 1 h, then rt overnight; (b) $NaBH_4$, AcOH, DCM, 0 °C for 2 h, then rt overnight, 22% over two steps; (c) benzyl alcohol, toluene, 80 °C overnight; (d) paraformaldehyde, pyridine, piperidine, 80 °C overnight, 66% over two steps; (e) Eschenmoser's salt, benzyl alcohol, 65 °C overnight, 45%; (f) HMDS, 28, 110 °C for 3 h, 80%; (g) 1-bromoadamantane, $Ag_2O$, $CHCl_3$, 44%; (h) $H_2$, 10% Pd/C, MeOH, rt, 15 psi, 4 h; (i) $HCOONH_4$, 10% Pd/C, MeOH, rt, 12 min; (j) Fmoc-OSu, $NaHCO_3$, $H_2O$/dioxane, 0 °C for 1 h, then rt overnight, 42% or 38% over two steps; (k) $H_2$, 10% Pd/C, MeOH, rt, 15 psi, 8 h; (l) Fmoc-OSu, $NaHCO_3$, $H_2O$/dioxane, 0 °C for 1 h, then rt overnight, 55% over two steps.

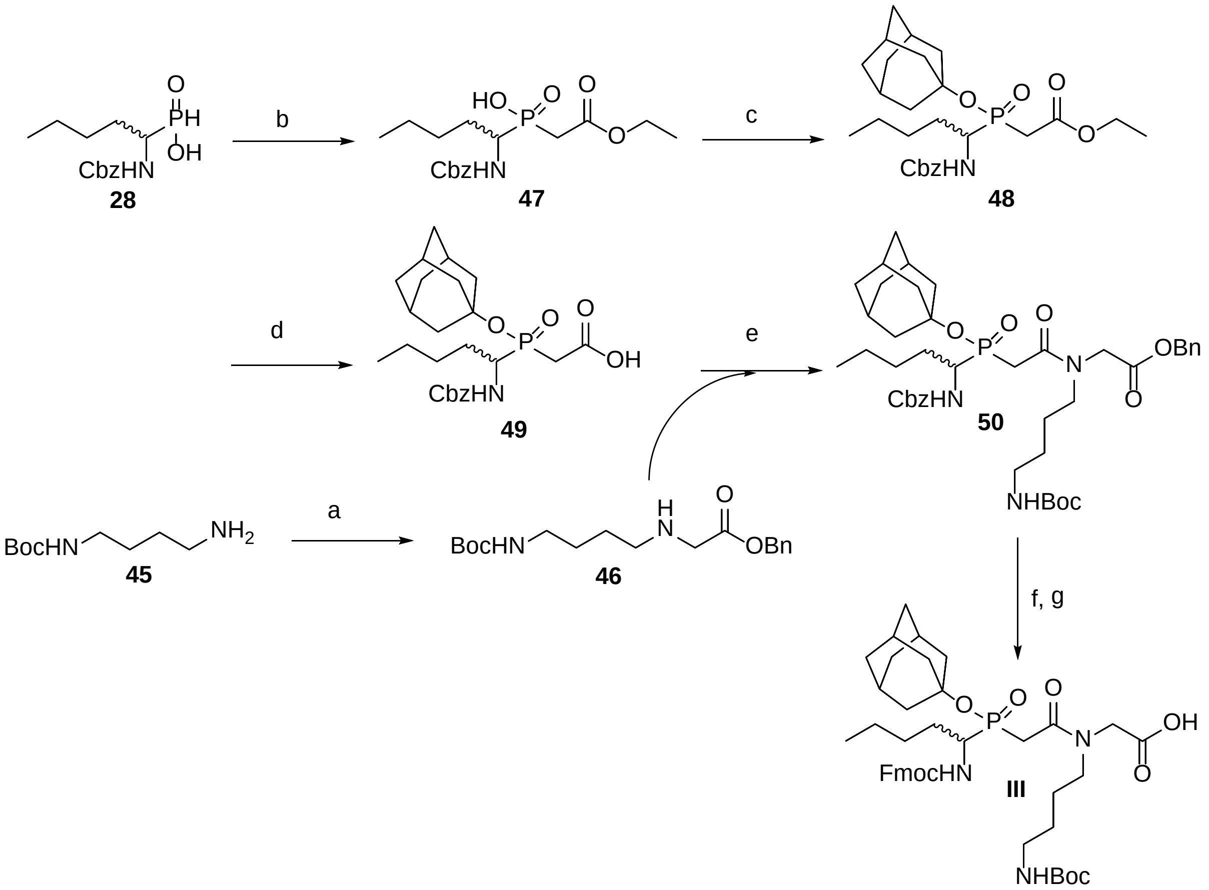


**Scheme 5: Synthesis of pseudo-dipeptide precursor III**

Reagents, conditions, and yields: (a) benzyl 2-bromoacetate, TEA, THF, rt overnight, 74%; (b) HMDS, ethyl 2-bromoacetate, 110 °C for 3 h, 80%; (c) $Ag_2O$, 1-bromoadamantane, $CHCl_3$, 60 °C overnight, 79%; (d) NaOH, MeOH/$H_2O$, rt overnight, 88%; (e) 46, DIC, DCM, 0 °C for 1 h, then rt overnight, 89%; (f) $H_2$, 10% Pd/C, MeOH, rt, 15 psi, 8 h; (g) Fmoc-OSu, $NaHCO_3$, $H_2O$/dioxane, 0 °C for 1 h, then rt overnight, 71% over two steps.

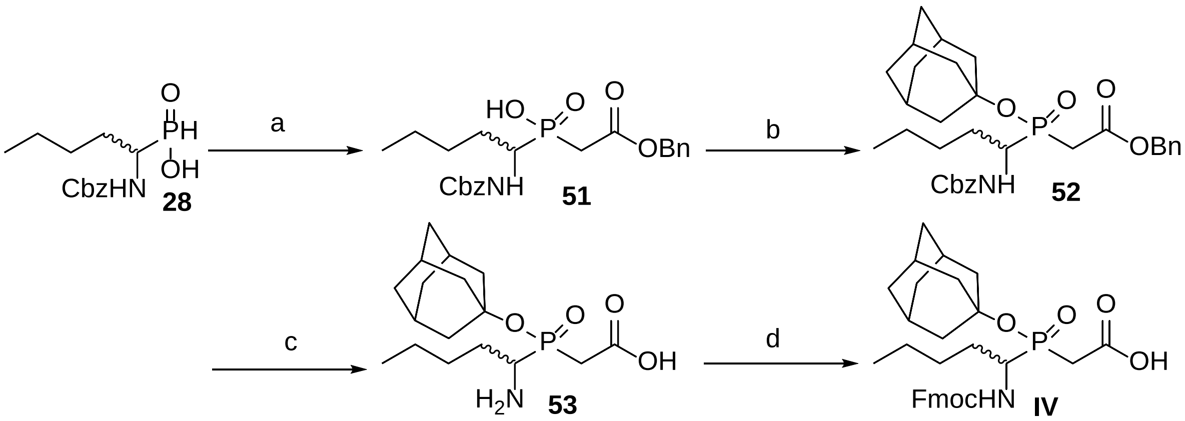


**Scheme 6: Synthesis of precursor IV**

Reagents, conditions, and yields: (a) HMDS, benzyl 2-bromoacetate, 110 °C for 3 h, 84%; (b) $Ag_2O$, 1-bromoadamantane, $CHCl_3$, 60 °C overnight, 77%; (c) $H_2$, 10% Pd/C, MeOH, rt, 15 psi, 12 h, 86%; (d) Fmoc-OSu, $Na_2CO_3$, $H_2O$/dioxane, 0 °C for 1 h, then rt overnight, 53%.

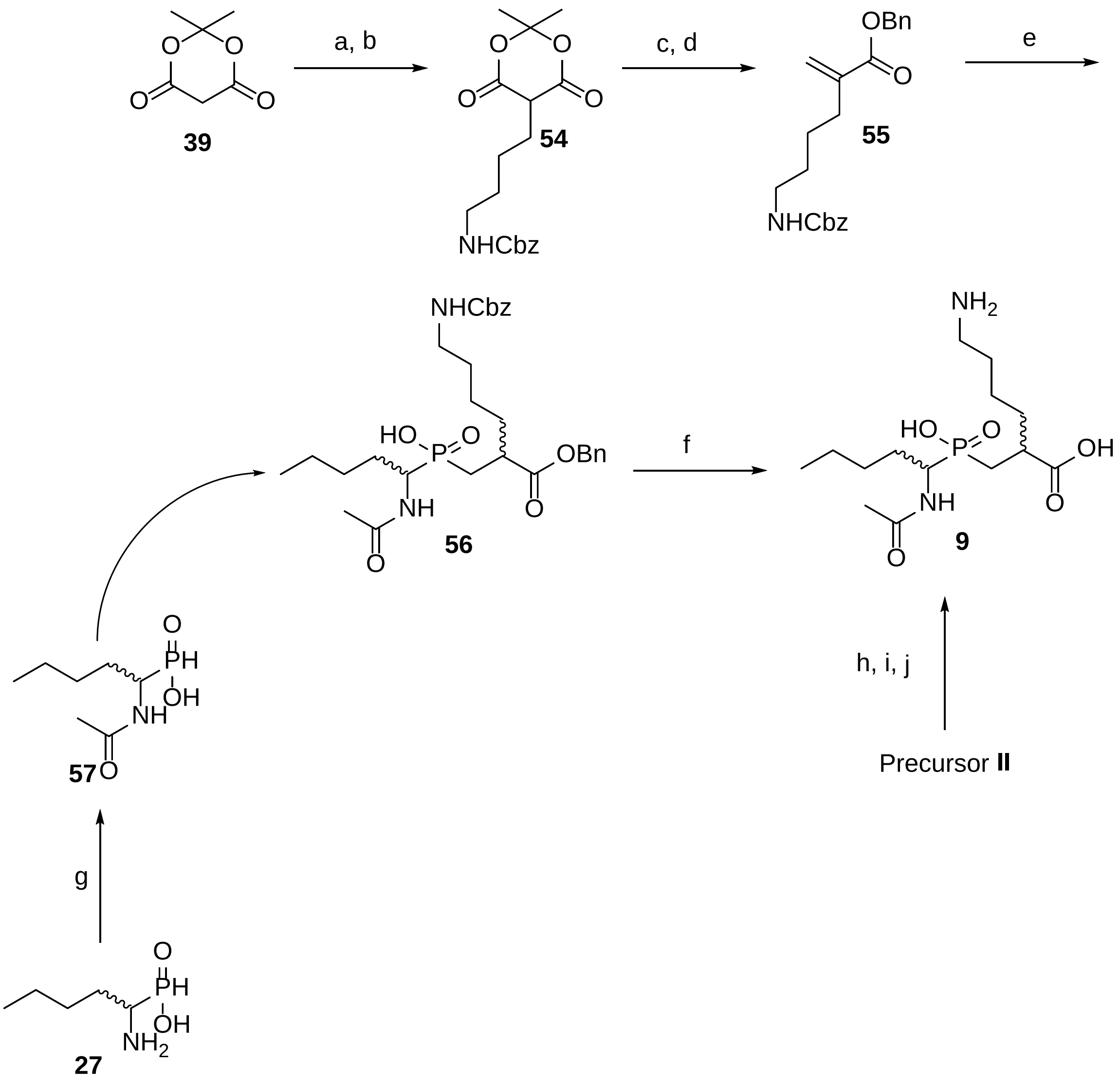


**Scheme 7: Synthesis of pseudodipeptide 9**

Reagents, conditions and yields: Cbz-GABA-OH, DMAP, DIC, DCM, 0°C for 1 hour then rt overnight; (b) $NaBH_4$, AcOH, DCM, 0°C for 2 hours then rt overnight (62% over two steps); (c) benzyl alcohol, toluene, 80°C overnight; (d) paraformaldehyde, pyridine, piperidine, 80°C overnight (71% over two steps); (e) HMDS, **57**, 110°C for 3 hours (72 %); (f) 10% Pd/C, $H_2$, 15 psi, methanol, rt for 18 hours (90%); (g) $Ac_2O$, $NaHCO_3$, water, dioxane, 1 hour at 0°C then rt overnight (51%); (h) **II**, 20% piperidine/DMF; (i) $Ac_2O$, DIPEA, DMF, (j) TFA (15% over three steps).

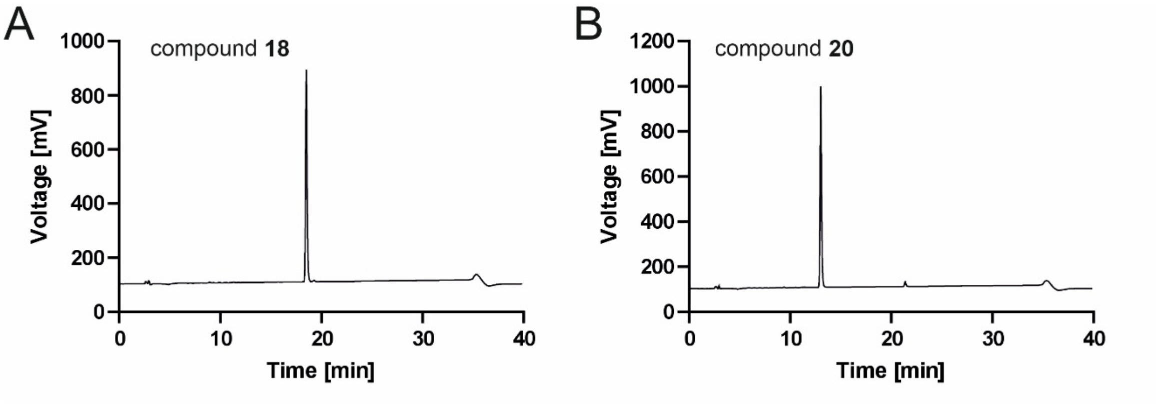


**Figure S1**: **Analytical RP-HPLC of nonstandard amino acids**
**A**) 2-(*S*)-[(9H-Fluoren-9-ylmethoxycarbonyl)amino]-6-[[2-(*tert*-butoxycarbonyl)amino] benzoyl-amino]hexanoate **18**. **B**) 2-(*S*)-[(9H-Fluoren-9-ylmethoxycarbonyl)amino]-3-(4-hydroxy-3-nitrophenyl)hexanoic acid **20**. Purified compounds were analyzed on a C8 column (EC 250-4.6 Nucleosil 100-5 C8, Macherey-Nagel) using solvents A (0.1% TFA v/v in water) and B (80% v/v acetonitrile in 0.1% v/v TFA in water) in gradient t= 0 min. (50% B in A), t = 30 min (100% B), 31 min (50% B in A) at a flow 1 ml/min.

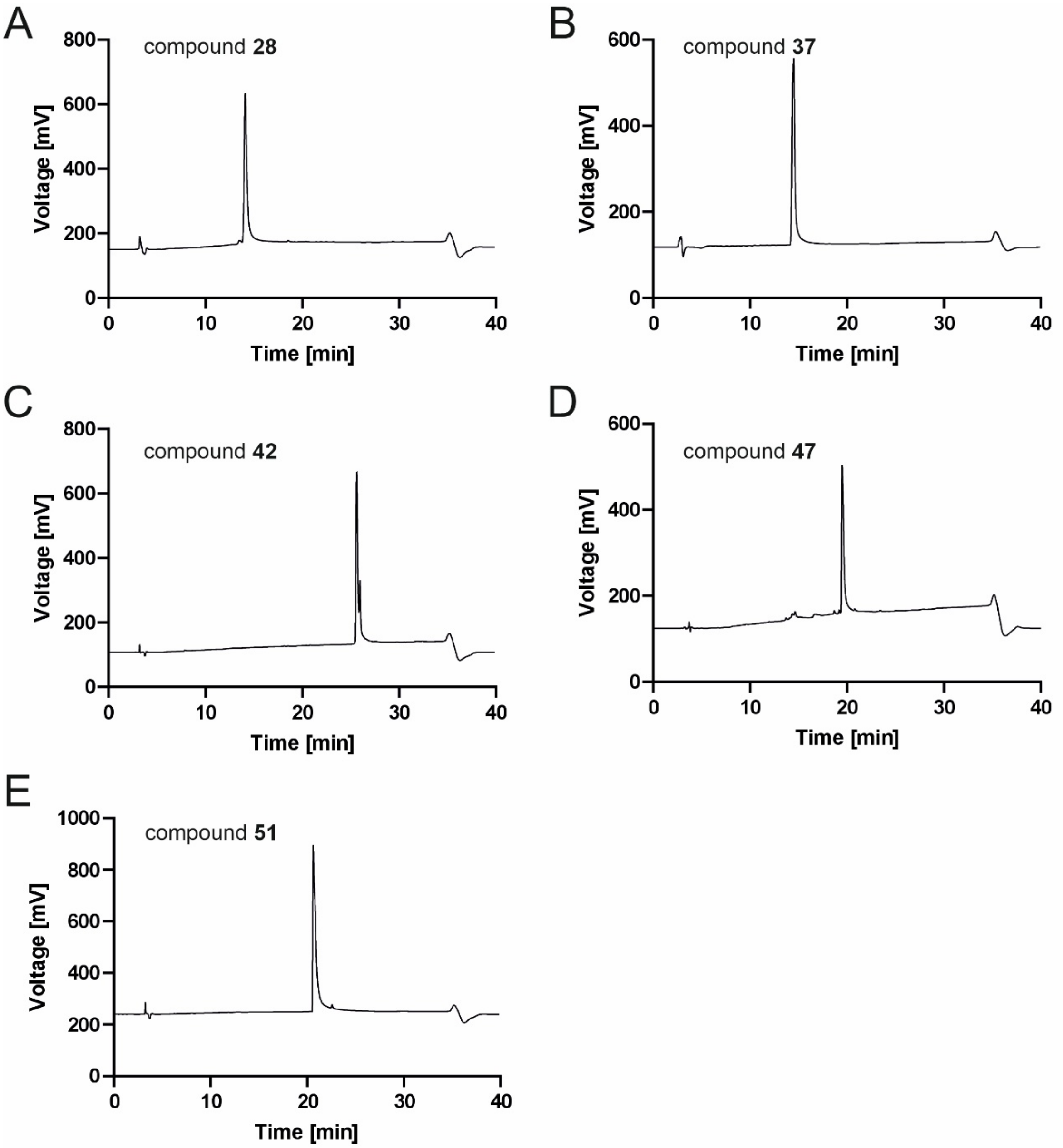


**Figure S2: Analytical RP-HPLC of key synthetic intermediates**

**A)** ((*R*,*S*) -1-{[(Benzyloxy)carbonyl]amino}pentyl)phosphinic acid **28**. **B**) Adamantan-1-yl hydrogen [(*R*,*S*)-1-(benzyloxycarbonylamino)-pentyl]phosphonate **37**. **C**) Benzyl (*R*,*S*)-2-[((*R*,*S*)-1-{[(benzyloxy)car-bonyl]amino}pentyl)(hydroxyphosphoryl)methyl]-6-[(tert-butoxycarbonyl)-amino]hexanoate **42**. **D**) Ethyl [((*R*,*S*)-1-{[(benzyloxy)carbonyl]amino}pentyl)(hydro-xy)phosphoryl]acetate **47**. **E**) Benzyl [((*R*,*S*)-1-{[(benzyloxy)carbonyl]amino}pentyl)(hydro-xy)phosphoryl]acetate **51**. Purified compounds were analyzed on a C8 column (EC 250-4.6 Nucleosil 100-5 C8, Macherey-Nagel) using solvent A (0.1% TFA v/v in water) and solvent B (80% v/v acetonitrile in 0.1% v/v TFA in water). Compounds **28**, **42**, and **51** were analyzed using the following gradient: t = 0 min, 20% B; t = 30 min, 100% B; t = 31 min, 20% B. Compound **47** was analyzed using the gradient: t = 0 min, 10% B; t = 30 min, 100% B; t = 31 min, 10% B. The flow rate was 1 mL/min.

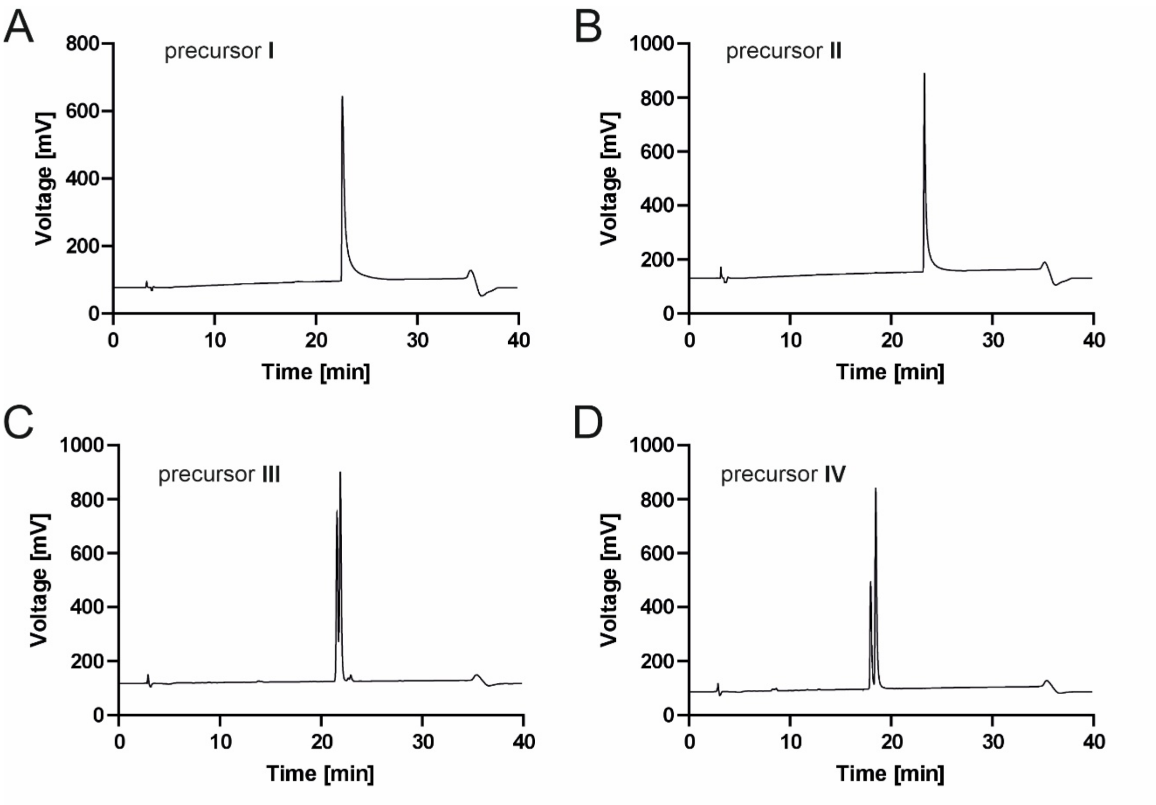


**Figure S3: Analytical RP-HPLC of pseudo-dipeptide precursors I-IV**
**A)** precursor **I**, **B)** precursor **II**, **C)** precursor **III**, **D)** precursor **IV**. Purified compounds were analyzed on a C8 column (EC 250/4.6 Nucleosil 100-5 C8, Macherey-Nagel) using solvent A (0.1% v/v TFA in water) and solvent B (80% v/v acetonitrile in 0.1% v/v TFA in water). For compounds **I** and **II**, the following gradient was applied: 20% B at 0 min, 100% B at 30 min, and 20% B at 31 min. For compounds **III** and **IV**, the gradient was: 50% B at 0 min, 100% B at 30 min, and 50% B at 31 min. The flow rate was 1 mL/min.

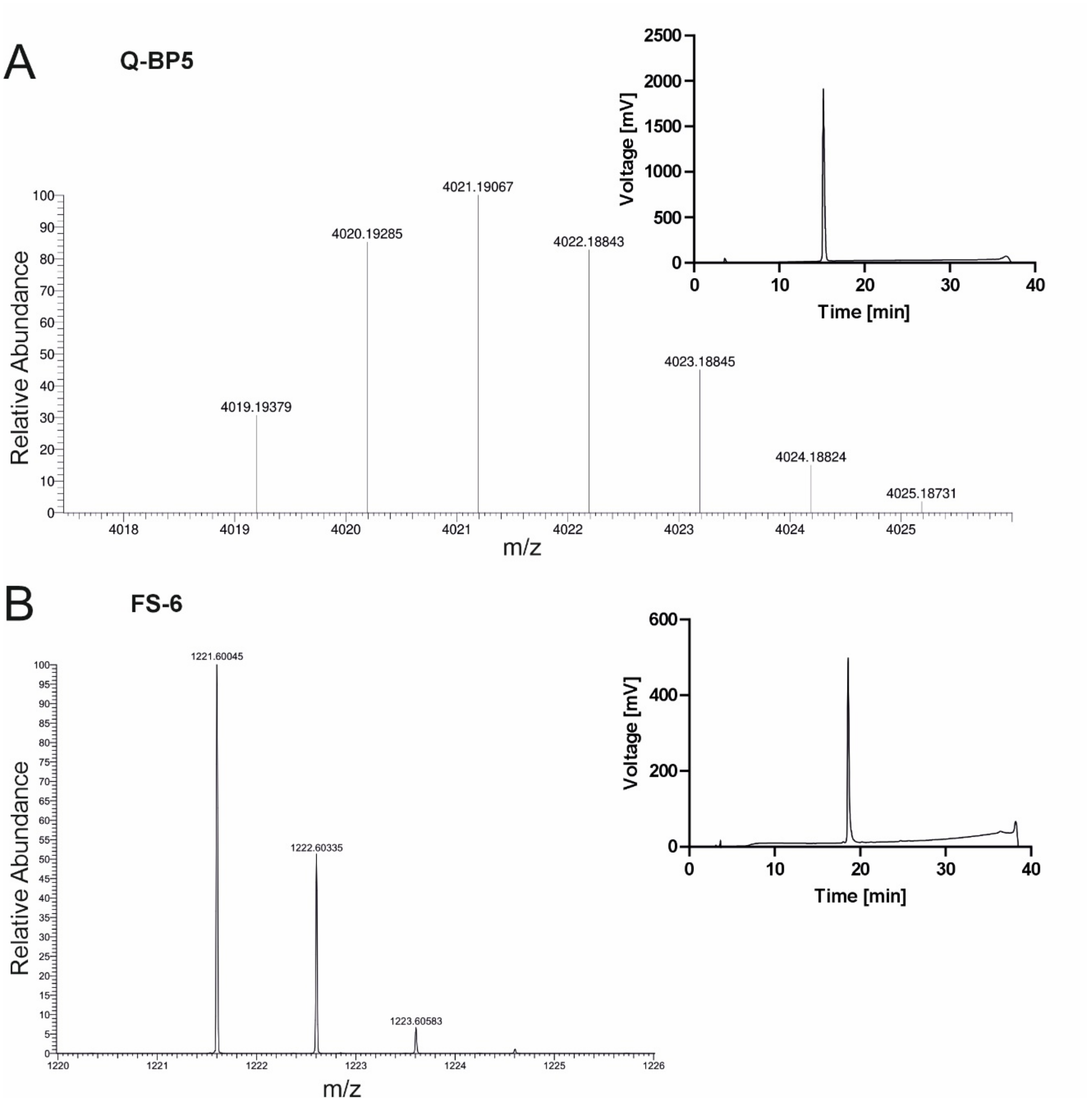


**Figure S4**: **HR-MS and analytical RP-HPLC of fluorescent substrates Q-BP5 and FS-6**
**A**) Q-BP-5 ESI high resolution spectra for $C_{177}H_{291}N_{55}O_{52}$, exact mass 4019.2 and **B**) FS-6 ESI high resolution spectra for C55H81N16O16, exact mass 1221.6011. HPLC chromatograms of purified peptides are in the inserts. Peptides were analyzed on a C8 column (EC 250-4.6 Nucleosil 100-5 C8, Macherey-Nagel) using solvents A (0.1% v/v TFA in water) and B (80% v/v acetonitrile in 0.1% v/v TFA in water) with the following gradient: 10% B at 0 min, 100% B at 30 min, and 10% B at 31 m. The flow rate was 1 mL/min.

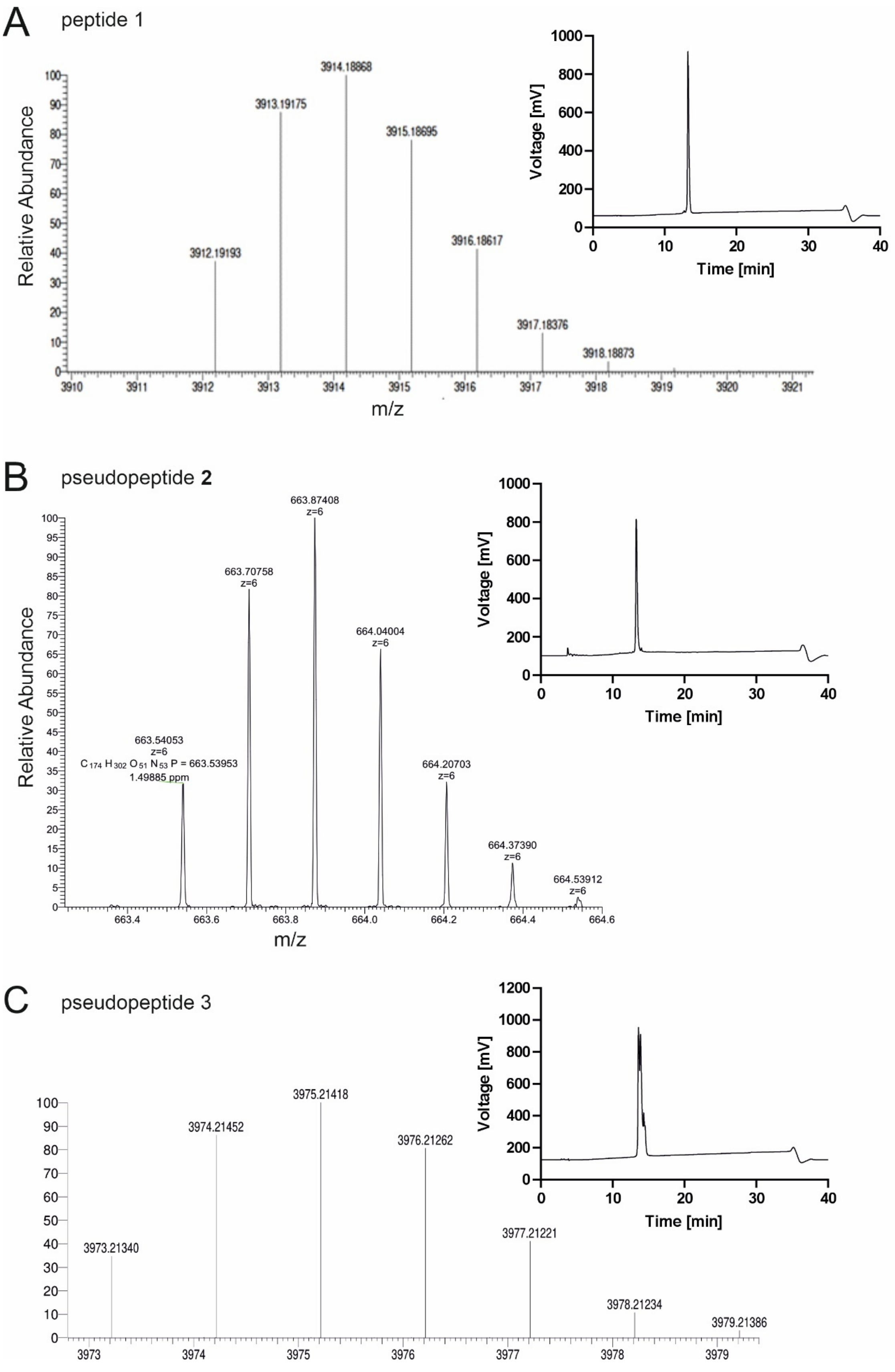
A
peptide 1
Relative Abundance
m/z
3912.19193
3913.19175
3914.18868
3915.18695
3916.18617
3917.18376
3918.18873
Voltage [mV]
Time [min]
B
pseudopeptide **2**
Relative Abundance
m/z
663.54053
z=6
$C_{174}H_{302}O_{51}N_{53}P$ = 663.53953
1.49885 ppm
663.70758
z=6
663.87408
z=6
664.04004
z=6
664.20703
z=6
664.37390
z=6
664.53912
z=6
Voltage [mV]
Time [min]
C
pseudopeptide 3
3973.21340
3974.21452
3975.21418
3976.21262
3977.21221
3978.21234
3979.21386
Voltage [mV]
Time [min]

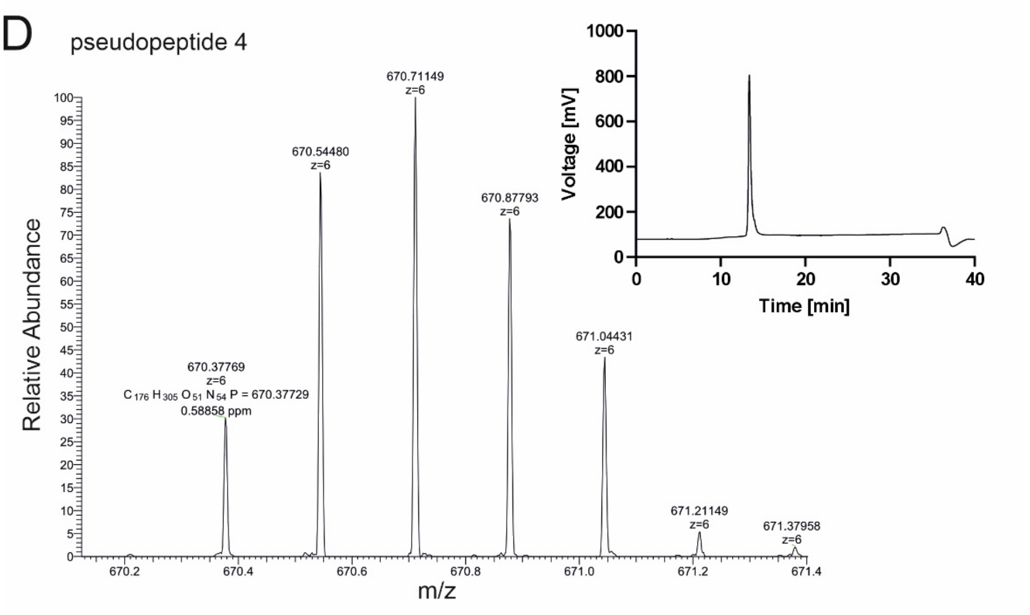


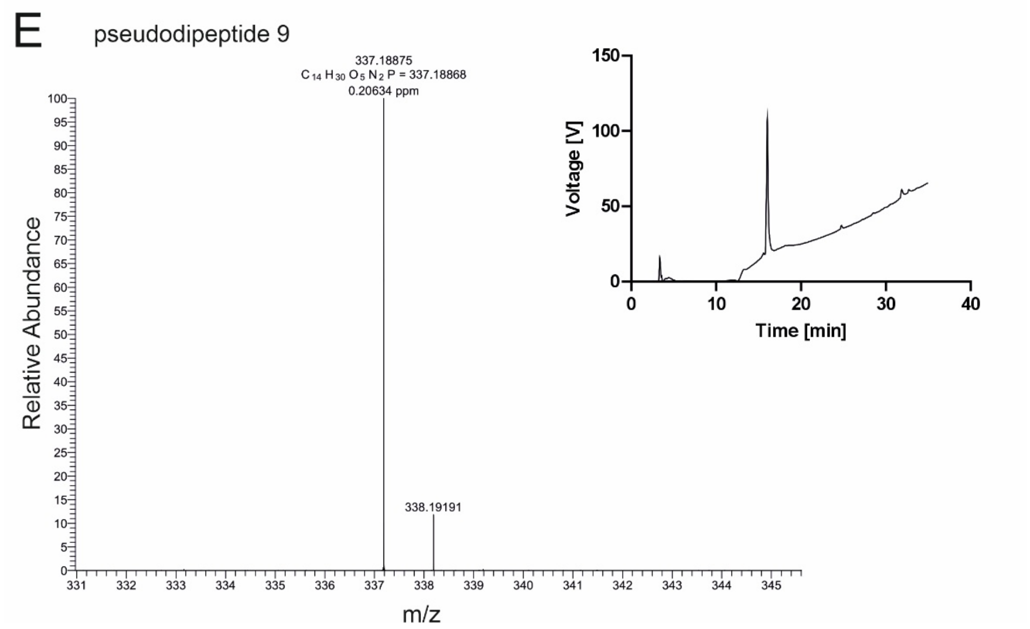


**Figure S5**: **HR-MS and analytical RP-HPLC of purified peptide 1, pseudopeptides 2-4 and pseudodipeptide 9**

**A**) peptide **1** ESI high resolution spectra for $C_{172}H_{290}N_{54}O_{50}$, exact mass 3912.2 **B**) pseudopeptide **2** ESI high resolution spectra for $C_{174}H_{296}N_{53}O_{51}P$, exact mass 3975.19 **C**) pseudopeptide **3** ESI high resolution spectra for $C_{175}H_{298}N_{53}O_{50}P$, exact mass 3973.2 **D**) pseudopeptide **4** ESI high resolution spectra for $C_{176}H_{299}N_{54}O_{51}P$, exact mass 4016.22 and **E**) pseudopeptide **9** ESI high resolution spectra for $C_{14}H_{30}N_{2}O_{5}P$, exact mass 337.1894. HPLC chromatograms of purified peptides are in the inserts. Peptides were analyzed on a C8 column (EC 250-4.6 Nucleosil 100-5 C8, Macherey-Nagel) using solvents A (0.1% v/v TFA in water) and B (80% v/v acetonitrile in 0.1% v/v TFA in water) with the following gradient: 10% B at 0 min, 100% B at 30 min, and 10% B at 31 m in. The flow rate was 1 mL/min.

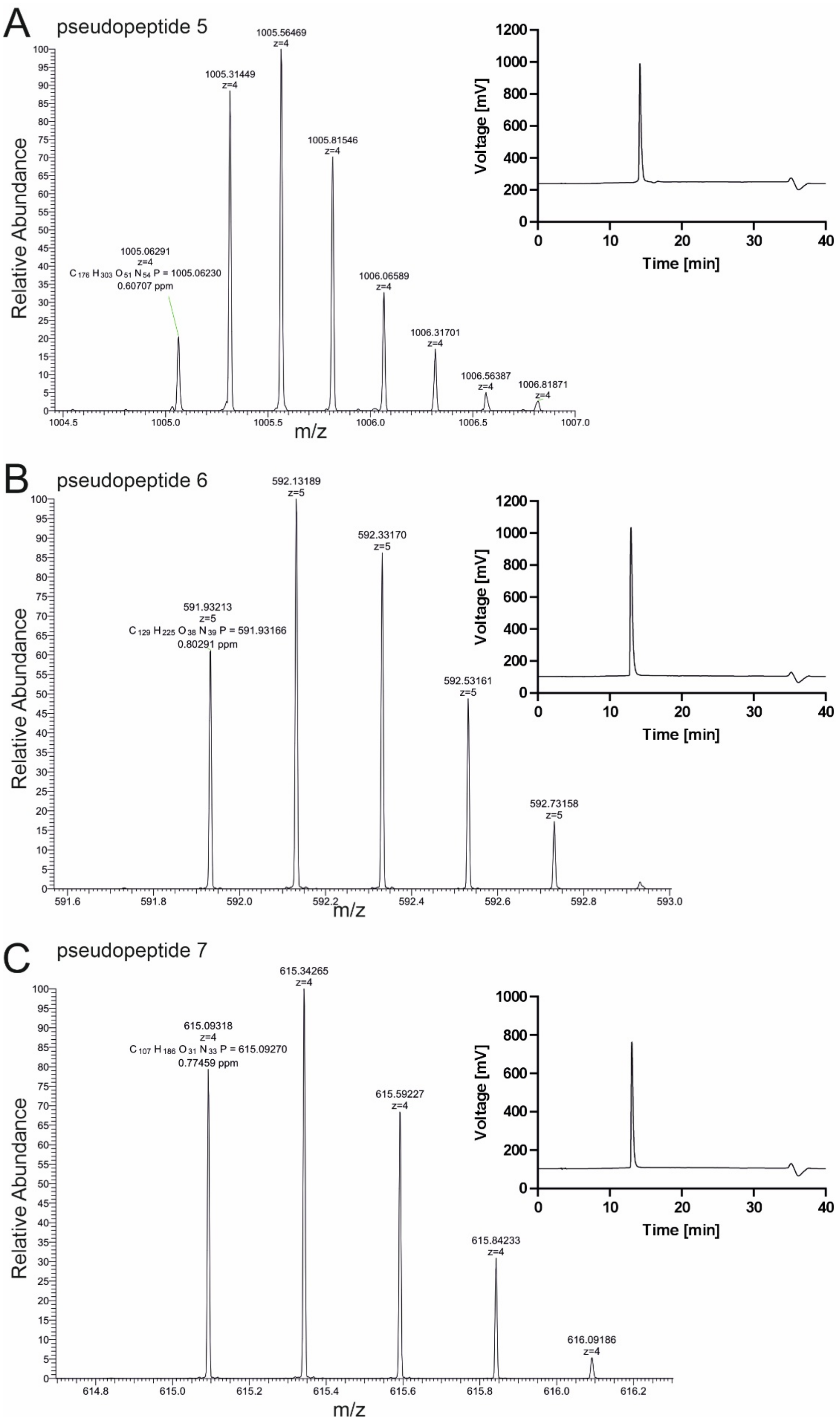
A
pseudopeptide 5
1005.56469
z=4
1005.31449
z=4
1005.81546
z=4
1005.06291
z=4
C 176 H 303 O 51 N 54 P = 1005.06230
0.60707 ppm
1006.06589
z=4
1006.31701
z=4
1006.56387
z=4
1006.81871
z=4
Relative Abundance
m/z
Voltage [mV]
Time [min]
B
pseudopeptide 6
592.13189
z=5
592.33170
z=5
591.93213
z=5
C 129 H 225 O 38 N 39 P = 591.93166
0.80291 ppm
592.53161
z=5
592.73158
z=5
Relative Abundance
m/z
Voltage [mV]
Time [min]
C
pseudopeptide 7
615.34265
z=4
615.09318
z=4
C 107 H 186 O 31 N 33 P = 615.09270
0.77459 ppm
615.59227
z=4
615.84233
z=4
616.09186
z=4
Relative Abundance
m/z
Voltage [mV]
Time [min]

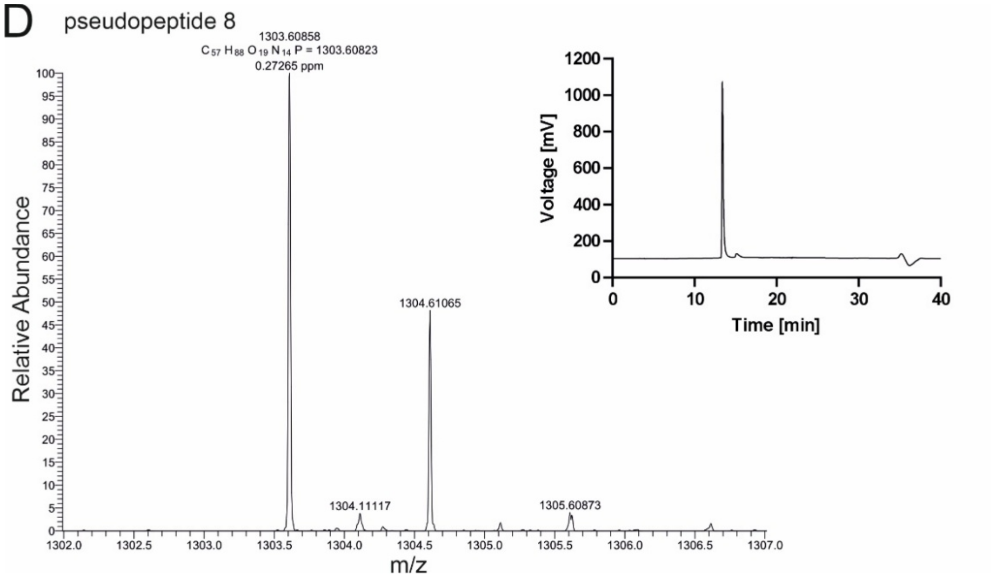


**Figure S6**: **HR-MS and analytical RP-HPLC of purified pseudopeptide 5 and its shortened versions 6-8**

**A**) pseudopeptide **5** ESI high resolution spectra for $C_{176}H_{299}N_{54}O_{51}P$, exact mass 4016.2 **B**) pseudopeptide **6** ESI high resolution spectra for $C_{129}H_{220}N_{39}O_{38}P$, exact mass 2954.6 **C**) pseudopeptide **7** ESI high resolution spectra for $C_{107}H_{182}N_{33}O_{31}P$, exact mass 2456.3 and **D**) pseudopeptide **8** ESI high resolution spectra for $C_{57}H_{87}N_{14}O_{19}P$, exact mass 1302.6. HPLC chromatograms of purified peptides are in the inserts. Peptides were analyzed on a C8 column (EC 250-4.6 Nucleosil 100-5 C8, Macherey-Nagel) using solvents A (0.1% v/v TFA in water) and B (80% v/v acetonitrile in 0.1% v/v TFA in water) with the following gradient: 10% B at 0 min, 100% B at 30 min, and 10% B at 31 m in. The flow rate was 1 mL/min.

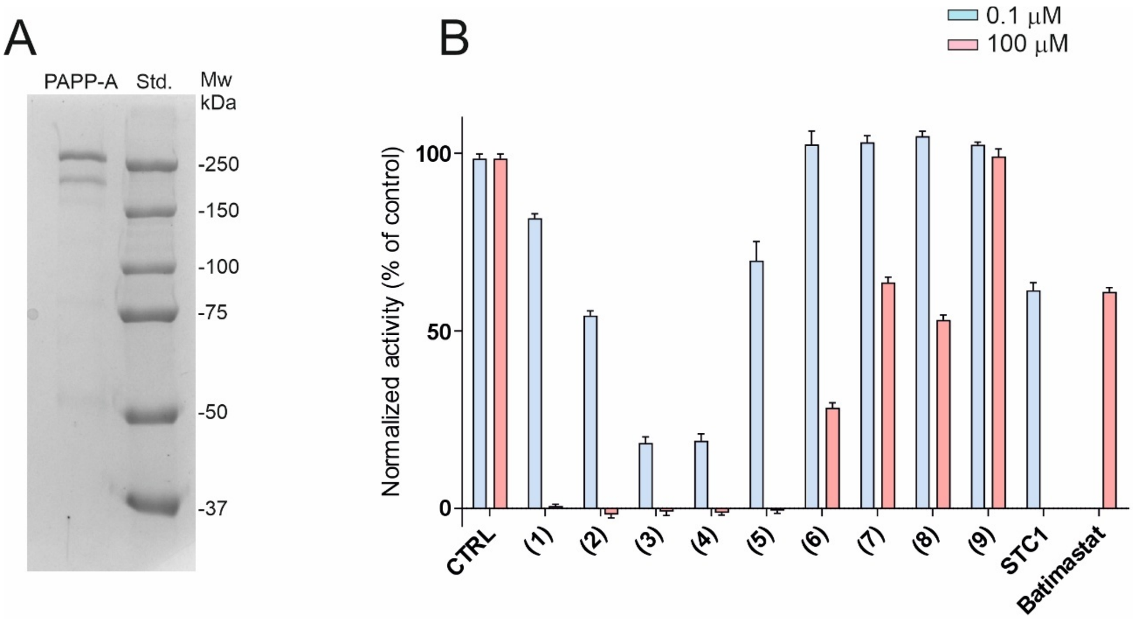


**Figure S7: PAPP-A purification and inhibition of activity**

**A**) Representative Coomassie- stained SDS-PAGE gel (8 %) showing PAPP-A enzyme purified from HEK293T cultivation media. **B**) Inhibition of PAPP-A activity in the quenched fluorescence assay at 0.1 and 100 μM concentrations of peptide **1**, pseudopeptides **2-9** and 0.1 μM STC1 and 100 μM batimastat. Data are shown relative to the extent of cleavage of Q-BP5 without inhibition (CTRL = 100 %) as mean ± SD (n=3).

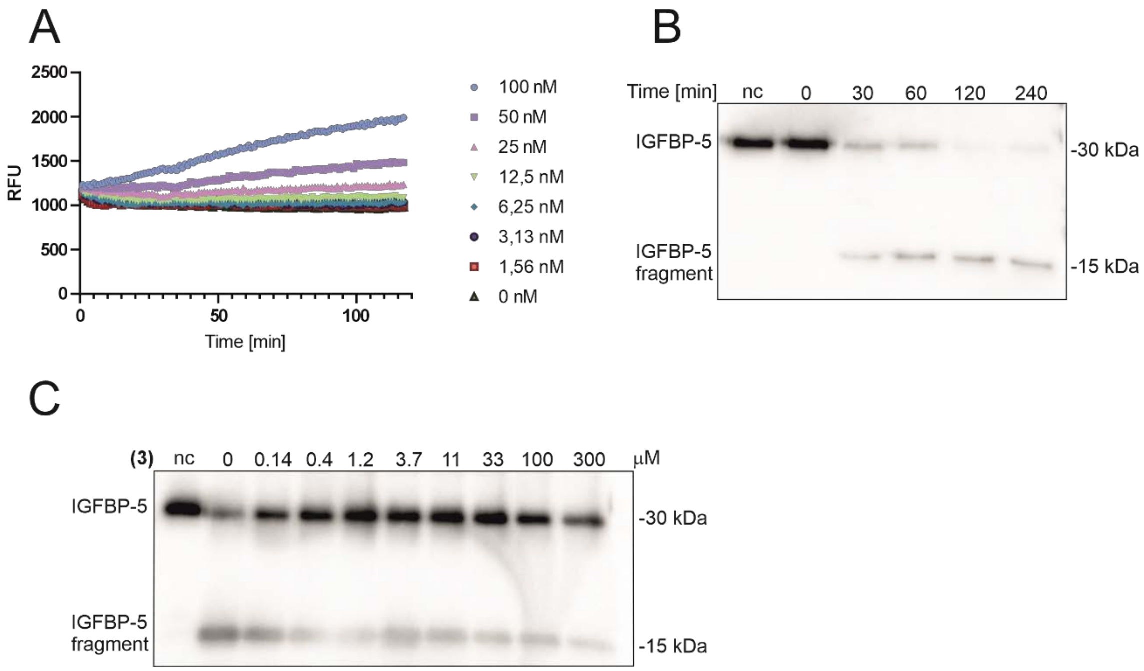


**Figure S8: Activity and inhibition of PAPP-A2.**
**A**) Increase in released fluorescence after cleavage of 5 µM Q-BP5 by PAPP-A2 (Sino Biological) at 0-100 nM concentrations of the enzyme. **B**) Western blot showing time course of IGFBP-5 (333 nM) cleavage by PAPP-A2 (100 nM). **C**) Inhibition of PAPP-A2 cleavage of IGFBP-5 by pseudopeptide **3 (3)**. IGFBP-5 (333 nM) incubated with PAPP-A2 (100 nM) for 1 h at 37 °C in absence or presence of serially diluted psedopeptide **3** (0.14 µM to 300 µM). The proteins and their cleaved fragments were detected using western blot with anti-6 × His tag primary antibody (nc – negative control without enzyme).

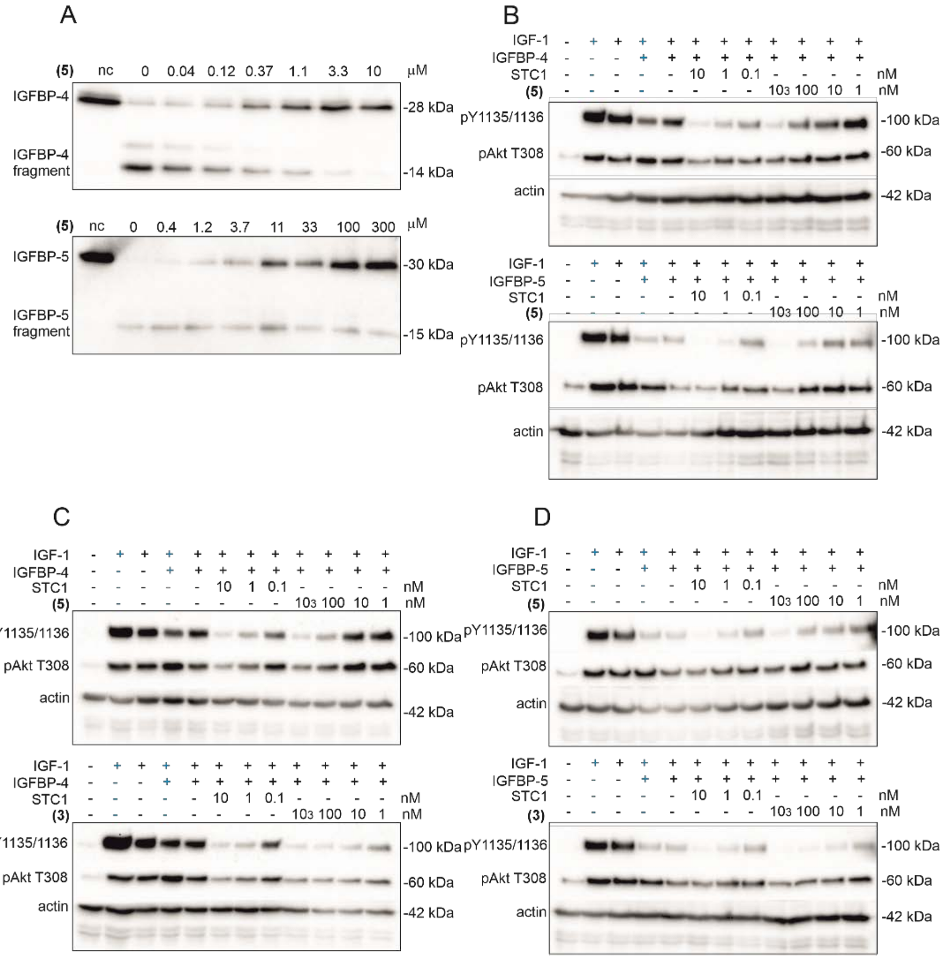


**Figure S9: Inhibition of PAPP-A activity by pseudopeptide 5 and 3. A** and **B**) Representative western blots showing pseudopeptide **5** that are complementary to main Figure 2. **A**) Inhibition of cleavage of IGFBP-4 and IGFBP-5 by pseudopeptide **5 (5)**. IGFBP-4 was pre-incubated with IGF-1 (1 : 3) and incubated with recombinant PAPP-A for 1 h at 37 °C in absence or presence of serially diluted pseudopeptide **5** (0.04 µM to 10 µM). IGFBP-5 was treated in the same conditions without pre-incubation with IGF-1 and the concentration range of pseudopeptide **5** was 0.4 µM to 300 µM. The proteins and their cleaved fragments were detected using western blot with anti-IGFBP-4 antibody and anti-6 × His tag primary antibody for IGFBP-5, (nc – negative control without enzyme). **B**) Inhibition of IGF-1R signaling in A549 cell line by pseudopeptide **5 (5)**. The experiment was performed in the same conditions and with the same batch of cells as in the main **Figure 2C**. Cells were pre-treated with pseudopeptide **5** or STC1 at the indicated concentrations for 1 h. IGF-1 (10 nM) was preincubated with IGFBP-4 or IGFBP-5 (20 nM) for 30 min and added to the cells for two hours (black signs) or 20 min. (blue signs). (-) means that the compound was not present, (+) means treatment with IGF-1 alone or in complex with IGFBPs. **C** and **D**) Comparison of inhibition of IGF-1R signaling in A549 cell line by pseudopeptide **5 (5)** and pseudopeptide **3 (3)** using another batch of cells when incubated with IGF-1 in complex with IGFBP-4 (**C**) and in complex with IGFBP-5 (**D**). Stimulation of IGF-1R signaling was analyzed using standard Western blotting. The membranes were cut at 75 kDa and 50 kDa standards and respective parts were analyzed using phospho-IGF-1Rβ (Tyr1135/1136)/IRβ (Tyr1150/1151) (19H7) (gel top-75 kDa), phospho-Akt (Thr308) (C31E5E) (75-50 kDa) and anti-actin (20-33) (below 50 kDA) as a loading control.

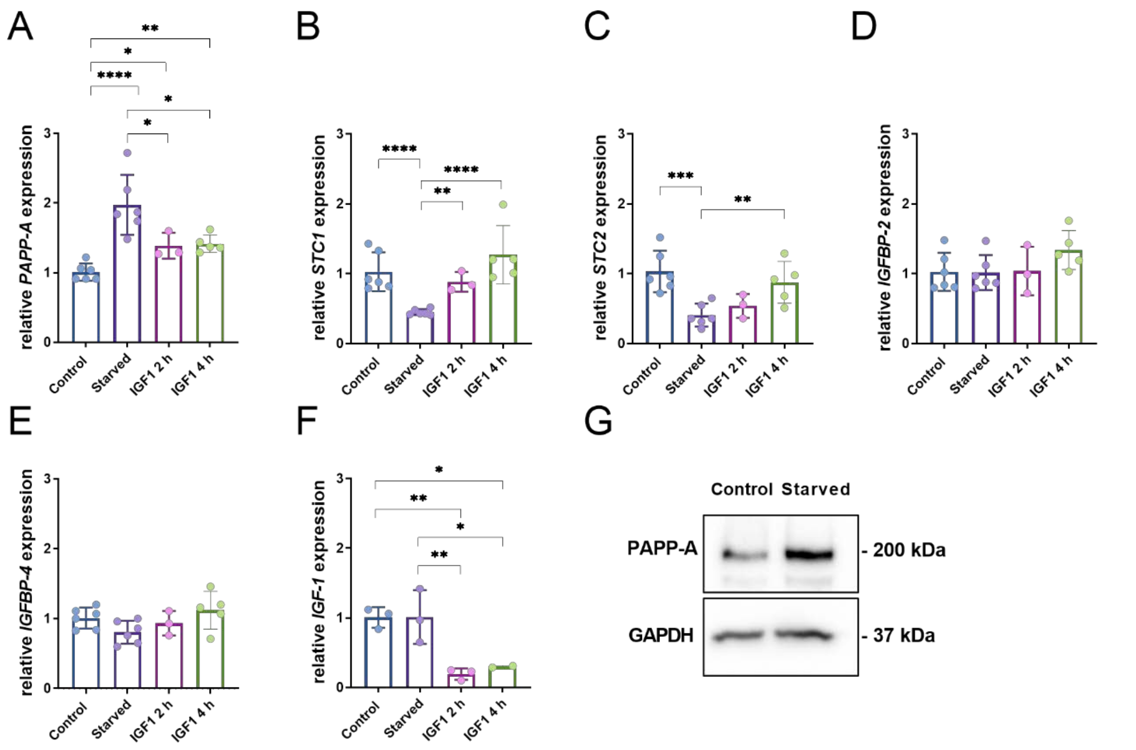


**Figure S10: Expression of genes in the *STC-PAPP-A-IGFBP* axis**

Relative mRNA expression of **A**) *PAPP-A*, **B**) *STC1*, **C**) *STC2*, **D**) *IGFBP-2*, **E**) *IGFBP-4* and **F**) *IGF-1* in A549 cells after overnight incubation in serum-free medium (starved) and subsequent 2 h and 4 h IGF-1 stimulation. Data are shown as mean ± SD. Expression levels were normalized to *GAPDH* and calculated using $2^{-\Delta\Delta Ct}$ method relative to control condition. Statistical significance using one-way ANOVA followed by Tukey's multiple comparison test: *p < 0.05, **p < 0.01, ***p < 0.001, ****p < 0.0001. **G**) Representative western blot showing PAPP-A protein expression in A549 lysates in the control and serum-free incubation (starved) conditions. The membrane was cut on 75 kDa standard and proteins were visualized using anti-PAPP-A antibody and anti-GAPDH antibody as loading control.

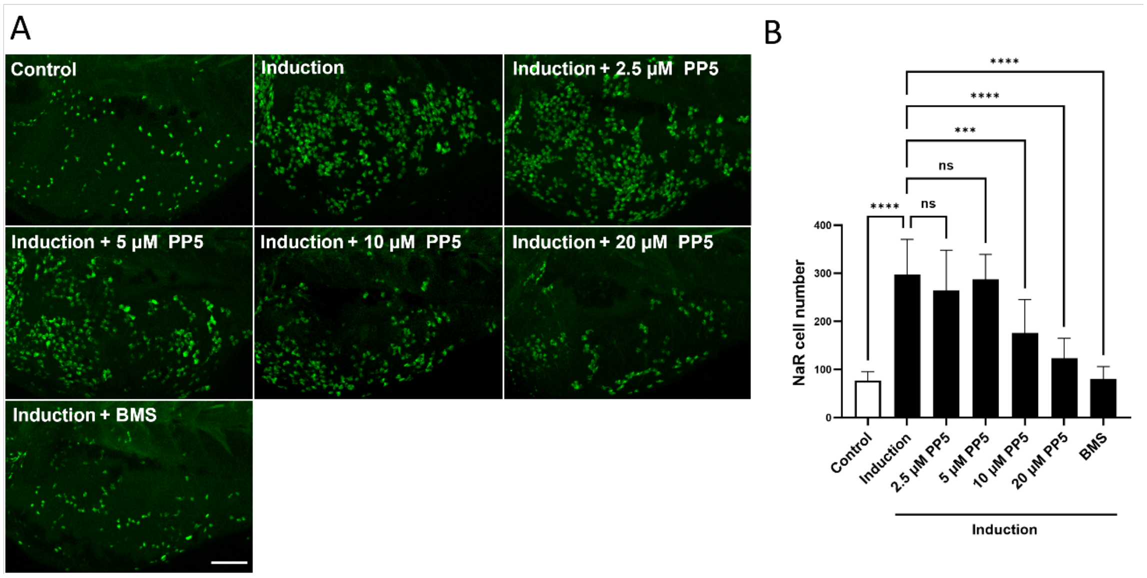


**Figure S11: Pseudopeptide 5 inhibits IGF signaling-mediated cell proliferation *in vivo*** Tg(igfbp5a:GFP) fish were raised to 3 days post-fertilization (dpf) and then transferred to control medium or induction medium. 0.5 μM BMS-754807 (BMS) or Pseudopeptide **5** (PP5) at the indicated concentrations was added, and treatment solutions were changed daily. Two days later, GFP expressing NaR cell numbers in the yolk-sac region in each fish were quantified. Representative images of the yolk-sac region are shown in (**A**), and quantified results are shown in (**B**). Images are lateral views with anterior to the left and dorsal up. Scale bar = 100 μm. Data in **B** are presented as mean ± SD. n = 8–12 fish/group. ns, not significant; ***$p < 0.001$, ****$p < 0.0001$.

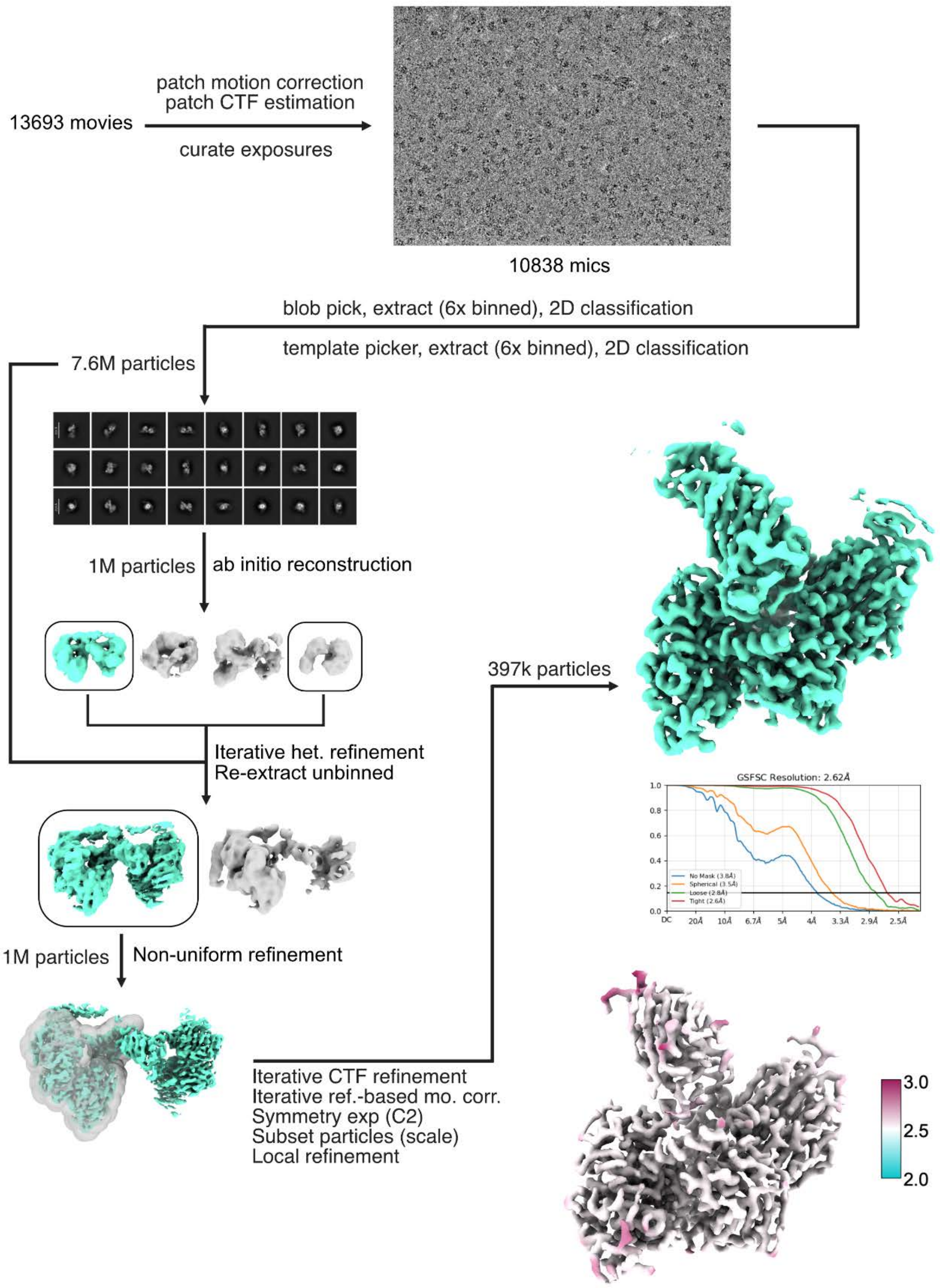


**Figure S12: Cryo-EM processing workflow**

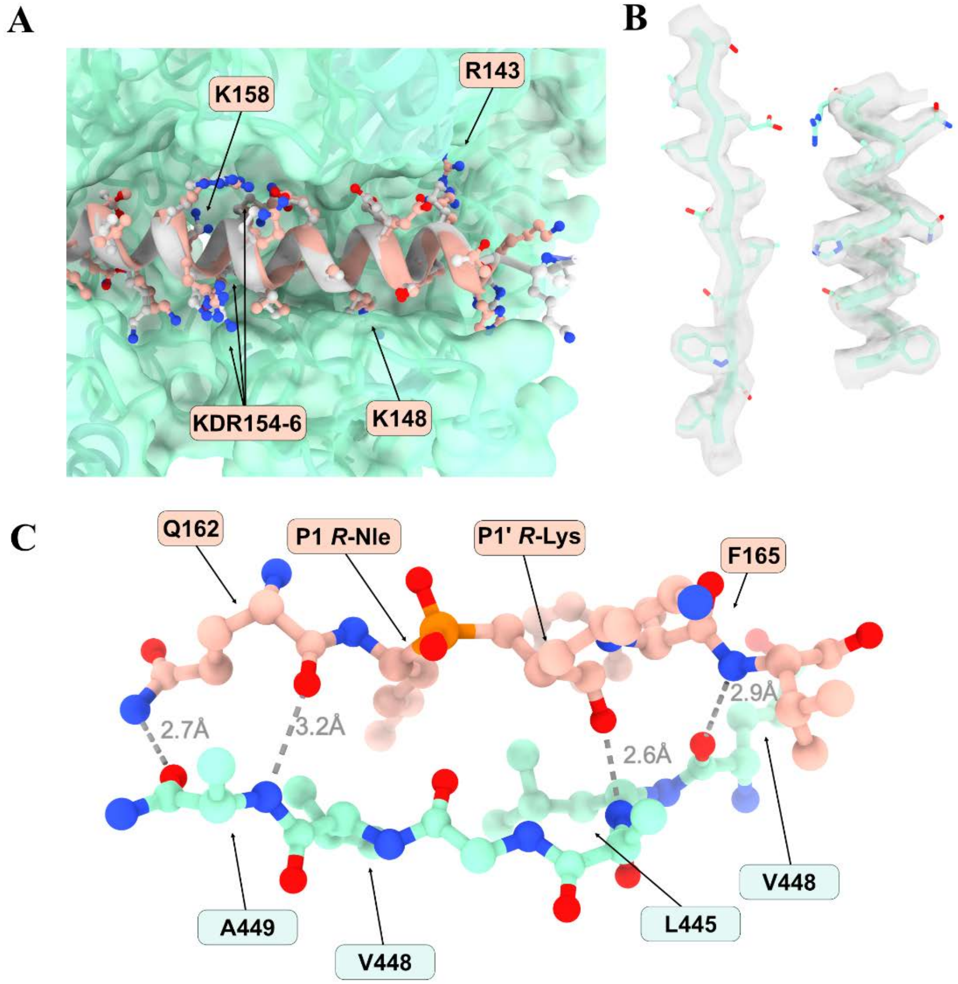


**Figure S13: Supplementary modelling. A)** Overlay of IGFBP5 from PDB 7UFG (grey) with pseudopeptide **3** (salmon) and PAPP-A (turquoise). Key binding residues are indicated with arrows. **B**) Example density fits of beta-strand and alpha-helix to unsharpened cryo-EM map **C**) Pseudo-beta sheet formed between pseudopeptide **3** (salmon) and PAPP-A (turquoise). H-bonds are indicated by dotted lines with N-O distances noted in grey.

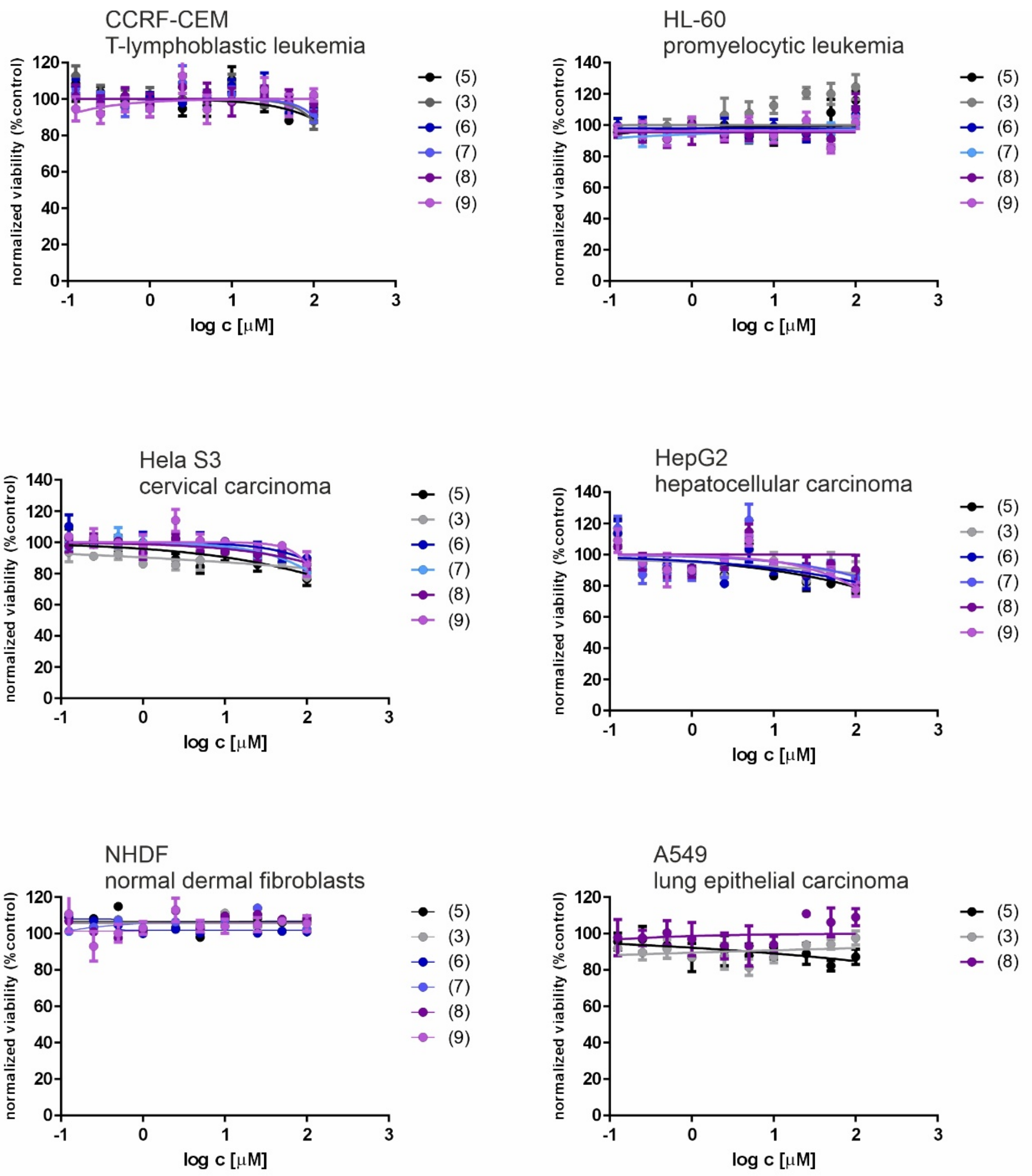


**Figure S14: Cytotoxicity of pseudopeptides 3, 5 and 6-9**

Four cancer cell lines (CCRF-CEM, HepG2, Hela S3, HL-60) and non-tumor human dermal fibroblasts (NHDF) were treated with 100 μM pseudopeptides 72 hours. Cell line A549 was treated with pseudopeptides **3**,**5** and **8**. Cell viability was determined by standard CellTiter-Glo® 2.0 assay (Promega). Luminescence was measured by a multimode plate reader. The signal of the compound-treated cells was related to the value of untreated control which was arbitrarily set to 100% viability. $IC_{50}$ values were calculated from dose-response curves using non-linear regression method using GraphPad Prism software. Data are presented as mean ± SD.

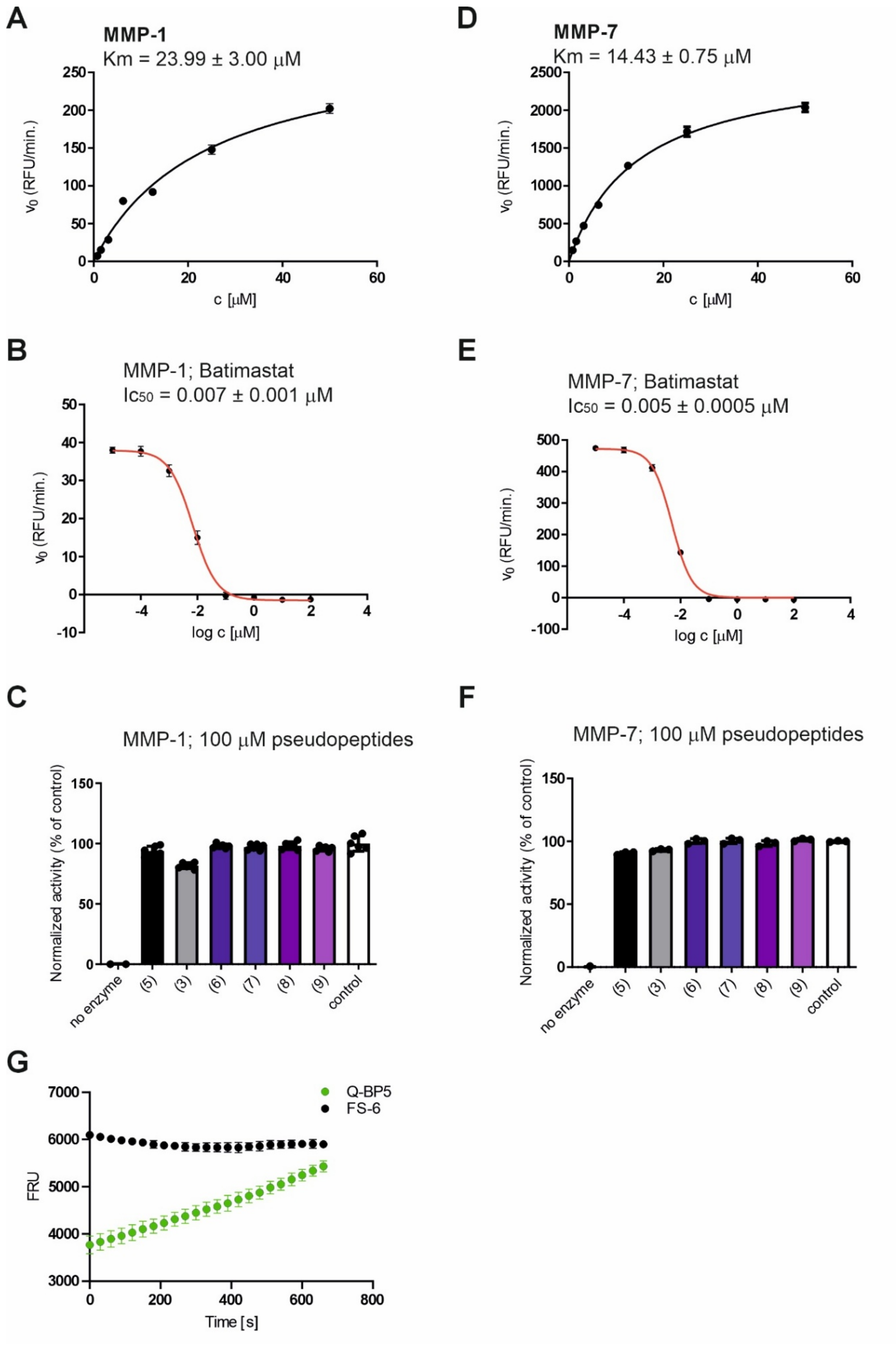
A
MMP-1
Km = 23.99 ± 3.00 μM
v0 (RFU/min.)
c [μM]
D
MMP-7
Km = 14.43 ± 0.75 μM
v0 (RFU/min.)
c [μM]
B
MMP-1; Batimastat
Ic50 = 0.007 ± 0.001 μM
v0 (RFU/min.)
log c [μM]
E
MMP-7; Batimastat
Ic50 = 0.005 ± 0.0005 μM
v0 (RFU/min.)
log c [μM]
C
MMP-1; 100 μM pseudopeptides
Normalized activity (% of control)
no enzyme
(5)
(3)
(6)
(7)
(8)
(9)
control
F
MMP-7; 100 μM pseudopeptides
Normalized activity (% of control)
no enzyme
(5)
(3)
(6)
(7)
(8)
(9)
control
G
Q-BP5
FS-6
FRU
Time [s]

**Figure S15: Inhibition of MMP-1 and MMP-7 with pseudopeptides 3, 5 and 6-9**

Activity of MMP-1 (**A**) and MMP-7 (**D**) was assessed by quenched fluorescence assay using fluorogenic substrate FS-6. Cleavage of the FS-6 substrate was followed at 325 nm and emission at 400 nm. The FS-6 substrate was serially diluted to yield final concentrations 0.78 – 100 µM to assess $K_m$ value of the enzymes, 10 nM MMP-1 and 2 nM MMP-7 were used. For $IC_{50}$ measurement of inhibition of cleavage of FS-6 by MMP-1 (**B**) and MMP-7 (**E**) with batimastat, the FS-6 concentration was set at 5 µM and batimastat was tested in concentration range $10^{-11}$ - $10^{-4}$ M. The inhibition of MMP-1 (**C**) and MMP-7 (**F**) with pseudopeptides was tested at single concentration 100 µM and normalized to activity of the enzymes with no inhibitor (control). The data were analyzed using GraphPad Prism 8 (GraphPad, USA), applying Michaelis-Menten non-linear regression model for $K_m$ measurement. $IC_{50}$ values were assessed by non-linear fitting of log(inhibitor) *vs* response variable slope. **G**) Time profile of cleavage of substrates Q-BP5 and FS-6 at 10 µM with PAPP-A followed at 315 nm and emission at 410 nm. Data are presented as mean ± SD.

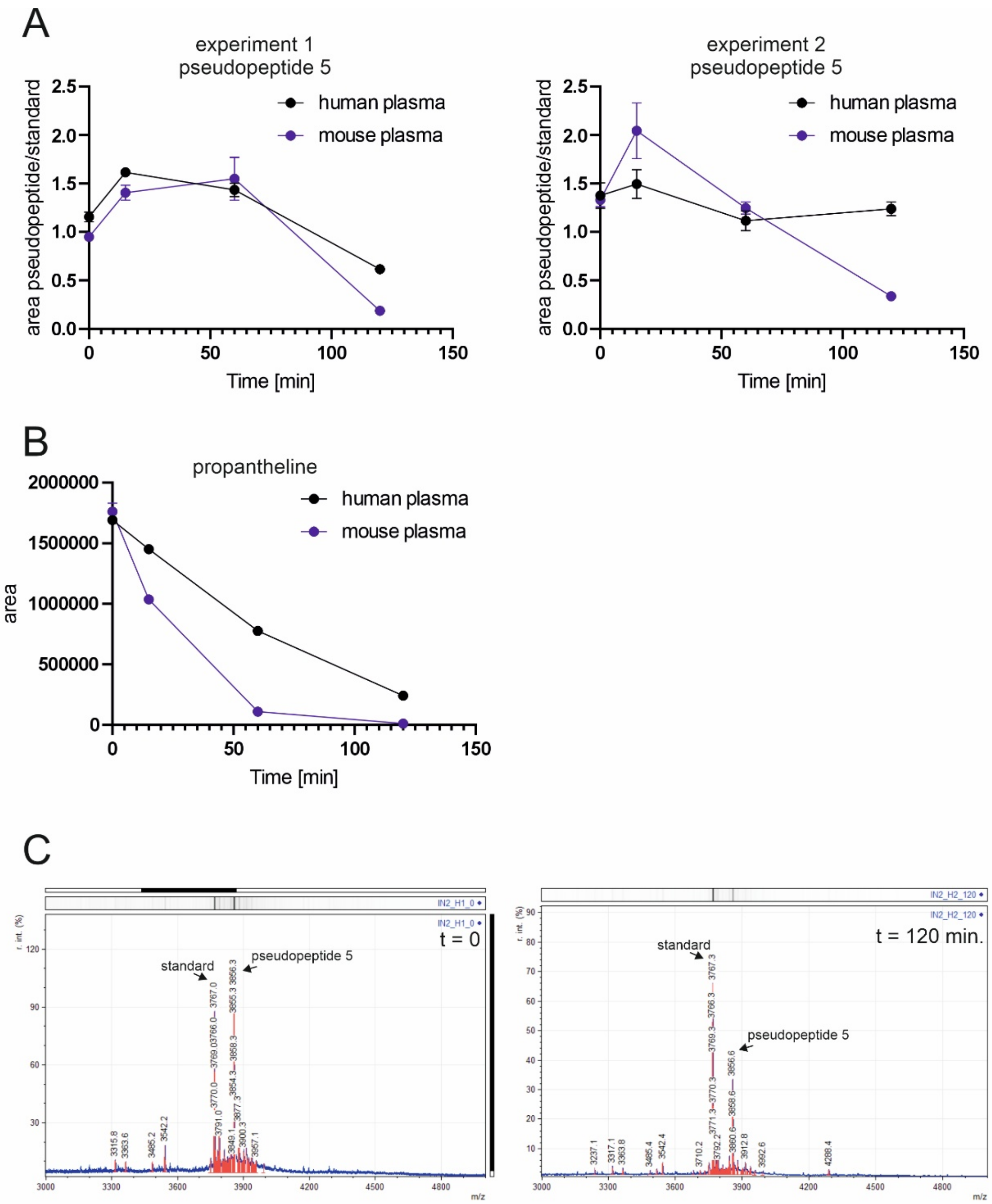


**Figure S16: Stability of pseudopeptide 5 in plasma**

**A)** Intensity of peak corresponding to pseudopeptide 5 (Mw 3856.5) was related to intensity of added standard (Mw 3767) in human and mouse plasma in individual time points (0, 15, 60 and 120 min.). Each point was measured 6x. Data are presented as mean ± SD. **B**) Control experiment showing degradation of propantheline under the same conditions as the experiments with pseudopeptide 5. **C**) Representative MALDI-TOF spectra of pseudopeptide 5 in human plasma at t = 0 and t = 120 min. from experiment 1 shown in A.

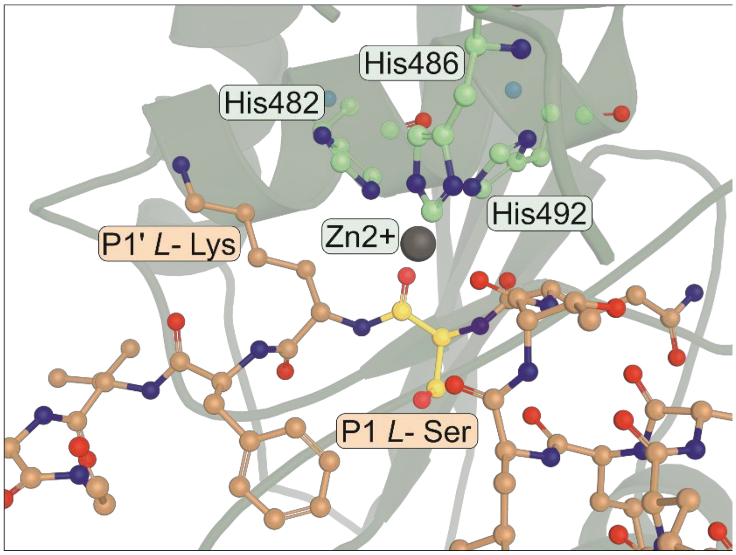


**Figure S17: AlphaFold 3 modeled active site of native peptide.** Closeup view of AlphaFold 3 predicted conformation of peptide **1** (salmon) in active site of PAPP-A (turquoise) with key residues in atomic details.

**Table S1: Averaged root-mean-square deviations for trajectories in MD simulation**

S-RMSD of the backbone atoms and C-RMSD of the entire complex values and their standard deviations (SD) for each trajectory of simulated system consisting of the truncated PAPP-A fragment (386 residues), the IGFBP5(138-171)+Y substrate (34 residues), and a zinc ion ($Zn^{2+}$). Raw data are available via the Open Science Framework repository: 10.17605/OSF.IO/7CZ2B

| Run | S-RMSD ± SD | C-RMSD ± SD |
|---|---|---|
| 1 | 3.52 ± 0.34 | 3.77 ± 0.10 |
| 2 | 3.98 ± 0.32 | 3.75 ± 0.33 |
| 3 | 4.15 ± 0.15 | 4.60 ± 0.14 |
| 4 | 3.14 ± 0.22 | 2.90 ± 0.17 |

**Table S2: Refinement statistics for cryo-EM structure**

| Model | | |
|---|---|---|
| Composition (#) | | |
| Chains | 4 | |
| Atoms | 8391 (Hydrogens: 0) | |
| Residues | Protein: 1049 Nucleotide: 0 | |
| Water | 0 | |
| Ligands | R5M: 1 | |
| | NAG: 7 | |
| | BMA: 1 | |
| | ZN: 1 | |
| Bonds (RMSD) | | |
| Length (Å) (# > 4σ) | 0.004 (0) | |
| Angles (°) (# > 4σ) | 0.949 (0) | |
| MolProbity score | 1.00 | |
| Clash score | 0.61 | |
| Ramachandran plot (%) | | |
| Outliers | 0.00 | |
| Allowed | 2.88 | |
| Favored | 97.12 | |
| Rama-Z (Ramachandran plot Z-score, RMSD) | | |
| whole (N = 1041) | -0.25 (0.25) | |
| helix (N = 133) | -1.17 (0.40) | |
| Sheet | 0.49 (0.30) | |
| Loop | -0.18 (0.24) | |
| Rotamer outliers (%) | 1.52 | |
| Cβ outliers (%) | 0.10 | |
| Peptide plane (%) | | |
| Cis proline/general | 3.1/0.0 | |
| Twisted proline/general | 0.0/0.0 | |
| CaBLAM outliers (%) | 1.45 | |
| ADP (B-factors) | | |
| Iso/Aniso (#) | 8391/0 | |
| min/max/mean | | |
| Protein | 49.25/228.78/125.94 | |
| Nucleotide | --- | |
| Ligand | 20.00/177.55/126.45 | |
| Water | --- | |
| Occupancy | | |
| Mean | 1.00 | |
| occ = 1 (%) | 100.00 | |
| 0 < occ < 1 (%) | 0.00 | |
| occ > 1 (%) | 0.00 | |
| **Data** | | |
| Box | | |
| Lengths (Å) | 96.55, 104.50, 107.91 | |
| Angles (°) | 90.00, 90.00, 90.00 | |
| Supplied Resolution (Å) | 2.6 | |
| Resolution Estimates (Å) | Masked | Unmasked |
| d FSC (half maps; 0.143) | --- | --- |
| d 99 (full/half1/half2) | 3.3/---/--- | 3.2/---/--- |
| d model | 2.9 | 2.9 |
| d FSC model (0/0.143/0.5) | 2.5/2.6/2.9 | 2.5/2.6/3.0 |
| Map min/max/mean | -0.14/0.48/0.01 | |
| **Model vs. Data** | | |
| CC (mask) | 0.83 | |
| CC (box) | 0.81 | |
| CC (peaks) | 0.69 | |
| CC (volume) | 0.83 | |
| Mean CC for ligands | 0.75 | |

**Table S3: Primers for qRT-PCR**

| | Forward | Reverse |
|---|---|---|
| **PAPP-A** | ACAAAGACCCACGCTACTTTTT | CATGAACTGCCCATCATAGGTG |
| **PAPP-A2** | GCATCTCAGCTGTGGCTCTA | CAAGCAGCAATGACGGACAG |
| **STC1** | CATCCCTATGGGCTTGGACC | AGTCTCCCACCCCATCATCA |
| **STC2** | CAGCGSGGAATGCTACCTCAA | ACAGGTCAGCAGCAAGTTCA |
| **IGFBP-2** | CGAGGGCACTTGTGAGAAGC | CCAGCTCCTTCATACCCGAC |
| **IGFBP-4** | ACCCACGAGGACCTCTACATCA | CTTCCGGTCCACACACCAGC |
| **IGFBP-5** | TTCCCTGCAGGAGCTCAAAG | GGCAGCTTCATCCCGTACTT |
| **IGF-1** | [illegible] | [illegible] |
| **IGF-2** | [illegible] | [illegible] |
| **Actin** | [illegible] | [illegible] |
| **GAPDH** | GAAGGTGAAGGTCGGAGTCAAC | CAGAGTTAAAAGCAGCCCTGGT |

### 1. Synthetic methods

Reagents and solvents were purchased from commercial suppliers (Merck, Iris Biotech GmBH or Fluorochem, Ltd., Penta chemicals, Lach-Ner, VWR Chemicals) and were of analytical grade. Starting compounds were purchased from the company BLD Pharm. TLC was performed on silica gel coated aluminium plates (Merck). The compounds were visualized by exposure to UV light at 254 nm or by ninhydrin spraying followed by heating (dark blue stain of BocHN, free amine). Flash chromatography were carried out on silica gel (40-63 μm, VWR Chemiclas). Melting points were determined on a Boetius block and are uncorrected. Analytical RP-HPLC chromatography was carried out at flow rate 1 ml/min on C8 column (Macherey – Nagel 120-5C8; 250 x 4 mm, 5μm). $^{1}$H and $^{13}$C NMR spectra were acquired on Bruker AVANCE-600 spectrometer ($^{1}$H at 600.13 MHz, $^{13}$C at 150.9 MHz, $^{31}$P NMR at 202.4 MHz) in $CDCl_3$ or DMSO-$d_6$ at 300 K. The 2D H H COSY, 2D H C HSQC and 2D H C HMBC spectra were recorded and used for the structural assignment of proton and carbon signals. IR spectra were recorded on a Bruker IFS 55 Equinox apparatus. HRMS data were obtained on a FTMS mass spectrometer LTQ-orbitrap XL (Thermo Fisher, Bremen, Germany) in electrospray ionization mode. Optical rotation values were measured in 100 mm cell on Perkin Elmer 241 MC under Na lamp radiation.

#### 1.1: Synthesis of modified amino acids

Syntheses of the protected amino acids *o*-aminobenzoic acid-modified lysine (compound **18**) and 3-nitrotyrosine (compound **20**) used for preparation of fluorogenic substrate Q-BP5 are shown in **Scheme 1**. The synthesis started from commercially available Boc-Lys(Cbz)-OH **13**, which was first converted into the corresponding benzyl ester **14**. Subsequent treatment with TFA resulted in complete removal of the Boc protecting group from the ω-amino functionality, providing trifluoroacetate salt **15**, which was used without additional isolation or characterization.

Acylation with the 2-(Boc-amino)benzoyl moiety was then carried out either directly using Boc-2-Abz-OH **11** or *via* its NHS-activated derivative **12**. From the isolated ester **16**, both the benzyl and benzyloxycarbonyl protecting groups were removed in a single hydrogenolysis step to yield the zwitterionic compound **17**. Introduction of the Fmoc protecting group at the α-amino moiety subsequently provided compound **18**, suitable for solid-phase peptide synthesis applications.

Next, commercially available 3-nitrotyrosine **19** was converted into the corresponding Fmoc-protected amino acid **20** by standard Fmoc protection of the amino group, isolated material fulfilled all solid-phase synthesis requirements.

Analytical HPLC profiles of the final compounds **18** and **20** are shown in **Figure S1**.

**1.1.1: Synthetic protocols for compounds in Scheme S1**

2-[(*tert*-Butoxycarbonyl)amino]benzoic acid **11**

A mixture of anthranilic acid **10** (8.2 g, 60 mmol) and $NaHCO_3$ (15.1 g, 180 mmol) in water (150 mL) was stirred at ambient temperature. A solution of $Boc_2O$ (15.7 g, 72 mmol) in dioxane (150 mL) was added dropwise, and the resulting reaction mixture was allowed to react for 72 h at room temperature. Upon completion of the reaction, the mixture was cooled in an ice bath and acidified with 2 M citric acid until the pH ~ 3 was reached as monitored by pH-indicator paper. The reaction mixture was transferred to a separatory funnel, diluted with water (200 mL), and extracted with ethyl acetate (4 × 150 mL). The combined organic phases were washed successively with water (4 × 100 mL), brine (2 × 100 mL) and dried over anhydrous $Na_2SO_4$. The drying agent was removed by filtration, and the filtrate was concentrated under reduced pressure to dryness, affording 13.1 g of a yellow residue. The residue was dissolved in methanol (50 mL) and loaded on an ion-exchange column packed with 50 g Dowex 50 in the $H^+$ form. The column was eluted with a methanol:water (1:1) mixture (total wash volume ~2 L). The methanol from the combined eluate was removed under reduced pressure, during which significant precipitation occurred. The precipitate was collected by vacuum filtration on a Büchner funnel and washed with chilled water (50 mL), dried over $P_2O_5$ in a desiccator overnight, and recrystallized from ethyl acetate. Yield 10 g (70%). Colorless solid, m. p. 159 – 161°C. $R_f$ = 0.59 (DCM : MeOH : $NH_4OH$ 75 : 22 : 3).

$^1H$ NMR (600 MHz, DMSO; T = 25 °C): 1.55 s (9H, 3x $-CH_3$, *t*-Bu), 7.05 ddd (1H, *J* = 8.1, 7.2 and 1.2 Hz, Ar-H), 7.57 ddd (1H, *J* = 8.6, 7.2 and 1.6 Hz, Ar-H), 8.12 dd (1H, *J* = 8.1 and 1.6 Hz, Ar-H), 8.48 dd (1H, *J* = 8.6 and 1.2 Hz, Ar-H), 10.01 br (1H, >NH). $^{13}C$ NMR (150.9 MHz, DMSO; T = 25 °C): 28.30 (3x $-CH_3$, *t*-Bu), 80.88 (>C<, *t*-Bu), 113.18 (Ar-C=, $C_6H_5$), 118.95 (Ar-CH=, $C_6H_5$), 121.34 (Ar-CH=, $C_6H_5$), 131.91 (Ar-CH=, $C_6H_5$), 135.63 (Ar-CH=, $C_6H_5$), 142.92 (Ar-C=, $C_6H_5$), 152.75 (N-CO-O), 172.90 (-COOH). IR (KBr, $\nu_{max}$ $cm^{-1}$) 3319 m (NH); 2984 s, 2878 m, 1455 s, 1390 s, 1369 s ($CH_3$); 1728 vs (C=O) acid; 1670 vs (C=O) carbamate, 1534 s (amide II); 3077 m, 1606 s, 1587 vs, 1474 s, 1251 vs, 1150 vs, 777 m, 759 s (benzene ring); 1170 s ((C($CH_3$)$_3$). HRMS ESI calcd for $C_{12}H_{15}O_4NNa$ $[M+Na]^+$ 260.08933; found: 260.08950.

Succinimidyl 2-[(*tert*-butoxycarbonyl)amino]benzoate **12**

Acid **11** (7.7 g, 29.9 mmol) and NHS (3.4 g, 29.9 mmol) were dissolved in THF (120 mL). While the solution was stirred and cooled in an ice bath, DIC (4.7 mL, 29.9 mmol) was added dropwise. The reaction mixture was stirred at 0 °C for 1 h and then at room temperature overnight. After completion of the reaction, the precipitated DIU was removed by filtration, and the filter cake was washed with cold THF (50 mL). The combined filtrates were concentrated under reduced pressure, and the resulting residue was dissolved in ethyl acetate (100 mL). The solution was placed in a refrigerator at 5 °C overnight to induce crystallization. The second crop of DIU was removed by filtration, and the solid was washed with cold ethyl acetate (20 mL). The filtrate was concentrated under reduced pressure, and the resulting solid was dissolved in the minimal amount of hot ethyl acetate. Petroleum ether was added until crystal formation was observed. The precipitated crystals were collected by filtration, washed with petroleum ether (50 mL), and dried under high vacuum. The product (8.9 g) was used without further purification in the next reaction step.

Benzyl 2-(*S*)-[(benzyloxycarbonyl)amino]-6-[(*tert*-butoxycarbonyl)amino]hexanoate **14**

A suspension of 2-(*S*)-[(benzyloxycarbonyl)amino]-6-[(*tert*-butoxycarbonyl)amino]hexanoic acid **13** (7.6 g, 20 mmol), benzyl bromide (2.6 mL, 22 mmol), and potassium carbonate (3.0 g, 22 mmol) in anhydrous DMF (100 mL) was stirred under a calcium chloride drying tube overnight at room temperature. After completion of the reaction, the mixture was cooled in an ice bath and diluted with chilled water (300 mL). The resulting mixture was transferred to a separatory funnel and extracted with ethyl acetate (4 × 100 mL). The combined organic layers were washed successively with water (100 mL) and brine (2 × 10 mL), dried over anhydrous $Na_2SO_4$, and filtered. The filtrate was concentrated under reduced pressure to give a crude residue, which was purified by column chromatography on silica gel using a linear gradient of ethyl acetate in toluene to afford a colorless oil. The oil was triturated at –20 °C with a mixture of ethyl acetate and petroleum ether, yielding the product as a crystalline solid. Yield 7,9 g (84%). White solid, m. p. 75 – 77 °C. $R_f$ = 0.52 (ethyl acetate : toluene 20 : 80). $[\alpha]_D^{20}$ = -5.6 (c = 2 ; $CHCl_3$ ).

$^1$H NMR (600 MHz, $CDCl_3$; T = 25 °C): 1.28 m + 1.34 m (2H, >$CH_2$), 1.43 s (9H, 3x -$CH_3$, *t*-Bu), 1.43 m (2H, >$CH_2$), 1.69 m + 1.85 m (2H, >$CH_2$), 3.05 m (2H, -$CH_2$-N), 4.40 td (1H, *J* = 7.9 and 5.1 Hz, >CH-N), 4.52 br (1H, >N-H), 5.09 d, *J* = 12.0 Hz and 5.1 d, *J* = 12.0 Hz (2H, -$CH_2$-O), 5.14 d, *J* = 11.8 Hz and 5.19 d, *J* = 11.8 Hz (2H, -$CH_2$-O), 7.31 - 7.37 m (10H, Ar-CH=, 2x $C_6H_5$).$^{13}$C NMR (150.9 MHz, $CDCl_3$; T = 25 °C): 22.22 (>$CH_2$), 28.38 (3x -$CH_3$, *t*-Bu), 29.52 (>$CH_2$), 32.12 (>$CH_2$), 39.97 (-$CH_2$-N), 53.75 (>CH-N), 66.97 (-$CH_2$-O), 67.10 (-$CH_2$-O), 79.12 (>C<, *t*-Bu), 128.10 (Ar-CH=, $C_6H_5$), 128.14 (2x Ar-CH=, $C_6H_5$), 128.31 (2x Ar-CH=, $C_6H_5$), 128.47 (Ar-CH=, $C_6H_5$), 128.48 (2x Ar-CH=, $C_6H_5$), 128.60 (2x Ar-CH=, $C_6H_5$), 135.57 (Ar-C=, $C_6H_5$), 136.20 (Ar-C=, $C_6H_5$), ), 155.92 (N-CO-O), 156.00 (N-CO-O), 172.22 (-COOH). IR ($CHCl_3$, $\nu_{max}$ $cm^{-1}$) 3360 vs, 3346 s (NH); 2985 m, 2869 m 1466 m, 1391 m, 1364 m ($CH_3$); 2935 m, 1454 m ($CH_2$); 1746 vs (C=O) ester; 1718 vs, 1686 vs (C=O) carbamates; 1532 s (amide II); 3092 w, 3063 w, 3036 w, 1609 w, 1314 m, 1272 s, 1084 m, 1030 m, 998 m, 753 m (benzene rings);

1164 s ((C($CH_3$)$_3$); 1061 m, 1251 s (C-O). HRMS ESI calcd for $C_{26}H_{35}O_6N_2$ [M+H]$^+$ 471.24896; found: 471.24856.

Benzyl 2-(*S*)-[(benzyloxycarbonyl)amino]-6-[[2-(*tert*butoxycarbonyl)amino]benzoylamino] hexanoate **16**

Ester **14** (9.3 g, 26.6 mmol) was dissolved in DCM (40 mL), and the solution was cooled in an ice bath. Trifluoroacetic acid (40 mL) was added dropwise, and the reaction mixture was stirred at 0 °C for 1 h, followed by an additional 1 h at room temperature. TLC chromatography indicated complete consumption of the starting material. The volatile components were removed under reduced pressure to afford a light brown oil, which was used directly in the subsequent step without further purification.

Benzyl 2-(*S*)-[(benzyloxycarbonyl)amino]-6-aminohexanoate trifluoroacetic acid salt **15** was dissolved in DCM (50 mL) and cooled in an ice bath. TEA (7.4 mL, 53.2 mmol) was added, followed by compound **12** (8.9 g) in DCM (50 mL). The cooling bath was removed, and the reaction mixture was stirred at ambient temperature overnight. After completion of the reaction, the solvent was removed under reduced pressure, and the residue was taken up in ethyl acetate (200 mL). The solution was transferred to a separatory funnel and washed sequentially with water (100 mL) and brine (2 × 100 mL), then dried over anhydrous $Na_2SO_4$. The drying agent was filtered off, and the filtrate was concentrated under reduced pressure to yield a crude residue, which was purified by flash column chromatography on silica gel using a linear gradient of ethyl acetate in toluene. Yield 13.9 g (89%). Colorless foam.

Alternatively, ester **14** (4.7 g, 10 mmol) was dissolved in DCM (30 mL), and the solution was cooled in an ice bath. Trifluoroacetic acid (30 mL) was added dropwise, and the reaction mixture was stirred at 0 °C for 1 h, followed by an additional 1 h at room temperature. TLC chromatography indicated complete consumption of the starting material. The volatile components were removed under reduced pressure to afford a light brown oil, which was dissolved in DCM (50 mL) and washed in a separatory funnel with 1 M aqueous solution of $Na_2CO_3$ (100 mL). The aqueous layer was separated and extracted with DCM (3 × 50 mL). The combined organic layers were washed with brine (2 × 100 mL) and dried over anhydrous $Na_2SO_4$. The drying agent was removed by filtration, and the filtrate was concentrated to dryness under reduced pressure. The resulting light brown oil (3.6 g) was obtained and used directly in the next reaction step without additional purification.

Benzyl 2-(*S*)-[(benzyloxycarbonyl)amino]-6-aminohexanoate (3.6 g), acid **11** (2.3 g, 9.7 mmol), and PyBroP (5.4 g, 11.6 mmol) were dissolved in DMF (30 mL), and DIPEA (6.8 mL, 38.8 mmol) was added in one portion. The reaction mixture was allowed to stir at room temperature overnight. After completion of the reaction, water (100 mL) was added, the mixture was cooled in an ice bath, and the aqueous phase was acidified with 1 M HCl to pH 2–3 (as indicated by pH indicator paper). The reaction mixture was transferred to separatory funnel and extracted with ethyl acetate (4 × 50 mL). The combined organic extracts were washed with water (50 mL) and brine (2 × 50 mL), then dried over anhydrous $Na_2SO_4$. The drying agent was removed by filtration, and the filtrate was concentrated under reduced pressure. The resulting crude residue was purified by flash column chromatography on silica gel using a linear gradient of ethyl acetate in toluene to afford the product. Yield 3.0 g (51%). Colorless foam. $R_f$ = 0.54 (ethyl acetate : toluene 20 : 80). $[\alpha]_D^{20}$ = -6.7 (c = 2 ; $CHCl_3$ ).

$^1$H NMR (600 MHz, $CDCl_3$; T = 25 °C): 1.36 m + 1.43 m (2H, >$CH_2$), 1.51 s (9H, 3x -$CH_3$, *t*-Bu), 1.58 m + 1.64 m (2H, >$CH_2$), 1.70 m + 1.88 m (2H, >$CH_2$), 3.35 m (2H, -$CH_2$-N), 4.43 ddd (1H, *J* = 8.3, 8.1 and 5.0 Hz, >CH-N), 5.02 d, *J* = 12.2 Hz and 5.08 d, *J* = 12.2 Hz (2H, -$CH_2$-O), 5.15 d, *J* = 12.0 Hz and 5.19 d, *J* = 12.0 Hz (2H, -$CH_2$-O), 5.42 d (1H, *J* = 8.1 Hz, >N-H), 6.40 br t (1H, *J* = 5.0 Hz, >N-H), 6.94 td (1H, *J* = 7.6 and 1.5 Hz, Ar-CH=), 7.30 – 7.35 m (10 H, 2x $C_6H_5$), 7.40 ddd (1H, *J* = 8.4, 7.6 and 1.5 Hz, Ar-CH=), 7.41 dd (1H, *J* = 7.6 and 1.5 Hz, Ar-CH=), 8.34 dd (1H, *J* = 8.4 and 1.0 Hz, Ar-CH=). $^{13}$C NMR (150.9 MHz, $CDCl_3$; T = 25 °C): 22.47 (>$CH_2$), 28.30 (3x -$CH_3$, *t*-Bu), 28.48 (>$CH_2$), 32.47 (>$CH_2$), 39.54 (-$CH_2$-N), 53.47 (>CH-N), 67.01 (-$CH_2$-O), 67.20 (-$CH_2$-O), 80.16 (>C<, *t*-Bu), 119.76 (Ar-CH=, $C_6H_5$), 119.84 (Ar-C=, $C_6H_5$), 121.24 (Ar-CH=, $C_6H_5$), 126.60 (Ar-CH=, $C_6H_5$),128.00 (2x Ar-CH=, $C_6H_5$), 128.17 (2x Ar-CH=, $C_6H_5$), 128.29 (Ar-CH=, $C_6H_5$), 128.48 (2x Ar-CH=, $C_6H_5$), 128.51 (Ar-CH=, $C_6H_5$), 128.60 (2x Ar-CH=, $C_6H_5$), 132.28 (Ar-CH=, $C_6H_5$),135.13 (Ar-C=, $C_6H_5$), 136.04 (Ar-C=, $C_6H_5$), 140.16 (Ar-C=, $C_6H_5$), 153.07 (N-CO-O), 156.03 (-CO-O), 169.00 (N-CO-), 172.12 (-COOH).

IR ($CHCl_3$, $\nu_{max}$ $cm^{-1}$) 3321 m (NH); 2983 m, 2866 m, 1476 w, 1393 m, 1369 m ($CH_3$); 2942 m ($CH_2$); 1742 vs (C=O) ester; 1721 vs (C=O) carbamates; 1647 s (C=O) amide; 1518 vs (amide II); 3092 w, 3069 w, 3029 w, 1600 m, 1587 m, 1455 m, 1267 m, 698 m (benzene rings); 1160 s (($C(CH_3)_3$); 1049 m (C-O). HRMS ESI calcd for $C_{33}H_{40}O_7N_3$ $[M+H]^+$ 590.28608; found: 590.28561.

2-(*S*) - Amino-6-[[2-(*tert*-butoxycarbonyl)amino]benzoylamino]hexanoic acid **17**

Ester **16** (13.9 g, 23.6 mmol) was dissolved in methanol (200 mL), and the solution was vigorously stirred. 1 g of 10% Pd/C was carefully added, and the reaction mixture was allowed to react under atmosphere of hydrogen at pressure 15 psi for a period 18 h. After completion of reaction, TLC analysis revealed full disappearance of the starting material. The catalyst was removed by filtration through a pad of Celite, which was washed subsequently with methanol (250 mL). The combined filtrates were concentrated under reduced pressure to afford a yellow oil. The crude product was dissolved in the minimal amount of hot methanol, and diethyl ether was added dropwise until crystallization occurred. The precipitated solid was collected by filtration on a Büchner funnel, washed with diethyl ether (200 mL), and dried overnight in a desiccator under high vacuum. Yield 6.9 g (80%). White solid, m. p. 244 – 246 °C. $R_f$ = 0.78 (isopropanol : conc. $NH_4OH$ : water 7 : 1 : 2). $[\alpha]_D^{20}$ = +7.1 (c = 2 ; $CH_3COOH$ ).

$^1$H NMR (600 MHz, DMSO; T = 25 °C): 1.39 m (2H, >$CH_2$), 1.44 s (9H, 3x -$CH_3$, *t*-Bu), 1.51 m (2H, >$CH_2$), 1.62 m + 1.74 m (2H, >$CH_2$), 3.15 dd (1H, *J* = 7.0 and 5.2 Hz, >CH-N), 3.23 m (2H, -$CH_2$-N), 7.04 ddd (1H, *J* = 7.8, 7.2 and 1.0 Hz, Ar-CH=), 7.45 ddd (1H, *J* = 8.5, 7.2 and 1.5 Hz, Ar-CH=), 7.59 br (2H, -$NH_2$), 7.79 dd (1H, *J* = 7.8 and 1.5 Hz, Ar-CH=), 8.20 dd (1H, *J* = 8.5 and 1.0 Hz, Ar-CH=), 8.88 t (1H, *J* = 5.6 Hz, >N-H), 10.79 br (1H, >NH).$^{13}$C NMR (150.9 MHz, DMSO; T = 25 °C): 22.78 (>$CH_2$), 28.14 (3x -$CH_3$, *t*-Bu), 28.66 (>$CH_2$), 30.86 (>$CH_2$), 39.19 (>CH-N), 54.30 (>CH-N), 80.88 (>C<, *t*-Bu), 118.53 (Ar-CH=, $C_6H_5$), 119.60 (Ar-C=, $C_6H_5$), 121.46 (Ar-CH=, $C_6H_5$), 128.42 (Ar-CH=, $C_6H_5$), 132.10 (Ar-CH=, $C_6H_5$), 139.78 (Ar-C=, $C_6H_5$), 152.29 (N-CO-O), 168.35 (N-CO), 170.15 (-COOH). IR (KBr, $\nu_{max}$ $cm^{-1}$) 2978 s, 2869 m, 1477 m, 1393 s, 1368 s ($CH_3$); 2935 m ($CH_2$); 1599 s, 1589 s, 1447 s ($CO_2^-$) carboxylate;1729 s, 1706 s (C=O) carbamate; 1632 vs (C=O) amide; 1521 s (amide II); 3062 w, 1599 s, 1589 s, 1477 m, 1302 m, 1280 m, 1048 m, 753 m (benzene ring); 1160 vs (($C(CH_3)_3$). HRMS ESI calcd for $C_{18}H_{28}O_5N_3$ $[M+H]^+$ 366.20235; found: 366.20218.

2-(*S*)-[(9H-Fluoren-9-ylmethoxycarbonyl)amino]-6-[[2-(*tert*-butoxycarbonyl)amino]benzoylamino]hexanoate **18**

Acid **17** (5.5 g, 15 mmol) and $NaHCO_3$ (5 g, 60 mmol) were suspended in water (100 mL). The turbid reaction mixture was cooled to 0 °C in an ice bath, and a solution of Fmoc-OSu (5.1 g, 15 mmol) in dioxane (100 mL) was added dropwise with stirring. The mixture was allowed to stir at 0 °C for 1 h and then at room temperature overnight. The reaction mixture was cooled to 0 °C and acidified to pH 1–2 by addition of 1 M HCl (monitored by pH indicator paper). The mixture was transferred to a separatory funnel, 200 ml of water was added, and extracted with ethyl acetate (4 × 100 mL). The combined organic layers were washed with water (100 mL) and brine (2 × 100 mL), dried over anhydrous $Na_2SO_4$, filtered, and concentrated in vacuo. The crude residue was subjected flash chromatography on silica using a linear gradient of ethyl acetate in toluene. Resulting product was dissolved in 100 ml dioxane, frozen and lyophilized. Yield 4.2 g (48%). $R_f$

= 0.56 (ethyl acetate : methanol : acetone : water 6 : 2 :1 : 0.5; TLC plate was pretreated by gaseous ammonia). Colorless lyophilizate. $[\alpha]_D^{20}$ = -5.5 (c = 2 ; DMF ).

$^1$H NMR (600 MHz, DMSO; T = 25 °C): 1.39 m (2H, >$CH_2$), 1.46 s (9H, 3x -$CH_3$, *t*-Bu), 1.53 m (2H, >$CH_2$), 1.64 m + 1.75 m (2H, >$CH_2$), 3.24 m (2H, -$CH_2$-N), 3.94 ddd (1H, *J* = 9.6, 8.0 and 5.5 Hz, >CH-N), 4.22 m (1H, -CH<, Fmoc), 4.27 m (2H, -$CH_2$-O), 7.04 td (1H, *J* = 7.6 and 1.0 Hz, Ar-CH=), 7.32 m (2H, 2x Ar-CH=, Fmoc), 7.40 m (2H, 2x Ar-CH=, Fmoc), 7.45 ddd (1H, *J* = 8.4, 7.6 and 1.0 Hz, Ar-CH=), 7.62 d (1H, *J* = 8.0 Hz, >N-H), 7.70 dd (1H, *J* = 7.6 and 1.0 Hz, Ar-CH=), 7.72 m (2H, 2x Ar-CH=, Fmoc), 7.89 m (2H, 2x Ar-CH=, Fmoc), 8.20 m (1H, *J* = 8.4 and 1.0 Hz, Ar-CH=), 8.72 t (1H, *J* = 5.6 Hz, >N-H), 10.72 br s (1H, >N-H), 12.56 br (1H, □COOH).$^{13}$C NMR (150.9 MHz, DMSO; T = 25 °C): 23.33 (>$CH_2$), 28.13 (3x -$CH_3$, *t*-Bu), 28.55 (>$CH_2$), 30.63 (>$CH_2$), 39.14 (>$CH_2$), 46.82 (>CH-, Fmoc), 53.93 (>CH-N), 65.75 (□$CH_2$-O), 79.86 (>C<, *t*-Bu), 118.61 (Ar-CH=), 119.66 (Ar-C=), 120.26 and 120.27 (2x Ar-CH=, Fmoc), 121.47 (Ar-CH=),125.43 and 125.45 (2x Ar-CH=, Fmoc), 127.21 and 127.22 (2x Ar-CH=, Fmoc), 127.79 (2x Ar-CH=, Fmoc), 128.24 (Ar-C=), 132.82 (Ar-CH=), 139.74 (Ar-C=), 140.88 and 140.89 (2x Ar >C=, Fmoc), 143.96 and 144.02 (2x Ar >C=, Fmoc), 152.28 (O-CO-N), 156.33 (O-CO-N), 168.36 (-CO-N), 174.14 (-COOH). IR (KBr, $\nu_{max}$ $cm^{-1}$) 3304 m (NH); 2977 m, 2865 m, 1478 m, 1393 m, 1368 m ($CH_3$); 2977 m ($CH_2$); 1725 vs (C=O) acid; 1707 vs (C=O) carbamate; 1641 (C=O) amide; 1521 vs (amide II); 3067 m, 3041 m, 1599 m, 1587 m, 1449 s, 1115 m, 1025 m, 758 m, 742 m (rings); 1160 vs (($C(CH_3)_3$). HRMS ESI calcd for $C_{33}H_{37}O_7N_3Na$ $[M+Na]^+$ 610.25237; found: 610.25220.

2-(*S*)-[(9H-Fluoren-9-ylmethoxycarbonyl)amino]-3-(4-hydroxy-3-nitrophenyl)hexanoic acid **20**

2-(*S*)-Amino-3-(4-hydroxy-3-nitrophenyl)hexanoic acid **19** (4.5 g, 20 mmol) and $NaHCO_3$ (6.7 g, 80 mmol) were dissolved with using sonication in water (100 mL). The reaction mixture was cooled to 0 °C in an ice bath, and a solution of Fmoc-OSu (6.7 g, 20 mmol) in dioxane (100 mL) was added dropwise with stirring. The mixture was allowed to stirr at 0 °C for 1 h and then at room temperature overnight. The reaction mixture was cooled to 0 °C and acidified to pH 1–2 by addition of 1 M HCl (monitored by pH indicator paper). The mixture was transferred to a separatory funnel, 200 ml of water was added, and extracted with ethyl acetate (4 × 100 mL). The combined organic layers were washed with water (100 mL) and brine (2 × 100 mL), dried over anhydrous $Na_2SO_4$, filtered, and concentrated in vacuo. The crude solid was recrystallized from ethyl acetate/petroleum ether to afford the product as a solid. Yield 8.4 g (93%). Yellow solid, m. p. 144 - 145°C. $R_f$ = 0.51

(ethyl acetate : methanol : acetone : water 6 : 2 :1 : 0.5; TLC plate was pretreated by gaseous ammonia). $[\alpha]_D^{20}$ = -19.2 (c = 2 ; DMF ).

$^1$H NMR (600 MHz, DMSO; T = 25 °C): 2.85 dd, *J* = 13.8 + 10.6 Hz and 3.08 dd *J* = 13.8 + 4.4 Hz (2H, >$CH_2$), 4.16 m (1H, -CH<, Fmoc), 4.17 ddd (1H, *J* = 10.6, 4.4 and 8.4 Hz, >CH-N), 4.20 m (2H, -$CH_2$-O), 7.05 d (1H, *J* = 2.2 Hz, Ar-CH=), 7.28 m (2H, 2x Ar-CH=, Fmoc), 7.40 m (2H, 2x Ar-CH=, Fmoc), 7.62 m (2H, 2x Ar-CH=, Fmoc), 7.76 d (1H, *J* = 8.4 Hz, >N-H), 7.84 d (1H, *J* = 2.2 Hz, Ar-CH=), 7.87 m (2H, 2x Ar-CH=, Fmoc), 10.79 br (1H, -OH), 12.80 br (1H, -COOH). $^{13}$C NMR (150.9 MHz, DMSO; T = 25 °C): 35.21 (>$CH_2$), 46.74 (>CH-, Fmoc), 55.41 (>CH-N), 65.83 (-$CH_2$-O), 119.18 (Ar-CH=, $C_6H_5$), 120.26 and 120.28 (2x Ar-CH=, Fmoc), 125.33 and 125.38 (2x Ar-CH=, Fmoc), 125.64 (Ar-CH=, $C_6H_5$), 127.18 and 127.20 (2x Ar-CH=, Fmoc), 127.80 (2x Ar-CH=, Fmoc), 129.37 (Ar >C=, $C_6H_5$), 136.33 (Ar >C=, $C_6H_5$), 136.41 (Ar >C=, $C_6H_5$), 140.84 and 140.87 (2x Ar >C=, Fmoc), 143.86 and 143.93 (2x Ar >C=, Fmoc), 151.05 (Ar >C=, $C_6H_5$), 156.14 (O-CO-N), 173.20 (-COOH). IR (KBr, $\nu_{max}$ $cm^{-1}$) 3418 s (OH); 3322 s (NH); 3067 w, 3042 w, 1630 m, 1490 m, 1451 m, 1428 m, 760 m, 741 m (rings); 1538 vs, 1325 s ($NO_2$). HRMS ESI calcd for $C_{24}H_{19}O_7N_2$ $[M-H]^-$ 447.11977; found: 447.11953.

#### 1.2: Synthesis of pseudo-dipeptide precursors I-IV

Syntheses of pseudo-dipeptide block precursors that were used for preparations of pseudopeptides **2-8** are shown in **Schemes 3-6**. The syntheses of phosphinate intermediates **II**, **III** and **IV** were based on the reaction of Cbz-Nleψ[PO(OH)H] **28**, the synthesis of which is shown in **Scheme 2**. Analytical HPLC profiles of the key synthetic intermediates **28, 37, 42, 47** and **51** are shown in **Figure S2**. Analytical HPLC profiles of the final pseudo-dipeptide precursors **I-IV** are shown in **Figure S3**.

#### 1.2.1: Synthesis of Cbz-Nleψ[PO(OH)

We developed two independent synthetic routes based on established protocols for the preparation of α-aminophosphinic acids using the "Ciba-Geigy synthon" **21** (Ref.[1]), both of which are applicable on a multigram scale. In pathway A, compound **21** was refluxed in benzene with pentanal and triphenylmethyl-amine or diphenylmethyl-amine in a three-component aza-Pudovik reaction using a Dean–Stark apparatus. Water was removed azeotropically, and the imine formed *in situ* subsequently reacted with phosphinic acid ester **21**. Fully protected intermediates **22** and **23** were obtained and isolated in moderate yields. In pathway B, the starting synthon **21** was converted into α-hydroxyphosphinate **24** *via* the Abramov reaction by heating with valeraldehyde in the presence of a base. Introduction of the azido group using the Mitsunobu reaction provided compound **25**, which was smoothly reduced to the amino intermediate **26**. The acid-labile protecting groups of intermediates **22**, **23**, and **26** were removed by heating in concentrated hydrochloric acid to provide Nleψ[PO(OH)H] as the corresponding zwitterion **27**, which, upon protection of the amino group with a benzyloxycarbonyl group, yielded the desired compound **28**.

**1.2.1.1: Synthetic protocols for compounds in Scheme 2**

*O*-Ethyl-[(*R*,*S*)-1-(triphenylmethylamino)pentyl]diethoxymethylphosphinate **22**

A 500 mL flask equipped with a Dean–Stark apparatus was charged with triphenylmethylamine (13.0 g, 0.05 mol), valeraldehyde (5.3 mL, 0.05 mol), and *O*-ethyl diethoxymethylphosphinate (**21**, Ciba–Geigy reagent; 9.8 g, 0.05 mol) in benzene (150 mL). The reaction mixture was heated to reflux and maintained under these conditions until the theoretical amount of water had been collected in the Dean–Stark trap. Benzene was removed under reduced pressure, and the resulting residue was purified by column chromatography on silica gel using a linear gradient of ethyl acetate in cyclohexane. Yield 15.4 g (59%) as a colorless oil. Rf = 0.46 (ethyl acetate/toluene, 1:1). Two stereoisomers were observed in a ratio of 53:47. Only the signals of the major isomer are reported below.

$^{1}$H NMR (600 MHz, $CDCl_3$; T = 25 °C): 0.69 t (*J* = 7.1 Hz, $-CH_3$), 0.87 m ($>CH_2$), 0.92 m + 1.25 m ($>CH_2$), 0.93 m + 1.085 m ($>CH_2$), 1.19 t (*J* = 7.1 Hz, $-CH_3$), 1.26 t (*J* = 7.1 Hz, $-CH_3$), 1.34 t (*J* = 7.1 Hz, $-CH_3$), 2.63 dd (*J* = 8.8 and 3.4 Hz, >NH), 3.08 m (N-CH-P), 3.55 m + 3.90 m ($-CH_2$-O), 3.77 m + 3.88 m ($-CH_2$-O), 4.21 m ($-CH_2$-O), 4.90 d (*J* = 9.2 Hz, P-CH<O), 7.20 m (3x Ar-H), 7.27 m (6x Ar-H), 7.57 m (6x Ar-H).$^{13}$C NMR (150.9 MHz, $CDCl_3$; T = 25 °C): 13.77 ($-CH_3$), 15.22 + 15.20 (2x $-CH_3$), 16.68 d (*J* = 5.2 Hz, $-CH_3$), 22.79 ($-CH_2$-), 28.45 d (*J* = 1.7 Hz, $-CH_2$-), 29.56 d (*J* = 2.4 Hz, $-CH_2$-), 52.05 d (*J* = 100.0 Hz, N-CH-P), 61.47 d (*J* = 7.3 Hz, $-CH_2$-O), 64.70 d (*J* = 8.9 Hz, $-CH_2$-O), 65.76 d (*J* = 8.1 Hz, $-CH_2$-O), 71.40 d (*J* = 5.4 Hz, >C<), 100.81 d (*J* = 136.5 Hz, O-CH-P). 126.41 (2x Ar-CH=), 127.68 (4x Ar-CH=), 129.01 (4x Ar-CH=), 146.44 (2x Ar-C=).$^{31}$P NMR (202.4 MHz, $CDCl_3$; T = 25 °C): 44.61, 43.18. IR $\nu_{max}/cm^{-1}$ ($CHCl_3$): 3351 w (NH); 3086 w, 3062 w, 1596 m, 1490 m, 1448 m, 1003 m, 707 vs, 641 m (rings); 1249 m (P=O); 1035 s, 958 m (P-O-C); 1058 vs (COCOC); 2982 s, 1392 m, 1375 m 1456 m ($CH_3$); 2931 m, 1467 m ($CH_2$). HRMS (ESI) [$M+H^+$] calc. for $C_{31}H_{43}O_4NP$: 524.29242; found: 524.29252.

*O*-Ethyl-[(*R*,*S*)-1-(diphenylmethylamino)pentyl)]diethoxymethylphosphinate **23**

Phosphinate **23** was prepared according to the same procedure used for compound **22** by the reaction of diphenylmethylamine (18.3 g, 0.10 mol), valeraldehyde (10.6 mL, 0.10 mol), and *O*-ethyl-diethoxymethylphosphinate **21** (19.6 g, 0.10 mol) in benzene (200 mL). Upon completion of the reaction, the reaction mixture was worked up and purified as described for compound **13**. Yield 31 g (69%). Colorless oil. $R_f$ = 0.37 (ethyl acetate/toluene 1:1). Two diastereoisomers observed in the ratio 53 : 47. Only the signals of the major isomer are given below.

$^1$H NMR (600 MHz, $CDCl_3$; T = 25 °C): 0.86 t (*J* = 7.1 Hz, -$CH_3$), 1.15 t (*J* = 7.1 Hz, -$CH_3$), 1.21 m (>$CH_2$), 1.22 t (*J* = 7.1 Hz, -$CH_3$), 1.32 m + 1.51 m, >$CH_2$), 1.34 t (*J* = 7.1 Hz, -$CH_3$), 1.58 m + 1.84 m (>$CH_2$), 2.96 m (N-CH-P), 3.55 m + 3.84 m (-$CH_2$-O), 3.67 m + 3.89 m (-$CH_2$-O), 4.19 – 4.29 m (-$CH_2$-O), 4.89 – 4.91 m (P-CH<O), 5.22 s (N-CH<), 7.20 m (2x Ar-H), 7.28 m (4x Ar-H), 7.42 m (4x Ar-H). $^{13}$C NMR (150.9 MHz, $CDCl_3$; T = 25 °C): 13.90 (-$CH_3$), 15.15 + 15.18 (2x -$CH_3$), 16.76 d (*J* = 5.1 Hz, -$CH_3$), 22.62 (-$CH_2$-), 28.16 d (*J* = 7.6 Hz, -$CH_2$-), 28.75 d (*J* = 2.9 Hz, -$CH_2$-), 51.90 d (*J* = 94.5 Hz, N-CH-P), 61.78 d (*J* = 7.5 Hz, -$CH_2$-O), 64.58 d (*J* = 7.6 Hz, -$CH_2$-O), 64.82 d (*J* = 8.3 Hz, -$CH_2$-O), 64.99 d (*J* = 9.5 Hz, -$CH_2$-O), 100.82 d (*J* = 132.8 Hz, O>CH-P). 126.99 – 127.13 (2x Ar-CH=), 127.38 – 128.34 (8x Ar-CH=), 143.63 (2x Ar-C=).$^{31}$P NMR (202.4 MHz, $CDCl_3$; T = 25 °C): 44.32, 44.04. IR $v_{max}$/cm$^{-1}$ ($CHCl_3$): 3342 w (NH); 3087 w, 3065w, 1599 w, 1493 m, 1453 m, 707 vs, 836 w, 704 s, 620 w (rings); 1058 vs, 1033 s, 957 m (P-O-C); 1058 vs (COCOC); 2982 s, 2963 m, 1392 m, 1370 w ($CH_3$); 2931 m, 2864 m, 1479 w ($CH_2$). HRMS (ESI) [M+Na$^+$] calc. for $C_{25}H_{38}O_4NNaP$: 470.24307; found: 470.24288

*O*-Ethyl-((*R*,*S*)-1-hydroxypentyl)diethoxymethylphosphinate **24**

A 250 mL flask was charged with DIPEA (26.1 mL, 0.15 mol), valeraldehyde (16 mL, 0.15 mol), and *O*-ethyl diethoxymethylphosphinate **21** (29.4 g, 0.15 mol), and the mixture was heated at 95 °C with stirring for 48 h. After cooling to room temperature, the reaction mixture was taken up in ethyl acetate (250 mL) and transferred to a separatory funnel. The organic layer was washed with 0.1 M $KHSO_4$ (250 mL). The upper organic phase was separated, washed with brine (2 × 150 mL), and dried over anhydrous $Na_2SO_4$. The drying agent was removed by filtration, and the filtrate was concentrated under reduced pressure. The resulting yellow residue was purified by column chromatography on silica gel using a linear gradient of ethyl acetate/methanol/acetone (6:2:1) in ethyl acetate. Yield 25.5 g (60%). Light yellow oil. $R_f$ = 0,54 and 0,46 ($CHCl_3$/$CH_3OH$ 95:5). Mixture of four stereoisomers. NMR data of the two major isomers are given below.

$^1$H NMR (600 MHz, $CDCl_3$; T = 25 °C): 0.90 t (*J* = 7.1 Hz, -$CH_3$), 1.24 – 1.26 t (*J* = 7.1 Hz, 2x -$CH_3$), 1.335 t (*J* = 7.1 Hz, -$CH_3$), 1.35 (>$CH_2$), 1.38 – 1.61 m (>$CH_2$), 1.65 – 1.79 m (>$CH_2$), 2.90 br (-OH), 3.70 m + 3.85 m (2x -$CH_2$-O), 3.93 m + 4.02 m (O-CH-P), 4.19 m + 4.24 m (-$CH_2$-O), 4.86 d (J = 6.8 Hz, O>CH-P).$^{13}$C NMR (150.9 MHz, $CDCl_3$; T = 25 °C): 13.88 (-$CH_3$), 15.11 - 15.20 (2x -$CH_3$), 16.44 – 16.70 (-$CH_3$), 22.22 + 22.34 (-$CH_2$-), 27.86 d + 27.94 d (*J* = 11.6 Hz, -$CH_2$-), 29.40 + 29.87 (-$CH_2$-),62.22 d, *J* = 7.6 Hz + 62.30 d, *J* = 7.5 Hz (-$CH_2$-O), 65.58 d, *J* = 8.8

Hz + 65.80 d, $J$ = 8.1 Hz -$CH_2$-O), 65.74 d, $J$ = 9.8 Hz + 66.18 d, $J$ = 7.9 Hz (-$CH_2$-O), 65.80 d , $J$ = 8.1 Hz, 68.06 d, $J$ = 101.7 Hz + 68.37 d , $J$ = 99.9 Hz (O-CH-P), 100.80 d + 100.82 d ($J$ = 135.2 Hz, O>CH-P). $^{31}$P NMR (202.4 MHz, $CDCl_3$; T = 25 °C): 39.93, 39.91. IR $\nu_{max}$/cm$^{-1}$ ($CHCl_3$): 3564 w (OH); 2984 s, 2963 m, 2963 m, 1393 m, 1370 w ($CH_3$); 2932 m, 1444 m ($CH_2$); 1233 m (P=O); 1032 s, 959 m (P-O-C); 1058 vs (COCOC); HRMS (ESI) [M+Na$^+$] calc. for $C_{12}H_{27}O_5NaP$: 305.14883; found: 305.14899.

*O*-Ethyl-((*R*,*S*)-1-azidopentyl)diethoxymethylphosphinate **25**

To a three-necked 1 L flask equipped with gas inlet and outlet tubes were added alcohol **24** (25.2 g, 89.2 mmol) and triphenylphosphine (28.0 g, 107 mmol) dissolved in dry THF (400 mL). The reaction mixture was cooled in an ice bath, and under an inert atmosphere of argon, DIAD (21.6 g, 107 mmol) was added dropwise over 10 min, followed by the addition of DPPA (29.4 g, 107 mmol). The mixture was allowed to react at 0 °C for 1 h and then at room temperature overnight. Upon completion of the reaction, the solvent was removed under reduced pressure, and the residue was purified by column chromatography on silica gel using a linear gradient of ethyl acetate in cyclohexane. Yield 23.9 g (87%). Light yellow oil. $R_f$ = 0,48 (ethyl acetate/toluene 1:1). Two diastereoisomers observed in the ratio ca 2 : 1. Only the signals of the major isomer are given below.

$^1$H NMR (600 MHz, $CDCl_3$; T = 25 °C): 0.91 t ($J$ = 7.1 Hz, -$CH_3$), 1.25 t ($J$ = 7.1 Hz, 2x -$CH_3$), 1.34 t ($J$ = 7.1 Hz, -$CH_3$), 1.36 m (>$CH_2$), 1.39 m + 1.58 m (>$CH_2$), 3.47 m (N-CH-P), 3.74 m + 3.86 m (2x -$CH_2$-O), 4.26 m (-$CH_2$-O), 4.88 d ($J$ = 9.5 Hz, O-CH-P). $^{13}$C NMR (150.9 MHz, $CDCl_3$; T = 25 °C): 13.77 (-$CH_3$), 15.10 + 15.17 (2x -$CH_3$), 16.63 d ($J$ = 4.9 Hz, -$CH_3$), 22.07 (-$CH_2$-), 26.76 (-$CH_2$-), 28.87 d ($J$ = 11.1 Hz, -$CH_2$-), 57.62 d ($J$ = 99.0 Hz, N-CH-P), 62.65 d ($J$ = 7.2 Hz, -$CH_2$-O), 66.28 d ($J$ = 8.2 Hz, -$CH_2$-O), 100.08 d ($J$ = 142.9 Hz, O>CH-P).$^{31}$P NMR (202.4 MHz, $CDCl_3$; T = 25 °C): 38.95, 38.34. IR $\nu_{max}$/cm$^{-1}$ ($CHCl_3$): 2102 vs ($N_3$); 1238 m (P=O); 1034 s, 962 m (P-O-C); 1059 vs (COCOC); 2984 s, 2963 m, 1393 w, 1370 w ($CH_3$); 2931 m, 2875 w ($CH_2$). HRMS (ESI) [M+H$^+$] calc. for $C_{12}H_{27}N_3O_4P$: 308.17337; found: 308.17318.

*O*-Ethyl-((*R*,*S*)-1-aminopentyl)diethoxymethylphosphinate **26**

Azide **25** (22.1 g, 71.9 mmol) was dissolved in methanol (200 mL). To the resulting solution, 10% Pd/C (500 mg) was added, and the vigorously stirred mixture was subjected to hydrogenolysis under a hydrogen atmosphere at 15 psi. After 12 hours, the catalyst was removed by filtration through Celite, and the filter cake was washed with methanol (250 mL). Evaporation of the combined filtrates under reduced pressure afforded a yellow residue, which was purified by column chromatography using a linear gradient of ethyl acetate, methanol, and acetone (6:2:1) in ethyl acetate. Yield 17.7 g (87%). Light yellow oil. $R_f$ = 0,77 (ethylacetate/methanol/acetone/water 6 : 1 : 1 : 0,5). IR $v_{max}/cm^{-1}$ ($CHCl_3$): 3378 w, 3312 w (NH); 2983 vs, 2932 s, 1392 m, 1372 m ($CH_3$); 1242 s, 1231 (P=O); 1036 vs, 958 s (P-O-C); 1057 vs (COCOC). HRMS (ESI) [$M+H^+$] calc. for $C_{12}H_{29}NO_4P$: 282.18287; found: 282.18272.

((*R*,*S*) -1- aminopentyl)phosphinic acid **27**

Phosphinate **22** (15.0 g, 28.7 mmol) or phosphinate **23** (30.7 g, 68.6 mmol) were refluxed overnight in a 500 mL round-bottom flask equipped with a Dimmroth condenser in 35% hydrochloric acid (150 mL). After cooling to room temperature, the reaction mixture was transferred to a separatory funnel and extracted with chloroform (4 × 50 mL). The aqueous layer was separated and evaporated to dryness under reduced pressure. The residue was subsequently co-distilled with toluene (3 × 150 mL). The resulting solid was dissolved in ethanol (150 mL), and 1,2-epoxypropane was added dropwise with stirring and cooling in an ice bath until neutral pH was reached (monitored by pH indicator paper). The precipitated solid was collected by filtration on a fritted glass funnel, washed with ice-cold ethanol (50 mL) followed by diethyl ether (50 mL), and dried under vacuum.

To phosphinate **25**, 35% hydrochloric acid (150 mL) was carefully added dropwise from an addition funnel with stirring and cooling in an ice bath. The addition funnel was then replaced with a Dimmroth condenser, and the mixture was refluxed overnight. After completion of the reaction, the mixture was evaporated to dryness under reduced pressure. The subsequent isolation procedure was identical to that described for the hydrolysis of phosphinates **22** and **23**. Yield 3.3 g (77% from **22**), 7.4 g (71% from **23**), and 7.0 g (77% from **24**). The physico-chemical and spectral characteristics of the compound are fully consistent with the data previously reported by us (2).

((*R*,*S*) -1-{[(Benzyloxy)carbonyl]amino}pentyl)phosphinic acid **28**

In a 1 L round-bottom flask, phosphinic acid **27** (14.2 g, 94 mmol) and $Na_2CO_3$ (20.0 g, 188 mmol) were dissolved in water (200 mL). The reaction mixture was cooled in an ice bath, and a solution of Cbz-OSu (23.4 g, 94 mmol) in dioxane (150 mL) was added dropwise over 30 min with stirring. The mixture was maintained at 0 °C for 2 h and then allowed to react at rt overnight. Then, the solvent volume was reduced to approximately one-half by evaporation under reduced pressure. The concentrated mixture was cooled again in an ice bath and, under vigorous stirring, acidified with 0.1 M hydrochloric acid to pH 1 (monitored by pH indicator paper). Water (400 mL) was added, and the resulting suspension was collected by vacuum filtration using a Büchner funnel. The precipitate was rinsed with cold water until the filtrate reached neutral pH. The obtained crystals were dried overnight in a desiccator over $P_2O_5$. Final purification was achieved by recrystallization from ethyl acetate. Yield 22.1 (82%). The physico-chemical and spectral characteristics of the compound are fully consistent with the data previously reported by us (2).

#### 1.2.2.: Synthesis of precursor I

In an attempt to prepare pseudodipeptide **I** (**Scheme 3**), corresponding to the phosphonodepsipeptide Fmoc-Nleψ[PO(OH)O]Lys(Boc)-OH, the previously prepared monobenzyl ester **34** was used as the starting material (2). Intermediate **37** was obtained through three synthetic steps involving alkylation, hydrogenolysis, and acylation,. Subsequently, **37** was subjected to coupling with protected L- Lys(Boc)OH **30** using a range of common coupling reagents, including BOP, PyBroP, HCTU, and HATU; however, no desired product **38** was obtained. A standard Mitsunobu reaction between phosphonic acid **34** and D-Lys(Boc)OH **33** was also investigated, but no coupling product **38** was detected in the reaction mixture. The failure to prepare the asymmetrical ester **38** is most likely attributable to adamantyl steric hindrance. Therefore, compound **34** was instead coupled with alcohol **33**, which underwent smooth esterification under Mitsunobu reaction conditions. Subsequent hydrogenolysis removal of both benzyl and benzyloxycarbonyl (Cbz) protecting groups, followed by reaction of the α-amino functionality with Fmoc-OSu afforded pseudo-precursor **I** bearing a free P(O)OH moiety. With this protecting group arrangement, intermediate is fully compatible with the solid-phase peptide synthesis protocol, as we have previously reported[2].

#### 1.2.2.1: Synthetic protocols for compounds in Scheme 3

Benzyl 6-(*tert.* butoxycarbonylamino)-2-(*S*)-hydroxyhexanoate **30**

Compound **30** was prepared according to the reported protocol (3), starting from acid **29** (9.9 g; 40 mmol), $NaNO_2$ (5.5 g; 80 mmol), benzyl bromide (4.8 mL; 40 mmol), and $K_2CO_3$ (5.5 g; 40 mmol). Yield 4.7 g (35%). Light yellow oil. $R_f$ = 0.71 (ethyl acetate/toluen 50 : 50). $[\alpha]_D^{20}$ = - 14.7 (c = 2 ; $CHCl_3$).

$^1$H NMR (600 MHz, $CDCl_3$; T = 25 °C): 1.36 m + 1.46 m (2H, >$CH_2$), 1.43 s (9H, 3x -$CH_3$, *t*-Bu), 1.43 s (9H, 3x -$CH_3$, *t*-Bu), 1.46 m (2H, >$CH_2$), 1.66 m + 1.81 m (2H, >$CH_2$), 3.06 m (2H, -$CH_2$-N), 4.21 dd (1H, *J* = 7.5 and 4.1 Hz, >CH-O), 4.54 br (1H, >N-H), 5.18 d, *J* = 12.0 Hz and 5.21 d, *J* = 12.0 Hz (2H, -$CH_2$-O), 7.33 – 7.38 m (5x Ar-H, $C_6H_5$).$^{13}$C NMR (150.9 MHz, $CDCl_3$; T = 25 °C): 21.92 (>$CH_2$), 28.37 (3x -$CH_3$, *t*-Bu), 29.64 (>$CH_2$), 33.81 (>$CH_2$), 40.28 (-$CH_2$-N), 67.27 (-$CH_2$-O), 70.28 (>CH-O), 79.10 (>C<, *t*-Bu), 128.34 (2x Ar-CH=, $C_6H_5$), 128.55 (Ar-CH=, $C_6H_5$), 128.62 (2x Ar-CH=, $C_6H_5$), 135.14 (Ar-C=, $C_6H_5$), 155.96 (N-CO-O), 174.96 (-COOH).

IR $v_{max}/cm^{-1}$ ($CHCl_3$): 3543 w, 3456 w (OH); 2981 m, 2935 m, 1393 m, 1368 m ($CH_3$); 2951 m, 2867 w ($CH_2$); 1730 vs (C=O) ester; 1710 vs (C=O) carbamate; 1508 s (amide II); 1170 vs ($(CH_3)_3$); 3092 w, 3069 w, 1588 w, 1456 m, 698 m (benzene ring). HRMS (ESI): $[M+Na]^+$ for $C_{18}H_{27}NO_5Na$ calc: 360.17814; found: 360.17809.

6-(*tert.* Butoxycarbonylamino)-2-(*R*)-aminohexanoic acid **32**

Compound **32** was prepared according to the reported protocol by Picha et al.[4] for the *S*-isomer, starting from *D*-lysine·HCl **31** (18.26 g; 50 mmol), $Cu(OAc)_2 \cdot H_2O$ (10 g; 25 mmol), 2 M NaOH (100 mL), di-*tert*-butyl dicarbonate (24 g; 55 mmol), and 8-hydroxyquinoline (9.45 g ; 65 mmol). Yield 11.4 g (93%). Colorless solid, m. p. 245°C (decay). $R_f$ = 0,68 (isopropyl alcohol/conc. $NH_4OH$/water 6 : 2 : 1). $[\alpha]_D^{20}$ = - 15.1 (c = 2 ; glacial acetic acid).

$^1$H NMR (600 MHz, DMSO; T = 25 °C): 1.28 m + 1.32 m (2x >$CH_2$), 1.37 s (3x -$CH_3$ (*t*-Bu)), 1.53 m + 1.68 m (>$CH_2$), 2.87 (N-$CH_2$-), 3.06 (dd, *J* = 7.0 and 4.8 Hz, N-CH<), 6.75 t (*J* = 5.5 Hz, >NH). $^{13}$C NMR (150.9 MHz, DMSO; T = 25 °C): 22.72 (>$CH_2$), 28.48 (3x $CH_3$, *t*-Bu), 29.44 and 31.02 (2x >$CH_2$), 39.97 (N-$CH_2$-), 54.34 (N-CH<), 77.54 (>C<, *t*-Bu), 155.73 (O-CO-N), 169.80 (-COOH). IR $v_{max}/cm^{-1}$ (KBr): 3375 s, (NH); 2979 s, 2866 s, 1392 s, 1366 s ($CH_3$); 2937 s ($CH_2$); 1690 vs (C=O) amide, 1527 vs (amide II); 1583 vs, 1411 vs (C=O) carboxylate; 1180 vs ($(CH_3)_3$). HRMS (ESI): $[M+Na]^+$ pro $C_{11}H_{22}N_2O_4Na$ calc: 269.14718; found: 269.14722.

Benzyl 6-(*tert.* butoxycarbonylamino)-2-(*R*)-hydroxyhexanoate **33**

Compound **33** was prepared according to the reported protocol[3] for the *S*-isomer, starting from acid **32** (11.4 g; 46.3 mmol), $NaNO_2$ (6.4 g; 92.6 mmol), benzyl bromide (5.5 mL; 46.3 mmol), and $K_2CO_3$ (6.4 g; 46.3 mmol). Yield 6 g (38%). Light yellow oil. $R_f$ = 0.71 (ethyl acetate/toluene 50 : 50). $[\alpha]_D^{20}$ = + 11.1 (c = 2 ; $CHCl_3$).

$^1$H NMR (600 MHz, $CDCl_3$; T = 25 °C): 1.37 m + 1.48 m (-$CH_2$-), 1.45 s (9H, *t*-Bu), 1.48 m (-$CH_2$-), 1.67 m + 1.83 m (-$CH_2$-), 3.09 m (N-$CH_2$-), 4.23 dd (*J* = 7.5 and 4.1 Hz, >CH-O), 4.53 br (>NH), 5.21 d + 5.28 d (*J* = 12.1 Hz, O-$CH_2$-), 7.36 – 7.40 m (5H, $C_6H_5$). $^{13}$C NMR (150.9 MHz, $CDCl_3$; T = 25 °C): 21.93 (-$CH_2$-), 28.39 (3x -$CH_3$, *t*-Bu), 29.66 + 33.83 (2x >$CH_2$), 40.36 (-$CH_2$-N), 70.27 (>CH-O), 79.16 (>C<, *t*-Bu), 128.38 (2x Ar-CH=), 128.59 (Ar-CH=), 128.65 (2x Ar-CH=), 135.12 (Ar-C=), 155.98 (O-CO-N), 175.00 (O-CO-).IR $v_{max}/cm^{-1}$ ($CHCl_3$): 3541 w, 3456 w (OH); 2980 m, 2935 m, 1393 m, 1368 m ($CH_3$); 2951 m, 2867 w ($CH_2$); 1730 vs (C=O) ester; 1710 vs (C=O) carbamate; 1508 s (amide II); 1170 vs ($(CH_3)_3$); 3092 w, 3069 w, 1588 w, 1456 m, 698 m (benzene ring). HRMS (ESI): $[M+Na]^+$ for $C_{18}H_{27}NO_5Na$ calc: 360.17814; found: 360.17823.

Adamantan-1-yl benzyl [(*R*,*S*)-1-(benzyloxycarbonylamino)-pentyl]phosphonate **35**

In a 250 mL two-necked round-bottom flask equipped with a reflux condenser, benzyl hydrogen [(*R*,*S*)-1-(benzyloxycarbonylamino)-pentyl]phosphonate **34** (7.8 g, 20 mmol) and 1-bromoadamantane (5.2 g, 24 mmol) were dissolved in chloroform (100 mL). Silver(I) oxide (9.3 g, 40 mmol) was divided into five equal portions, which were added successively to the reaction mixture at 30 min intervals while maintaining the temperature at 60 °C. After complete addition of $Ag_2O$, the mixture was stirred and heated at 60 °C overnight. Upon completion of the reaction, the solvent was removed under reduced pressure. The resulting black–grey residue was suspended in diethyl ether (100 mL) and filtered through a pad of Celite. The filter cake was washed with diethyl ether (300 mL), and evaporation of the combined filtrates under reduced pressure afforded a pale-yellow residue. The crude product was purified by column chromatography using a linear gradient of ethyl acetate in cyclohexane to give the product as a colorless oil, which solidified upon standing in fridge at 5°C overnight. Yield 9.8 g (93%). Colorless solid, 78 – 80 °C. $R_f$ = 0.46 (ethyl acetate/toluene 20 : 80). A mixture of stereoisomers was obtained in an approximately 1:1 ratio. Specific structure assignment of signals to each stereoisomer was not possible.

$^1$H NMR (600 MHz, $CDCl_3$; T = 25 °C): 0.87 t (-$CH_3$), 1.26 + 1.33 m (-$CH_2$-), 1.33 m + 1.42 m (-$CH_2$-), 1.49 + 1.84 (-$CH_2$-), 1.58 m + 1.64 m (3x -$CH_2$-, Ada), 2.01 m + 2.05 m (3x -$CH_2$-, Ada), 2.13 m (3x -CH<, Ada), 3.98 – 4.03 m (N-CH-P), 4.90 dd (*J* = 10.5 and 1.8 Hz) + 4.84 dd (*J* = 10.4 and 2.3 Hz, >NH), 5.00 – 5.07 m (-$CH_2$-O), 5.12 d + 5.04 d (*J* = 12.0 Hz, -$CH_2$-O), 7.29 – 7.36 m (2x $C_6H_5$).$^{13}$C NMR (150.9 MHz, $CDCl_3$; T = 25 °C): 13.87 (-$CH_3$), 22.23 (-$CH_2$-), 27.90 d, *J* = 12.5 Hz + 27.96 d, *J* = 12.6 Hz (-$CH_2$-), 29.67 d, *J* = 2.7 Hz + 29.82 d, *J* = 2.7 Hz (-$CH_2$-), 31.12 + 31.13 (3x >CH-, Ada), 35.63 + 36.04 (3x -$CH_2$-, Ada), 43.77 d, *J* = 3.7 Hz + 43.90 d, *J* = 3.7 Hz (3x -$CH_2$-, Ada), 48.46 d, *J* = 157.9 Hz + 49.07 d, *J* = 159.1 Hz (-$CH_2$-P), 66.95 (-$CH_2$-O), 67.49 + 67.53 (-$CH_2$-O), 83.35 d, *J* = 10.1 Hz + 83.38 d, *J* = 10.3 Hz (N-CH-P), 127.85 – 128.47 (10x Ar-CH=), 136.38 (Ar-C=), 136.47 d, *J* = 6.8 Hz + 136.50 d, *J* = 6.3 Hz (Ar-C=), 156.03 d, *J* = 5.6 Hz + 126.06 d, *J* = 5.6 Hz (O-CO-N). IR $\nu_{max}$/cm$^{-1}$ (KBr): 3234 m (NH); 2954 m, 2864 m ($CH_3$); 2850 m, 1455 m ($CH_2$); 2914 s (C-H), 1721 vs (C=O) carbamate; 1545 m (amide II); 1222 s (P=O); 1062-984 s (P-O-C); 3087 w, 3063 w, 3034 w, 1497 m, 730 m, 695 m (rings). HRMS (ESI): $[M+Na]^+$ for $C_{30}H_{40}NO_5NaP$ calc: 548.25363; found: 548.25293.

Adamantan-1-yl hydrogen [(*R*,*S*)-1-(aminopentyl)]phosphonate **36**

Ester **35** (9.5 g, 18.1 mmol) was dissolved in methanol (200 mL). To the resulting solution, 10% Pd/C (500 mg) was added, and the vigorously stirred mixture was subjected to hydrogenolysis under a hydrogen atmosphere at 15 psi. After 18 hours, TLC analysis revealed full consumption of starting compound. The catalyst was removed by filtration through Celite, and the filter cake was washed with methanol (250 mL). The filtrate was evaporated under reduced pressure to afford yellowish foam. The residue was dissolved in the minimum volume of hot acetonitrile, and the solution was left in a freezer at −20 °C overnight. The precipitated crystals were collected by centrifugation for 10 min at 9000 rpm. The supernatant was removed, and the solid was dried under high vacuum overnight. Yield 4.4 g (80%). Colorless solid, 220-222 °C. $R_f$ = 0.83 (isopropyl alcohol/conc. $NH_4OH$/water 6 : 2 : 1).

$^1$H NMR (600 MHz, DMSO; T = 25 °C): 0.86 t (*J* = 7 Hz, -$CH_3$), 1.25 m, 1.39 m and 1.73 m (3x >$CH_2$), 1.55 m (3x -$CH_2$-, Ada), 1.96 m (3x -$CH_2$-, Ada), 2.04 m (3x >CH-, Ada), 2.56 ddd (*J* = 12.3, 7.7 and 6.0 Hz, N-CH-P).$^{13}$C NMR (150.9 MHz, DMSO; T = 25 °C): 14.02 ($CH_3$), 22.27 (>$CH_2$), 28.30 d (*J* = 7.1 Hz, (>$CH_2$), 29.43 (>$CH_2$), 30.50 (3x >CH-, Ada), 36.05 (3x >$CH_2$, Ada), 44.00 d (*J* = 3.5 Hz, 3x >$CH_2$, Ada), 50.04 d (*J* = 141.7 Hz, N-CH-P). 74.45 (>C<, Ada). IR $v_{max}$/cm$^{-1}$ (KBr): 2955 s, 2874 s ($CH_3$); 2938 s, 2853 s, 1456 m ($CH_2$); 2910 vs (CH); 1203 s (P=O); 1051 vs ($POO^-$); 994 s, 982 s, 982 s, 961 m (P-O-C); 1623 w, 1539 w ($NH_3^+$). HRMS (ESI): [M-H] for $C_{15}H_{27}NO_3P$ calc: 300.17340; found: 300.17330.

Adamantan-1-yl hydrogen [(*R*,*S*)-1-(benzyloxycarbonylamino)-pentyl]phosphonate **37**

In a 250 mL round-bottom flask, phosphinic acid **36** (4.7 g, 15.6 mmol) and $Na_2CO_3$ (1.7 g, 15.6 mmol) were dissolved in water (50 mL). The turbid reaction mixture was cooled in an ice bath, and a solution of Cbz-OSu (3.9 g, 15.6 mmol) in dioxane (50 mL) was added dropwise over 30 min with stirring. The mixture was maintained at 0 °C for 1 h and then allowed to react at room temperature overnight. The mixture was cooled again in an ice bath and, under vigorous stirring, acidified with 0.1 M hydrochloric acid to pH 1–2 (monitored by pH indicator paper). Water (100 mL) was added, and the resulting suspension was extracted with chloroform (4 × 200 mL). The combined organic layers were washed with brine (200 mL) and dried over $Na_2SO_4$. The drying agent was removed by filtration, and the filtrate was evaporated under reduced pressure to dryness. Final purification was achieved by recrystallization of the resulting precipitate from ethyl acetate to give compound **37**. Yield 5 g (74%). Colorless solid, 162-165 °C. $R_f$ = 0.86 ($CHCl_3$/MeOH/AcOH 6 : 2 : 1).

$^1$H NMR (600 MHz, DMSO; T = 25 °C): 0.84 t (*J* = 7 Hz, -$CH_3$), 1.20 m + 1.29 m (>$CH_2$), 1.35 s, 1.21 m + 1.33 m (>$CH_2$), 1.47 m + 1.66 m (>$CH_2$), 1.53 m + 1.56 m (3x >$CH_2$, Ada), 1.93 m + 1.97 m (3x >$CH_2$, Ada), 2.07 m (3x >CH-, Ada), 3.60 m (N-CH-P), 5.00 d + 5.08 d (*J* = 12.7 Hz, -$CH_2$-O), 7.175 d (*J* = 9.7 Hz, >NH), 7.29 – 7.36 m (5H, $C_6H_5$).$^{13}$C NMR (150.9 MHz, DMSO; T = 25 °C): 14.03 ($CH_3$), 21.78, 28.02 d (*J* = 13.3 Hz) and 28.66 (3x >$CH_2$), 30.57 (3x >CH-, Ada), 35.61

(3x >$CH_2$, Ada), 43.56 (3x >$CH_2$, Ada), 49.34 d ($J$ = 159.5 Hz, N-CH-P), 65.40 (-$CH_2$-O), 79.57 d ($J$ = 9.5 Hz, >C<, Ada), 127.75 (2x Ar-CH=), 127.88 (Ar-CH=), 128.45 (2x Ar -CH=), 137.55 (Ar >C=), 156.34 (O-CO-N).$^{31}$P NMR (202.4 MHz, $CDCl_3$; T = 25 °C): 20.37. IR $v_{max}$/cm$^{-1}$ (KBr): 3277 m (NH); 2953 m, 2872 m ($CH_3$); 2930 m, 2855 m ($CH_2$); 2913 s (CH); 1712 vs (C=O) carbamate; 1549 s (amide II); 1209 m (P=O); 1062 s, 1041 m, 1030 m, 1023 m, 975 m (P-O-C); 3089 w, 3063 w, 3033 w, 1454 m, 959 m, 757 m, 703 m (ring). HRMS (ESI): [M-H] for $C_{23}H_{33}NO_5P$ calc: 434.21018; found: 434.20982.

(2*S*)-2-{[(*R*,*S*)-1-(9H-Fluoren-9-ylmethoxycarbonylamino)pentyl(hydroxy)phosphoryl]oxy}-6-[(*tert*-butoxycarbonyl)amino]hexanoic acid **(I)**

In a 100 mL round-bottom flask, benzyl hydrogen[(*R*,*S*)-1-(benzyloxycarbonylamino)pentyl]phosphonate **34** (3.6 g, 9.2 mmol), alcohol **33** (2.1 g, 6.2 mmol), $PPh_3$ (2.4 g, 9.2 mmol), and TEA (5.1 mL, 36.8 mmol) were dissolved in anhydrous THF (40 mL). With stirring at room temperature under an argon atmosphere, DIAD (1.9 g, 9.2 mmol) was added and the reaction mixture was allowed to react overnight. After completion of reaction, tetrahydrofuran was removed under reduced pressure and the residue was purified by column chromatography using a linear gradient of ethyl acetate in cyclohexane as the eluent. A total of 6.6 g of product was obtained; it was partially contaminated with diisopropyl hydrazinedicarboxylate and was used in the next synthetic step without further purification. Benzyl (2*S*)-2-{[(*R*,*S*)-1-(benzyloxycarbonylamino)pentyl(benzyloxy)phosphoryl]oxy}-6-[(*tert*-butoxycarbonyl)amino]hexanoate was dissolved in methanol (100 mL). To the solution, 10% Pd/C (250 mg) was added and the mixture was subjected to hydrogenolysis in an autoclave under hydrogen at a pressure of 15 psi with vigorous stirring. After 12 h, the catalyst was removed by filtration through Celite and the filter cake was washed with methanol (100 mL). Evaporation of the filtrate under reduced pressure afforded a pale-yellow oil, which was dissolved in dioxane (100 mL), frozen, and lyophilized. A total of 4 g of the intermediate was obtained and used in the subsequent synthetic step without additional purification. (2*S*)-2-{[(*R*,*S*)-1-Aminopentyl(hydroxy)phosphoryl]oxy}-6-[(*tert*-butoxycarbonyl)amino]hexanoic acid together with $NaHCO_3$ (1 g, 12.4 mmol) was placed in a 100 mL round-bottom flask and dissolved in a mixture of water (20 mL) and dioxane (20 mL). The reaction mixture was cooled in an ice bath and a solution of Fmoc-OSu (2.5 g, 7.44 mmol) in dioxane (20 mL) was added dropwise over 30 min with stirring. The mixture was maintained at 0 °C for 1 h and then allowed to react at room temperature overnight. The mixture was cooled again in an ice bath and, under vigorous stirring, acidified with 0.1 M hydrochloric acid until pH 2 was reached (monitored by pH indicator paper).

Water (50 mL) was then added and the reaction mixture was extracted in a separatory funnel with ethyl acetate (4 × 30 mL). The combined organic layers were washed with brine (2 × 50 mL) and dried over anhydrous sodium sulfate. The drying agent was removed by filtration and the filtrate was evaporated under reduced pressure. The residue was purified by column chromatography using a linear gradient of the mixture chloroform/methanol/acetic acid (6:2:0.5) in chloroform as the eluent. The resulting pale-yellow oil was lyophilized twice from dioxane to afford of the final product. Yield 1.8 g (47% over three steps). Colorless lyophilizate. $R_f$ = 0.51 ($CHCl_3$/MeOH/AcOH 6 : 2 : 1). Mixture of stereoisomers was obtained. Only the signals of the major isomer and/or the center of signal group for given atom are presented below.

$^{1}$H NMR (600 MHz, DMSO; T = 25 °C): 0.84 m (-$CH_3$), 1.23 m and 1.30 m (>$CH_2$), 1.33 m (>$CH_2$), 1.35 s (3x -$CH_3$, *t*-Bu), 1.53 m (>$CH_2$), 1.72 m (2x >$CH_2$), 2.86 m (N-$CH_2$-), 3.76 m (N-CH--P),4.21 m (>CH-), 4.24 m (-$CH_2$-O), 4.62 m (O-CH<), 6.74 m (>NH), 7.32 m (2x Ar-CH=, Fmoc), 7.42 m (2x Ar-CH=, Fmoc), 7.46 m (>NH), 7.73 (2x Ar-CH=, Fmoc), 7.89 (2x Ar-CH=, Fmoc). $^{13}$C NMR (150.9 MHz, DMSO; T = 25 °C): 14.07 (-$CH_3$), 21.85 (>$CH_2$), 26.75 (>$CH_2$), 28.00 (>$CH_2$), 28.89 (>$CH_2$), 28.47 (3x -$CH_3$, t-Bu), 32.56 (>$CH_2$), 32.82 (>$CH_2$), 39.91 (N-$CH_2$-), 46.88 (>CH-), 48.38 ((N-CH-P), 66.11 m (-$CH_2$-O), 77.55 (>C<, *t*-Bu), 120.33 (2x Ar-CH=, Fmoc), 125.65 (2x Ar-CH=, Fmoc), 127.28 (2x Ar-CH=, Fmoc), 127.87 (2x Ar-CH=, Fmoc), 140.32 (2x Ar-C=, Fmoc), 143.88 (2x Ar-C=, Fmoc), 155.77 (O-CO-N), 156.40 (O-CO-N), 172.22 (-CO-N). $^{31}$P NMR (202.4 MHz, DMSO; T = 25 °C): 23.77. IR $v_{max}/cm^{-1}$ (KBr): 3334 m (NH); 2957 s, 2872 m, 1395 m, 1368 m ($CH_3$); 2934 s ($CH_2$); 1715 vs + vbr (C=O) acid, carbamates; 1210 s (P=O); 1043 s - 994 s (P-O-C); 1166 vs (($CH_3$)$_3$); 3099 w, 3067 w, 3020 w, 1478 m, 1451 s, 1100 m, 759 s, 740 s (fluorene). HRMS (ESI): [M-H] for $C_{31}H_{42}N_2O_9P$ calc: 617.26334; found: 617.26328.

#### 1.2.3: Synthesis of precursor II

Phosphinic building block **II** was synthesized by conventional N→C synthetic strategy as described in **Scheme 4**. The P1′ fragment, designed to mimic lysine, was prepared analogously in the form of an acrylate ester by acylation of the starting Meldrum's acid **39**. Intermediate **40** was subsequently converted to the corresponding benzyl acrylate by sequential reaction with benzyl alcohol and paraformaldehyde *via* the Knoevenagel condensation. Direct alkylation of dibenzyl malonate with *N*-Boc-4-bromobutylamine afforded an inseparable mixture of mono- and disubstituted malonate derivatives. In the subsequent step, the silylated protected phosphinic acid **28** readily underwent a phospho-Michael addition to the acrylate, furnishing intermediate **42**. Although the phosphinic acid functionality was successfully protected with adamantyl ester, hydrogenolytic removal of the Cbz and benzyl protecting groups from compound **43** always resulted in complete deprotection of the adamantyl group under all reaction conditions reported in the literature[5]. Therefore, for the preparation of the pseudodipeptide building block **II**, compound **42** was subjected directly to hydrogenolysis, and the free α-amino group of the phosphinonorleucine moiety was subsequently protected by acylation with Fmoc-OSu. The resulting pseudo-dipeptide precurssor **II** was fully compatible with the solid-phase peptide synthesis protocol and can serve as a practical alternative to compound **44**, which would otherwise be the preferred substrate for peptide synthesis, as previously demonstrated for closely related analogues[6].

**1.2.3.1: Synthetic protocols for compounds in Scheme 4**

2,2-Dimethyl-5-[4-(*tert*-butoxycarbonylamino)butyl]-1,3-dioxane-4,6-dione **40**

In a 500 mL round-bottom flask, Boc-GABA-OH (16.9 g, 83.2 mmol), Meldrum's acid **39** (12 g, 83.2 mmol), and DMAP (15.2 g, 124.8 mmol) were dissolved in 150 mL of DCM. The reaction mixture was cooled in an ice bath and DIC (14.3 mL, 91.5 mmol) was added dropwise over 10 minutes with stirring. The mixture was allowed to react for 1 h at 0 °C and then for an additional 12 h at room temperature. The precipitated DIU was filtered off, filtrate was transferred to a separatory funnel and washed with 200 mL of 1 M $KHSO_4$, followed by brine (2 × 200 mL), and dried over $Na_2SO_4$. After filtration of the drying agent, the filtrate was concentrated under reduced pressure. The crude intermediate was partially purified by crystallization from ethyl acetate and used directly in the subsequent synthetic step. A portion of the intermediate, 2,2-dimethyl-5-[1-oxo-4-(*tert*-butoxycarbonylamino)butyl]-1,3-dioxane-4,6-dione (8.7 g), was dissolved in 150 mL of DCM, cooled in an ice bath, and glacial acetic acid (9 mL, 158.4 mmol) was added with stirring. Sodium borohydride (2.6 g, 68.6 mmol) was then added portionwise. The mixture was maintained at 0 °C for 2 h and subsequently allowed to react overnight at room temperature. The resulting suspension was cooled again in an ice bath and carefully quenched by the slow addition of 100 mL of water with vigorous stirring. In a separatory funnel, the upper aqueous layer was separated and extracted with DCM (2 × 100 mL). The combined organic layers were washed with brine (2 × 100 mL) and dried over anhydrous $Na_2SO_4$. After filtration of the drying agent, the filtrate was concentrated under reduced pressure. The residue was purified by column chromatography using a linear gradient of ethyl acetate in cyclohexane, followed by crystallization from an ethyl acetate/petroleum ether mixture. Yield 5.7 g (22 % over two steps). Colorless solid, m. p. 114 - 115°C. $R_f$ = 0,59 (ethyl acetate/toluene 50 : 50).

$^1$H NMR (600 MHz, $CDCl_3$; T = 25 °C): δ = 1.43 (s, 9H, *t*-Bu), 1.48 (m, 2H, $CH_2$), 1.54 (m, 2H, $CH_2$), 1.75 (q, 3H, *J* = 0.7 Hz, $CH_3$), 1.78 (q, 3H, *J* = 0.7 Hz, $CH_3$), 2.10 (m, 2H, $CH_2$), 3.13 (t, 2H,

*J* = 6.8 Hz, $CH_2$–N), 3.52 (m, 1H, *J* = 5.1 Hz, CH), 4.55 (br, 1H, NH) ppm; $^{13}$C NMR (150.9 MHz, $CDCl_3$; T = 25 °C): δ = 23.62 ($CH_2$), 25.95 ($CH_2$), 26.80 ($CH_3$), 28.39 (3 × $CH_3$ (*t*-Bu)), 28.43 ($CH_3$), 29.86 ($CH_2$), 40.08 ($CH_2$–N), 79.15 (>C< (*t*-Bu)), 104.87 (O–C–O), 155.97 (O–CO–N), 165.41 (2 × O–CO) ppm. IR $v_{max}$/$cm^{-1}$ (KBr): 3386 m, 3351 m (NH); 2980 m, 1393 m, 1368 m ($CH_3$); 2936 m, 2869 w ($CH_2$); 1798 s, 1750 vs (C=O) ester; 1686 vs (C=O) carbamate; 1525 s (amide II); 1180 s (($C(CH_3)_3$). HRMS (ESI): $[M+H]^+$ for $C_{15}H_{26}NO_6$ calc: 316,17546; found: 316,17581.

Benzyl 6-[(*tert*-butoxycarbonyl)amino]-2-methylenehexanoate **41**

Dioxane **40** (5.6 g, 17.8 mmol) and benzyl alcohol (3.9 g, 35.6 mmol) were dissolved in 40 mL of toluene in a 100 mL round-bottom flask. The reaction mixture was heated at 80 °C overnight. After completion of the reaction, the solvent was removed under reduced pressure. The residue was purified by column chromatography using a linear gradient of ethyl acetate in cyclohexane to afford 6.1 g of a yellow oil, which was used directly in the subsequent synthetic step. A mixture of benzyl 6-[(*tert*-butoxycarbonyl)amino]-2-carboxyhexanoate, paraformaldehyde (0.5 g, 16.7 mmol), piperidine (0.165 mL, 1.67 mmol), and pyridine (10 mL) was placed in a 50 mL round-bottom flask equipped with a reflux condenser and heated at 80 °C overnight. After completion of the reaction, the volatile materials were removed under reduced pressure. The oily residue was dissolved in 100 mL of ethyl acetate and transferred to a separatory funnel. The solution was washed with 100 mL of 1 M $KHSO_4$. The aqueous phase was separated and extracted with ethyl acetate (2 × 100 mL). The combined organic layers were washed with brine (100 mL) and dried over anhydrous $Na_2SO_4$. The drying agent was removed by filtration and the filtrate was concentrated under reduced pressure. The residue was purified by column chromatography using a linear gradient of ethyl acetate in cyclohexane to afford 3.9 g (66%) of the desired product.

Alternatively, a mixture of dioxane **40** (3.15 g, 10 mmol), Eschenmoser's salt (4.8 g, 26 mmol), and benzyl alcohol (15 mL) was heated at 65 °C overnight in a 50 mL round-bottom flask. After cooling to room temperature, the reaction mixture was dissolved in 100 mL of ethyl acetate, transferred to a separatory funnel and washed with brine (100 mL). The aqueous phase was separated and extracted with ethyl acetate (100 mL). The combined organic layers were dried over anhydrous $Na_2SO_4$, filtered, and concentrated under reduced pressure. The residue was purified by column chromatography using a linear gradient of ethyl acetate in cyclohexane to give 1.5 g (45%) of the product. Slightly yellow oil. $R_f$ = 0,60 (ethyl acetate/toluen 20 : 80).

$^1$H NMR (600 MHz, $CDCl_3$; T = 25 °C): δ = 1.44 (s, 9H, *t*-Bu), 1.50 (m, 4H, 2 × $CH_2$), 2.34 (m, 2H, $CH_2$), 3.12 (br, 2H, $CH_2$–N), 4.51 (br, 1H, NH), 5.19 (s, 2H, $CH_2$–O), 5.56 (q, 1H, *J* = 1.5 Hz) a 6.20 (dt, 1H, $J$ = 1.5 and 0.6 Hz), (=$CH_2$), 7.33 - 7.39 (m, 5H, $C_6H_5$) ppm; $^{13}$C NMR (150.9 MHz, $CDCl_3$; T = 25 °C): δ = 25.60 ($CH_2$), 28.40 (3 × $CH_3$ (*t*-Bu)), 29.59 ($CH_2$), 31.48 ($CH_2$), 40.29 ($CH_2$–N), 66.38 (–$CH_2$–O), 79.05 (>C< (*t*-Bu)), 125.24 (=$CH_2$), 128.02 (2 × m–Ar–CH), 128.14 (p–ArH), 128.53 (2 × o–Ar–CH), 136.05 (i–Ar–C), 140.28 (C=), 155.94 (O–CO–N), 166.95 (O–CO) ppm. IR $v_{max}$/$cm^{-1}$ ($CHCl_3$): 3455 w (NH), 2981 w, 2866 w, 1393 w, 1368 w ($CH_3$); 2938 w ($CH_2$); 1630 m (C=$CH_2$); 1711 vvs (C=O) ester + carbamate; 1507 s (amide II); 1164 s (($CH_3$)$_3$); 3093 w, 3068 w, 1083 w, 698 m (phenyl). HRMS(ESI): $[M+Na]^+$ for $C_{19}H_{27}NNaO_4$ calc: 356.18323; found: 356.18327.

Benzyl (*R*,*S*)-2-[((*R*,*S*)-1-{[(benzyloxy)carbonyl]amino}pentyl)(hydroxyphosphoryl)methyl]-6-[(tert-butoxycarbonyl)amino]hexanoate **42**

In a 250 mL three-neck round-bottom flask equipped with a gas inlet and outlet, phosphinic acid **28** (5.2 g, 18 mmol) was suspended in HMDS (18.9 mL, 90 mmol). The suspension was stirred and heated to 110 °C under an argon atmosphere for 1 h. After this period, the mixture became completely homogeneous, giving a clear solution. The temperature was then lowered to 90 °C, and a solution of the unsaturated ester **41** (7.8 g, 23.4 mmol) dissolved in toluene (5 mL) was added dropwise to the resulting clear solution. The reaction mixture was subsequently maintained at 110 °C for an additional 3 h. After completion of the reaction, the mixture was cooled in an ice bath and quenched by the addition of 96% ethanol (100 mL). Volatile materials were removed under reduced pressure using a rotary evaporator. The residue was dissolved in ethyl acetate (50 mL) and washed with 1 M HCl (50 mL) in a separatory funnel. The aqueous phase was separated and extracted with ethyl acetate (3 × 50 mL). The combined organic layers were washed with brine (3 × 50 mL) and dried over anhydrous sodium sulfate. After filtration drying agent, the solvent was removed under reduced pressure. The crude residue was purified by column chromatography using a linear gradient of chloroform/methanol/acetic acid (6:2:0.5) in chloroform as the eluent to afford required product. Yield 7.8 g (80 %). Light yellow oil. $R_f$ = 0,59 (ethyl acetate/acetone/methanol/water 6 : 1 : 1 : 0.5).

$^1$H NMR (600 MHz, $CDCl_3$; T = 25 °C): δ = 0.86 (t, 3H, $J$ = 7 Hz, $CH_3$), 1.43 (s, 9H, *t*-Bu), 1.18 –1.35, 1.56 – 1.82 (m, 12H, 6 × $CH_2$), 3.91, 2.99 a 2.83 (m, 5H, N–CH, N–$CH_2$ a P–$CH_2$), 5.05 – 5.14 (m, 4H, 2 × O–$CH_2$), 5.46 (m, 2H, 2 × NH), 7.28 – 7.34 (m, 10 H, 2 × $C_6H_5$). 9.54 (br, 2H, P–OH) ppm; $^{13}$C NMR (150.9 MHz, $CDCl_3$; T = 25 °C): δ = 13.89 ($CH_3$), 22.19 – 29.33 (7 × $CH_2$), 23.38 (3 × $CH_3$ (*t*-Bu)), 38.66 – 49.89 (N–CH, N–$CH_2$ a P–$CH_2$), 66.70 – 67.10 (2 × O–$CH_2$), 127.85 – 128.49 (10 × Ar–CH=), 135.79 – 136.36 (2 × Ar>C=), 156.39 – 156.43 (2 × O–CO–NH),

174.45 – 174.50 (O–CO) ppm; $^{31}P$ NMR (202.4 MHz, $CDCl_3$; T = 25 °C): δ = 55.28, 54.99, 53.86, 53.12 ppm. $R_f$ = 0,59 ($SiO_2$; ethylacetate/acetone/methanol/water 6 : 1 : 1 : 0.5). IR $\nu_{max}/cm^{-1}$ ($CHCl_3$): 3454 w, 3430 w, 3400 w, 3306 w (NH); 2979 m, 2874 m, 1393 m, 1368 m ($CH_3$); 2934 m, 2863 w ($CH_2$); 1711vs + vbr (C=O) ester, carbamates; 1508 s (amide II); 1243 s (P=O); 966 m (P-O-H); 1170 s ($(CH_3)_3$); 3091 w, 3068 w, 1587 w, 1456 m, 1028 w, 698 m (rings). HRMS (ESI) $[M+Na]^+$ for $C_{32}H_{47}N_2NaO_8P$ calc: 641.29622; found: 641.29614.

Benzyl (*R,S*)-2-[((*R,S*)-1-{[(benzyloxy)carbonyl]amino}pentyl)(adamantan-1-yloxyphosphoryl)methyl]-6-[(*tert*-butoxycarbonyl)amino]hexanoate **43**

Compound **43** was prepared by the reaction of **42** (4.1 g; 6.6 mmol), 1-bromoadamantane (1.6 g; 7.3 mmol) and silver(I) oxide (3.1 g; 13.3 mmol) in chloroform (40 ml) according to the procedure used for the preparation of compound **35**. Yield 2.2 g (44%). Light yellow oil. $R_f$ = 0.44 (toluene/ethyl acetate 1 : 1). Mixture of stereoisomers was obtained. Specific structure assignment of signals to each stereoisomer was not possible. Only the signal regions corresponding to each type of proton and carbon atoms are shown.

$^1H$ NMR (600 MHz, $CDCl_3$; T = 25 °C): 0.87 t (-$CH_3$), 1.42 s (9H, *t*-Bu), 1.16 - 1.42 m (6x -$CH_2$-), 1.42 s (3x -$CH_3$, *t*-Bu), 1.58 – 1.63 m (3x -$CH_2$-, Ada), 1.58 – 1.68 m (P-$CH_2$-), 2.00 m (3x -$CH_2$-, Ada), 2.12 m (3x -CH<, Ada), 2.83 – 3.00 m (>CH-), 3.90 m (P-CH-N), 4.82 d (*J* = 10 Hz, >NH), 4.87 d (*J* = 10 Hz, >NH), 5.05 – 5.20 m (2x O-$CH_2$-), 7.27 – 7.37 m (10x Ar-CH, 2x $C_6H_5$). $^{13}C$ NMR (150.9 MHz, $CDCl_3$; T = 25 °C): 13.87 – 13.92 (-$CH_3$), 22.24 – 22.32 (-$CH_2$-), 23.68 – 23.82 (-$CH_2$-), 28.38 (3x $CH_3$, *t*-Bu), 27.82 – 28.58 (-$CH_2$-), 29.17 – 30.59 (-$CH_2$-), 30.79 – 31.18 (3x -CH<, Ada), 33.20 – 33.87 (P-$CH_2$-), 35.61 – 36.18 (3x -$CH_2$-, Ada), 39.17 – 39.43 (>CH-), 40.26 (-$CH_2$-N), 41.54 – 41.89 (-$CH_2$-N), 44.22 – 44.46 (3x -$CH_2$-, Ada), 49.34 – 51.78 (P-CH-N), 66.50 - 67.36 (2x O-$CH_2$-), 79.06 (>C<, *t*-Bu), 82.97 – 83.76 (>C<, Ada), 127.79 – 129.52 (10x Ar-CH=), 135.72 – 136.43 (2x Ar-C<), 155.56 – 156.34 (2x O-CO-N), 174.43 (–O-CO-). $^{31}P$ NMR (202.4 MHz, $CDCl_3$; T = 25 °C): 49.97, 48.78, 48.55, 47.50. IR $\nu_{max}/cm^{-1}$ ($CHCl_3$): 3459 w, 3431 w (NH); 2981 m, 2875 w, 1393 m, 1368 m; 2920 s, 2859 m ($CH_2$); 1731 vs (C=O) ester; 1714 vs (C=O) carbamates; 1508 s (amide II); 1239 s (P=O); 1049 s – 984 s (P-O-C); 1170 s ($(CH_3)_3$); 3092 w, 3068 w, 1607 w, 1587 w, 1456 m, 1081 w, 698 m (rings). HRMS (ESI) $[M+Na]^+$ for $C_{42}H_{61}N_2NaO_8P$ calc: 775.40577; found: 775.40521.

(*R*,*S*)-2-[((*R*,*S*)-1-{[(9H-Fluoren-9-ylmethoxy)carbonyl]aminopentyl)(hydroxyphosphoryl)methyl]-6-[(*tert*-butoxycarbonyl)amino]hexanoic acid **II**

Phosphinic acid **42** (10.5 g; 16.9 mmol) was dissolved in methanol (200 mL). A 10% Pd/C catalyst (700 mg) was added to the solution, and the mixture was subjected to hydrogenolysis in autoclave under hydrogen atmosphere at pressure 15 psi with vigorous stirring. After 8 hours, the catalyst was removed by filtration through Celite and the pad was washed with methanol (150 mL). The combined filtrates were concentrated under reduced pressure to afford a residue, which was dissolved in a minimal amount of hot ethyl acetate. Addition of petroleum ether induced precipitation of the product, which was collected by filtration on a Büchner funnel and washed with petroleum ether (50 mL). After removal of residual solvents under high vacuum, the obtained intermediate was used in the next synthetic step without additional purification. 5.3 g of intermediate (*R*,*S*)-2-[((*R*,*S*)-1-aminopentyl)(hydroxyphosphoryl)methyl]-6-[(*tert*-butoxycarbonyl)amino]hexanoic acid together with $NaHCO_3$ (6.8 g; 80.4 mmol) were placed in a 500 mL flask and dissolved in water (100 mL). The reaction mixture was cooled in an ice bath, and a solution of Fmoc-OSu (4.5 g; 13.4 mmol) in dioxane (50 mL) was added dropwise over 30 min with stirring. The mixture was maintained at 0 °C for 1 h and then allowed to react at room temperature overnight. The reaction mixture was again cooled in an ice bath and acidified with 0.1 M HCl under vigorous stirring until pH ~ 2 was reached (monitored by indicator paper). Water (50 mL) was then added, and the mixture was extracted with ethyl acetate (4 × 50 mL). The combined organic layers were washed with brine (2 × 50 mL) and dried over anhydrous sodium sulfate. After filtration of the drying agent, the filtrate was concentrated under reduced pressure and the residue was purified by column chromatography using a linear gradient of chloroform/methanol/AcOH (6:2:0.5) in chloroform as the eluent. The resulting pale-yellow oil was lyophilized twice from dioxane to afford the product. Yield: 7.0 g (55%). Colorless lyophilizate.

Alternatively, ester **43** (1.3 g; 1.8 mmol) was dissolved in methanol (50 mL). A 10% Pd/C catalyst (300 mg) was added to the solution, and the mixture was subjected to hydrogenolysis in autoclave under hydrogen atmosphere at pressure 15 psi with vigorous stirring. After 4 hours, TLC analysis revealed disappearance of starting compound. The catalyst was removed by filtration through Celite and the pad was washed with methanol (50 mL). The combined filtrates were concentrated under reduced pressure to afford a light brown foam, which was used in the next synthetic step without additional purification. 0.8 g of intermediate together with $NaHCO_3$ (0.8 g; 6 mmol) were placed in a 50 mL flask and dissolved in water (10 mL). The turbid reaction mixture was cooled in an ice bath, and a solution of Fmoc-OSu (0.5 g; 1.5 mmol) in dioxane (5 mL) was added dropwise over

30 min with stirring. The mixture was maintained at 0 °C for 1 h and then allowed to react at room temperature overnight. The reaction mixture was again cooled in an ice bath and acidified with 0.1 M HCl under vigorous stirring until pH ~ 2 was reached (monitored by indicator paper). Water (10 mL) was then added, and the mixture was extracted with ethyl acetate (4 × 10 mL). The combined organic layers were washed with brine (2 × 10 mL) and dried over anhydrous sodium sulfate. After filtration of the drying agent, the filtrate was concentrated under reduced pressure and the residue was purified by column chromatography using a linear gradient of chloroform/methanol/AcOH (6:2:0.5) in chloroform as the eluent. The resulting pale-yellow oil was lyophilized twice from dioxane to afford the product. Yield: 0.45 g (42 %). Colorless lyophilizate.

Alternatively, to a solution of **43** (2 g, 2.66 mmol) and ammonium formate (0.67 g, 10.64 mmol) was added 10% Pd/C (250 mg). After the initial severe reaction had subsided, the mixture was stirred for 12 min. The catalyst was removed by filtration through Celite, which was subsequently washed with methanol (20 mL). The combined filtrates were concentrated to dryness, and chloroform (50 mL) was added to the residue. The precipitated excess ammonium formate was removed by filtration and the filtrate was again evaporated to dryness, affording a foamy residue. The material was dissolved in 10% aqueous $NaHCO_3$ (15 mL), and a solution of Fmoc-OSu (0.98 g, 2.9 mmol) in dioxane (10 mL) was added dropwise with stirring while cooling in an ice bath. The mixture was maintained at 0 °C for 1 h and then allowed to react at room temperature overnight. The reaction mixture was again cooled in an ice bath and acidified with 0.1 M HCl under vigorous stirring until pH ≈ 2 was reached (monitored with indicator paper). Water (50 mL) was then added, and the mixture was extracted with ethyl acetate (4 × 50 mL). The combined organic layers were washed with brine (2 × 50 mL) and dried over anhydrous sodium sulfate. After filtration of the drying agent, the filtrate was concentrated under reduced pressure and the residue was purified by column chromatography using a linear gradient of chloroform/methanol/AcOH (6:2:0.5) in chloroform as the eluent. The resulting pale-yellow oil was lyophilized twice from dioxane to afford the product. Yield: 0.6 g (38 %). Colorless lyophilizate. $R_f$ = 0.63 (ethyl acetate/acetone/methanol/water 6:1:1:0.5). $^{1}H$ NMR (600 MHz, DMSO; T = 25 °C): δ = 0.85 (t, 3H, *J* = 7 Hz, $CH_3$), 1.17 – 1.31 (2H, $CH_2$), 1.35, 1.36 (s, 9H, *t*-Bu), 1.49 – 1.71, 1.94, 2.58, 2.84, 3.59 (m, 12H, 6 × $CH_2$), 4.21 (t), 4.29 (d) (3H, CH–$CH_2$ (Fmoc)), 6.73 (t), 6.74 (t), 7.51 (d), 7.52 (d) (2H, 2 × NH), 7.31, 7.32, 7.41, 7.72, 7.73, 7.89 (m, 8 × ArH (Fmoc)) ppm; $^{13}C$ NMR (150.9 MHz, DMSO; T = 25 °C): δ =14.02, 14.03 ($CH_3$), 28.45 (3 × $CH_3$ (*t*-Bu)), 21.79 – 29.62 (2 × $CH_2$), 30.31 – 46.89 (6 × $CH_2$), 50.32 d, 50.89 d (CH–P), 77.49 (>C< (*t*-Bu)), 120.29, 125.45, 125.49, 127.19, 127.24, 127.82 (8 × Ar–CH< (Fmoc)), 140.90, 143.87, 144.08 (4 × Ar>C=, Fmoc), 155.73, 156.49, 156.52 (O–CO–NH), 176.09, 176.15 (COOH) ppm; $^{31}P$ NMR (202.4 MHz, $CDCl_3$; T = 25

°C): δ = 46.71, 46.55, 46.47, 45.98, 45.92 ppm. IR $\nu_{max}/cm^{-1}$ (KBr): 3306 m (NH); 2974 m, 2873, 1394 m, 1367 m ($CH_3$); 2934 m, 2863 m ($CH_2$); 1713 vvs + vbr (C=O) carbamates, acid; 1527 s (amide II); 1249 s (P=O); 1040 m – 966 m (P-O-H); 3067 m, 3067 m, 3042 m, 1611 w, 1451 m, 759 m, 741 s (fluorene); 1168 s ($(CH_3)_3$). HRMS (ESI): $[M-H]^-$ for $C_{32}H_{44}N_2O_8P$ calc: 615.28408; found: 615.28419.

#### 1.2.4: Synthesis of precursor III

For the preparation of the shorter analogue **III**, the Arbuzov reaction of the alkyl bromide with the corresponding P(III) trimethylsilyl phosphinite ester was conveniently employed (**Scheme 5**). Thus, the tervalent ester form derived from phosphinic acid **28** readily reacted with ethyl bromoacetate to afford the corresponding α-ethoxycarbonylphosphinic acid **47**, which was subsequently protected by acid-labile adamantyl moiety. Chemoselective saponification of the ethyl ester then furnished carboxylic acid **49** and then the C-terminus was extended with a lysine-mimicking peptoid fragment **46** using a standard peptide coupling protocol. Subsequent hydrogenolytic removal of the benzyl and Cbz protecting groups, followed by acylation with Fmoc-OSu, afforded the desired building block **III**. Notably, α-carboxyphosphinic acids bearing an adamantyl-protected phosphinic acid functionality proved to be stable toward spontaneous deprotection of the adamantyl group.

##### 1.2.4.1: Synthetic protocols for compounds in Scheme 5

Ethyl [((*R*,*S*)-1-{[(benzyloxy)carbonyl]amino}pentyl)(hydroxy)phosphoryl]acetate **47**

In a 250 mL three-necked flask equipped with gas inlet and outlet, phosphinic acid **28** (3.5 g, 12.2 mmol) was suspended in HMDS (12.8 mL, 61.3 mmol). The suspension was stirred and heated to 110 °C under an argon atmosphere for 1 hour. The resulting clear solution was then treated with ethyl 2-bromoacetate (7.3 g, 15.9 mmol), added dropwise over 10 minutes, and the reaction mixture was heated for an additional 3 hours. After completion of the reaction, the mixture was cooled in an ice bath and quenched by the addition of 96% ethanol (100 mL). Volatile materials were removed under reduced pressure, and the residue was taken up in ethyl acetate (50 mL). The solution was washed with 1 M HCl (50 mL) in a separatory funnel. The aqueous phase was separated and extracted with ethyl acetate (3 × 50 mL). The combined organic layers were washed with brine (3 × 50 mL) and dried over anhydrous sodium sulfate. After filtration, the solvent was removed under reduced pressure. The crude product was purified by column chromatography using a linear gradient of a chloroform/methanol/acetic acid mixture (6:2:0.5) in chloroform. Yield 3.6 g (80%). Colorless oil. $R_f$ = 0.50 (chloroform/methanol/AcOH 6 : 2 : 1).

$^1$H NMR (600 MHz, $CDCl_3$; T = 25 °C): 0.88 t (3H, *J* = 7.1 Hz, $-CH_3$), 1.24 t (3H, *J* = 7.2 Hz, $-CH_3$), 1.28 m and 1.35 m (2H, $-CH_2-$), 1.34 m and 1.44 m (2H, $-CH_2-$), 1.56 m and 1.89 m (2H, $-CH_2-$), 4.08 m (1H, N-CH-P), 4.15 m (4H, $-CH_2$-O and $-CH_2$-P), 5.09 d and 5.14 d (2H, *J* = 12.3 Hz, O-$CH_2$-), 5.41 br d (1H, *J* = 10 Hz, >NH), 7.29 – 7.34 m (5H, $C_6H_5$). $^{13}$C NMR (150.9 MHz, $CDCl_3$; T = 25 °C): 13.82 and 13.96 (2x $-CH_3$), 22.13 ($-CH_2-$), 27.77 (d, *J* = 2.4 Hz, $-CH_2-$), 27.87 (d, *J* = 12.0 Hz, $-CH_2-$), 50.15 (d, *J* = 110.7 Hz, N-CH-P), 61.81 ($-CH_2$-O), 67.21 ($-CH_2$-O), 127.94 (2x Ar-CH=), 128.14 (Ar-CH=), 128.47 (2x Ar-CH=), (136.18 Ar-C=), 156.41 (d, *J* = 5.8 Hz, O-CO-N), 166.02 (d, *J* = 3.9 Hz, O-CO-). $^{31}$P NMR (202.4 MHz, $CDCl_3$; T = 25 °C): 46.25. IR $v_{max}/cm^{-1}$ ($CHCl_3$): 3426 w (NH); 2983 m, 2961 m, 1381 w, 1368 m ($CH_3$); 2874 w, 2862 m, 1476 m ($CH_2$); 1722 vvs + vbr (C=O) ester + carbamate; 1508 s (amide II); 1238 s (P=O); 973 s (P-OH); 3092 w, 3068 w, 3028 w, 1455 m, 1083 m, 1028 m, 1002 m, 697 m (ring). HRMS (ESI): $[M-H]^-$ for $C_{17}H_{25}NPO_6$ calc: 370.14250; found: 370.14242.

Ethyl [((*R*,*S*)-1-{[(benzyloxy)carbonyl]amino}pentyl)(adamantan-1-yloxy)phosphoryl]acetate **48**
In a 100 mL two-necked flask equipped with a reflux condenser, phosphinic acid **47** (3.5 g, 9.4 mmol) and 1-bromoadamantane (3.1 g, 14.6 mmol) were dissolved in chloroform (60 mL). Silver oxide (6.1 g, 33.2 mmol) was divided into five equal portions and added sequentially to the reaction mixture, preheated to 60 °C, at 30-minutes intervals. After complete addition of $Ag_2O$, the mixture was stirred and allowed to react at 60 °C overnight. Upon completion of the reaction, chloroform was removed under reduced pressure. The resulting black-grey residue was suspended in diethyl ether (100 mL) and filtered through Celite. The pad was washed with diethyl ether (300 mL), and the combined filtrates were concentrated under reduced pressure to afford a paleyellow residue. The crude product was purified by column chromatography using a linear gradient of ethyl acetate in cyclohexane. Yield 3.8 g (79%). Colorless oil. $R_f$ = 0.45 (toluen/ethylacetate 50 : 50). Mixture of two stereoisomers in the ratio ca 55:45 was obtained. NMR data for the major isomer are given only.
$^1$H NMR (600 MHz, $CDCl_3$; T = 25 °C): 0.89 t (3H, *J* = 7 Hz, $-CH_3$), 1.27 t (3H, *J* = 7 Hz, $-CH_3$), 1.38 m ($-CH_2-$), 1.60 m (3x $-CH_2-$), 1.91 m ($-CH_2-$), 2.03 – 2.06 m (3x $-CH_2-$), 2.15 m (3x >CH-), 2.86 dd, *J* = 19 + 14 Hz and 3.08 t, *J* = 14 Hz (P-$CH_2$-CO), 4.13 m (N-CH-P), 4.16 m (O-$CH_2$-), 5.105 d + 5.14 d (*J* = 12.5 Hz, O-$CH_2$-), 5.18 br d (*J* = 10 Hz, -NH-), 5.41 br d (*J* = 10 Hz, >NH), 7.30 – 7.36 m (5H, $C_6H_5$). $^{13}$C NMR (150.9 MHz, $CDCl_3$; T = 25 °C): 13.85 ($-CH_3$), 14.08 ($-CH_3$), 22.26 ($-CH_2-$), 28.00 (d, *J* = 11.6 Hz, $-CH_2-$), 28.56 (d, *J* = 3.2 Hz, $-CH_2-$), 31.19 (3x >CH-), 35.60 (3x $-CH_2-$), 37.16 (d, *J* = 75.9 Hz, P-$CH_2$-), 44.03 (d, *J* = 3.2 Hz, 3x $-CH_2-$), 51.08 (d, *J* = 114.0 Hz, N-CH-P), 61.56 (O-$CH_2$-), 66.93 (O-$CH_2$-), 84.55 (d, *J* = 10.3 Hz, >CH-O), 128.49 (10x Ar –CH=),

136.46 (Ar >C=), 156.22 (d, $J$ = 7.1 Hz, O-C<), 166.93 (d, $J$ = 4.3 Hz, -O-CO-). $^{31}$P NMR (202.4 MHz, $CDCl_3$; T = 25 °C): 39.98. IR $\nu_{max}$/$cm^{-1}$ ($CHCl_3$): 3424 m (NH); 2958 m, 2873 w, 1369 w ($CH_3$); 2942 m, 2920 s, 2858 m ($CH_2$); 1725 vvs+vbr (C=O) ester, carbamate; 1508 s (amide II); 1242 s (P=O); 999 s (P-O-C); 3091 w, 3068 w, 3030, 1456 m, 1028 m, 697 m (ring). HRMS (ESI): $[M+H]^+$ for $C_{27}H_{41}NPO_6$ calc: 506.26660; found: 506.26668.

[((R,S)-1-{[(Benzyloxy)carbonyl]amino}pentyl)(adamantan-1-yloxy)phosphoryl]acetic acid **49**
To a solution of ester **48** (3.5 g, 6.9 mmol) in ethanol (30 mL) was added a solution of NaOH (0.66 g, 16.6 mmol) in water (10 mL). The reaction mixture was stirred at room temperature overnight. After completion of the reaction, solid citric acid was added to the clear solution until pH 7–8 was reached, as checked by pH indicator paper. Ethanol was then removed under reduced pressure, and after cooling in an ice bath, the pH of the mixture was further adjusted to 1–2 by careful addition of 0.01 M HCl. Water (50 mL) was added, and the mixture was extracted with ethyl acetate (4 × 30 mL). The combined organic layers were washed with water (50 mL), then with brine (2 × 50 mL), and dried over anhydrous sodium sulfate. After filtration, the solvent was removed under reduced pressure. The crude product was purified by column chromatography using a linear gradient of a chloroform/methanol/acetic acid mixture (6:2:0.5) in chloroform. The resulting clear oil was lyophilized twice from dioxane. Yield 2.9 g (88%). Colorless lyophilizate. $R_f$ = 0,70 (ethyl acetate/acetone/methanol/water 6 : 1 : 1 : 0.5). A mixture of two stereoisomers was obtained in an approximately 58:42 ratio. NMR data are reported only for the major isomer.
$^1$H NMR (600 MHz, $CDCl_3$; T = 25 °C): 0.88 t (3H, $J$ = 7 Hz, $CH_3$), 1.28 m + 1.35 m, 1.62 m + 1.85 m, 1.45 m (3x -$CH_2$-), 1.58 m + 1.62 m (3x -$CH_2$-, Ada), 2.10 m (3x -$CH_2$-, Ada), 2.17 m (3x >CH-, Ada), 2.98 d (P-$CH_2$-, $J$ = 16.7 Hz), 4.05 m (N-CH-P), 5.16 d + 5.09 d ($J$ = 12.4 Hz, O-$CH_2$-), 5.33 d ($J$ = 10.0 Hz, >NH), 7.28 - 7.36 m (10 H, 2x -$C_6H_5$). $^{13}$C NMR (150.9 MHz, $CDCl_3$; T = 25 °C): 13.83 (-$CH_3$), 22.15, 27.16, 27.86 (3x -$CH_2$-), 31.28 (3x >CH-, Ada), 35.55 (3x -$CH_2$-, Ada), 37.52 d ($J$ = 74.1 Hz, P-$CH_2$-), 44.06 d ($J$ = 3.4 Hz, 3x -$CH_2$-, Ada), 50.91 d ($J$ = 117.2 Hz, P-$CH_2$-), 67.34 (O-$CH_2$-), 85.18 d ($J$ = 10.7 Hz, O-$CH_2$-), 128.49 – 127.80 (5x Ar-CH=), 136.29 (Ar-C=), 167.16 (-COOH). $^{31}$P NMR (202.4 MHz, $CDCl_3$; T = 25 °C): 42.06. IR $\nu_{max}$/$cm^{-1}$ (KBr): 3302 m (NH); 2914 vs, 2856 m ($CH_2$); 1725 vs (C=O) carbamate; 1701 vs (C=O) acid; 1529 s (amide II); 1274 s - 1216 s (P=O); 1050 s - 985 s (P-O-C); 3091 w, 3063 w, 3034 w, 1608w, 1587 w, 1499 m, 1456 m, 737 m, 697 m (benzene ring). HRMS (ESI): $[M-H]^-$ for $C_{25}H_{35}NPO_6$ calc: 476.22075; found: 476.22064.

*N*-(Benzyloxycarbonylmethyl)-*N*-[(4-(*tert*-butoxycarbonylamino)butyl)]-[((*R*,*S*)-1-{[(benzyloxy)carbonyl]amino}pentyl)(adamantan-1-yloxy)phosphoryl]acetamide **50**

To a solution of acid **49** (2.9 g, 6.07 mmol) and amine **46** (2.04 g, 6.07 mmol) in DCM (50 mL) was added DIC (1.1 mL, 6.68 mmol) dropwise over 10 minutes under stirring and cooling in an ice bath. The reaction mixture was stirred at 0 °C for 1 hour and then allowed to warm to room temperature and stirred for an additional 12 hours. Upon completion, the solvent was removed under reduced pressure, and the resulting yellow oily residue was dissolved in ethyl acetate (25 mL) and stored at –20 °C overnight. The precipitated DIU was filtered off, and the filtrate was concentrated under reduced pressure. The crude product was purified by column chromatography using a linear gradient of ethyl acetate in cyclohexane. Yield 4.3 g (89%). Slightly yellow oil. $R_f$ = 0.76 (ethyl acetate). A mixture of stereoisomers was obtained. Unambiguous assignment of NMR signals to the individual stereoisomers was not possible. Therefore, only the chemical shift ranges corresponding to the respective types of protons and carbon atoms are reported.

$^1$H NMR (600 MHz, $CDCl_3$; T = 25 °C): 1.25 t (-$CH_3$), 1.30 – 1.60 m (6x -$CH_2$), 1.42 s (3x -$CH_3$, *t*-Bu), 1.58 m (3x -$CH_2$-, Ada), 1.95 - 2.11 m (3x -$CH_2$-, Ada), 2.10 m (3x >CH-, Ada), 3.00 - 3.10 m (2x N-$CH_2$-), 3.26 – 3.47 m (P- $CH_2$-), 4.04 – 4.14 m (P-CH-N), 5.06 – 5.20 m (2x O-$CH_2$-), 5.06 – 5.20 m (3x O-$CH_2$- + N-$CH_2$-CO), 5.52 d – 6.25 d (2x >NH), 7.26 – 7.36 m (10x Ar-CH, 2x $C_6H_5$). $^{13}$C NMR (150.9 MHz, $CDCl_3$; T = 25 °C): 13.87 ($CH_3$), 22.20 – 27.11 (5x -$CH_2$-), 28.36 (3x -$CH_3$, tBu), 31.16 – 31.20 (3x >CH-, Ada), 34.80 – 36.44 (3x -$CH_2$, Ada), 39.72 - 39.93 (2x N-$CH_2$-), 44.01 - 44.34 (3x $CH_2$-, Ada), 51.88 - 53.56 (P-CH-N), 66.62 – 67.54 (3x O-$CH_2$-), 78.92 – 79.22 (>C<, *t*-Bu), 84.16 – 84.58 (>C<, Ada), 127.86 – 128.69 (10 x Ar-CH=), 134.82 – 136.86 (2x Ar >C=), 155.96 – 156.41 (N-CO-O), 166.40 – 167.36 (N-CO-), 168.50 - 169.39 (O-CO-). $^{31}$P NMR (202.4 MHz, $CDCl_3$; T = 25 °C): 42.67, 42.61, 42.40, 42.24. IR $v_{max}$/$cm^{-1}$ ($CHCl_3$): 3454 w, 3432 w (NH); 2921 s, 2859 m ($CH_2$); 2982 m, 2874 w, 1393 w, 1368 m ($CH_3$); 1745 s (C=O) ester; 1712 vs (C=O) carbamate; 1638 s (C=O) amide; 1509 s (amide II); 1242 s (P=O); 1048 s, 995 s (P-O-C); 3093 w, 3070 w, 1587 w, 1028 w, 697 m (benzene rings); 1174 s ($(CH_3)_3$). HRMS (ESI): $[M+H]^+$ for $C_{43}H_{63}N_3PO_9$ calc: 796.42964; found: 796.42953.

*N*-[(4-*tert*-Butoxycarbonylamino)butyl]-[((*R*,*S*)-1-{[(9H-fluoren-9-ylmethoxycarbonyl)amino]pentyl}(adamantan-1-yloxy)phosphoryl)acetylamino]acetic acid **III**

Phosphinate **50** (3.9 g, 4.9 mmol) was dissolved in methanol (100 mL). To this solution, 10% Pd/C (300 mg) was added, and the mixture was subjected to hydrogenolysis in an autoclave under a hydrogen pressure of 15 psi with vigorous stirring. After 8 hours, the catalyst was filtered off through Celite, and the pad was washed with methanol (150 mL). Removal of the solvent under reduced pressure afforded a pale-yellow foam (2.6 g), which was used without further purification in the subsequent synthetic step. The resulting *N*-[(4-*tert*-butoxycarbonylamino)butyl]-[((*R*,*S*)-1-(aminopentyl)(adamantan-1-yloxy)phosphoryl)acetylamino]acetic acid was suspended with $NaHCO_3$ (1.5 g, 18.2 mmol) in water (50 mL). The reaction mixture was cooled in an ice bath, and a solution of Fmoc-OSu (1.6 g, 4.6 mmol) in dioxane (50 mL) was added dropwise over 30 minutes under stirring. The mixture was maintained at 0 °C for 1 hour and then allowed to react at room temperature overnight. The mixture was cooled again in an ice bath and acidified with 0.1 M HCl under vigorous stirring until pH 2 was reached (as monitored with pH indicator paper). Water (50 mL) was added, and the mixture was extracted with ethyl acetate (4 × 30 mL). The combined organic layers were washed with brine (2 × 50 mL) and dried over anhydrous sodium sulfate. After filtration, the solvent was removed under reduced pressure, and the residue was purified by column chromatography using a linear gradient of ethyl acetate in cyclohexane. The resulting foam was lyophilized twice from dioxane. Yield 2.77 g (71%). Colorless lyophilizate. $R_f$ = 0,58 (ethyl acetate/acetone/methanol/water 6 : 1 : 1 : 0.5). Mixture of stereoisomers was obtained. Only signals of the major isomer are given below.

$^{1}$H NMR (600 MHz, DMSO; T = 45 °C): 0.86 m (-$CH_3$), 1.29 m (>$CH_2$), 1.37 s (3x -$CH_3$, *t*-Bu),1.37 m (>$CH_2$), 1.42 m (>$CH_2$), 1.54 m (>$CH_2$), 1.54 m (3x -$CH_2$, Ada), 2.00 (3x >$CH_2$, Ada), 2.00 (N-$CH_2$-), 2.07 (3x >CH-, Ada), 2.91 m (N-$CH_2$-), 2.93 m (N-$CH_2$-), 3.86 br (>NH), 4.01 m (N-CH-P), 4.24 m (>CH-), 4.34 m (-$CH_2$-), 6.61 br (>NH), 7.32 m (2x Ar-CH=), 7.41 m (2x Ar-CH=), 7.71 (2x Ar-CH=), 7.88 (2x Ar-CH=).$^{13}$C NMR (150.9 MHz, DMSO; T = 45 °C): 13.76 (-$CH_3$), 21.53 (>$CH_2$), 24.41 (>$CH_2$), 25.30 (>$CH_2$), 26.88 (>$CH_2$), 28.28 (3x -$CH_3$, t-Bu), 30.60 (3x >CH-, Ada), 43.51 (N-$CH_2$-), 43.78 (3x >$CH_2$, Ada), 46.82 (>CH-), 65.90 (>$CH_2$), 77.34 (>C<, *t*-Bu), 81.78 (>C<, Ada), 121.35 (2x Ar-CH=, Fmoc), 125.21 (2x Ar-CH=, Fmoc), 127.28 (2x A-CH=, Fmoc), 127.68 (2x Ar-CH=, Fmoc), 140.75 (2x Ar-C=, Fmoc), 143.75 (2x Ar-C=, Fmoc), 156.22 (-CO-N), 165.3 – 166.3 (3x -CO-N). $^{31}$P NMR (202.4 MHz, $CDCl_3$; T = 25 °C): 43.73. IR $v_{max}$/$cm^{-1}$ ($CHCl_3$): 3434 w, 3312 m (NH); 2915 s, 2857 m ($CH_2$); 2977 m, 1393 m, 1366 m ($CH_3$); 1715 vvs +vbr (C=O) acid + carbamates; 1635 s (C=O) amide; 1509 s (amide II); 1246 s (P=O); 1050 s, 998 s (P-O-C); 1173 s ($(CH_3)_3$); 3095 w, 3065 w, 3041 w, 759 m, 741 m (fluorene). HRMS (ESI): $[M+H]^+$ for $C_{43}H_{61}N_3PO_9$ calc: 794.41399; found: 794.41382.

#### 1.2.5: Synthesis of precursor IV

The synthesis of the shorter precursor **IV** followed a strategy analogous to that employed for building block **III** as depicted in **Scheme 6**. The activated P(III) phosphinite ester underwent an Arbuzov reaction with benzyl bromoacetate to afford intermediate **51**, which was subsequently protected as its acid-labile adamantyl ester. Following hydrogenolytic removal of the benzyl protecting groups, intermediate **53** was isolated from the reaction mixture as the corresponding zwitterion. Similarly, the compound exhibited excellent stability, and no spontaneous autocatalytic deprotection of the adamantyl phosphinate ester was observed under the reaction or storage conditions employed. Subsequent acylation with Fmoc-OSu gave the desired precursor **IV**.

**1.2.5.1: Synthetic protocols for compounds in Scheme 6**

Benzyl [((*R*,*S*)-1-{[(benzyloxy)carbonyl]amino}pentyl)(hydroxy)phosphoryl]acetate **51**

Compound **51** was prepared by reaction of phosphinic acid **28** (6.95 g, 24.4 mmol), HMDS (20.5 ml, 122 mmol), and benzyl 2-bromoacetate (7.3 ml, 31.72 mmol) according to the protocol described for the preparation of intermediate **47**. The crude product was purified by trituration from a mixture of ethyl acetate and petroleum ether at −20 °C. The resulting solid was decanted, and residual solvents were removed under high vacuum. Yield 8.9 g (84%). Semisolid. $R_f$ = 0.56 (ethyl acetate/acetone/methanol/water 6 : 1 : 1 : 0.5).

$^1$H NMR (600 MHz, $CDCl_3$; T = 25 °C): 0.85 t (*J*=7.1 Hz, $-CH_3$), 1.25 m + 1.30 m ($-CH_2-$), 1.29 m + 1.40 m ($-CH_2-$), 1.51 m + 1.86 m ($-CH_2-$), 3.02 m ($-CH_2$-P), 4.13 m (P-CH-N), 5.06 – 5.14 m (2x $-CH_2$-O), 7.29 - 7.34 m (10H, 2x $C_6H_5$).$^{13}$C NMR (150.9 MHz, $CDCl_3$; T = 25 °C): 13.87 ($-CH_3$), 22.14 ($-CH_2-$), 27.63 ($-CH_2-$), 27.85 d (*J* = 12.2 Hz, $-CH_2-$), 35.19 d (*J* = 82.3 Hz, N-CH-P), 67.31 ($-CH_2$-O), 67.49 ($-CH_2$-O), 128.03 (Ar-CH=), 128.22 (Ar-CH=), 128.37 (2x Ar-CH=), 128.38 (2x Ar-CH=), 128.52 (2x Ar-CH=), 128.55 (2x Ar-CH=), 135.14 (Ar-C=), 136.09 (Ar-C=), 156.36 d (*J* = 5.2 Hz, O-CO-N), 165.86 (d, *J* = 4.4 Hz, O-CO-).$^{31}$P NMR (202.4 MHz, $CDCl_3$; T = 25 °C): 47.62. IR $v_{max}$/cm$^{-1}$ (KBr): 3309 m (NH); 2959 m, 2872 m ($CH_2$); 1732 vs (C=O) ester; 1708 vs (C=O) carbamate; 1538 s (amide II); 1253 s (P=O); 985 s, 973 s (P-O-H); 3088 w, 3063 w, 3033 w, 1586 vw, 1495 m, 1455 m, 1079 m, 1026 m, 755 m, 701 m (rings). HRMS (ESI) $[M-H]^-$ for $C_{22}H_{27}NO_6P$ calc: 432.15815; found: 432.15780.

Benzyl [((*R*,*S*)-1-{[(benzyloxy)carbonyl]amino}pentyl)(adamantan-1-yloxy)phosphoryl]acetate **52**

Compound **52** was prepared by reacting phosphinic acid **51** (7.2 g, 16.6 mmol), 1-bromoadamantane (4.3 g, 19.9 mmol), and $Ag_2O$ (7.7 g, 33.2 mmol) in $CHCl_3$ according to the protocol described for the preparation of intermediate **48**. The crude product was purified by column chromatography using a linear gradient of ethyl acetate in cyclohexane. The product was isolated as a colorless oil, which solidified upon standing in a refrigerator at +5 °C. Yield 7.2 g (77%). Colorless solid, m. p. 70 - 72°C. $R_f$ = 0,.65 (ethyl acetate/toluene 50 : 50). Two isomers in the ratio about 1:1 were observed. Signals for both are given either separately or as the range of signals.

$^{1}$H NMR (600 MHz, $CDCl_3$; T = 25 °C): 0.87 t + 0.88 t (-$CH_3$), 1.27 m + 1.34 m (-$CH_2$-), 1.33 m + 1.43 m (-$CH_2$-), 1.54 m + 1.87 m (-$CH_2$-), 1.56 m (3x -$CH_2$-, Ada), 2.10 m (3x >CH-, Ada), 1.94 – 2.01 m (3x -$CH_2$-, Ada), 2.91 dd + 3.14 t and 3.02 dd + 3.06 t (P-$CH_2$-), 4.09 – 4.17 m (P-CH-N), 5.10 – 5.17 m (2x O-$CH_2$-), 5.26 d + 4.94 dd (1H, >NH), 7.29 – 7.38 m (10 Ar-CH=).$^{13}$C NMR (150.9 MHz, $CDCl_3$; T = 25 °C): 13.85 (-$CH_3$), 22.18 + 22.25 (-$CH_2$-), 27.83 d + 27.96 d (*J*(C,P) = 12.5 + 11.7 Hz, -$CH_2$-), 28.40 d + 28.46 d (*J*(C,P) = 2.8 + 4.0 Hz, -$CH_2$-), 31.15 (3x -CH<, Ada), 35.63 (3x -$CH_2$-, Ada), 37.02 d + 37.94 d (*J*(C,P) = 75.7 + 74.8 Hz, P-$CH_2$-), 43.93 d + 44.08 d (*J*(C,P) = 3.6 + 3.4 Hz, 3x -CH<, Ada), 50.86 d + 51.00 d (*J*(C,P) = 111.0 + 114.5 Hz, (P-CH-N), 66.97 + 67.07 (O-$CH_2$-), 67.27 + 67.34 (O-$CH_2$-), 84.31 d + 84.63 d (*J*(C,P) = 10.4 + 10.3 Hz, >C<, Ada), 127.95 – 128.56 (10x Ar-CH=), 135.19 – 136.42 (2x Ar-C=), 156.06 d + 156.24 d (*J*(C,P) = 5.8 + 7.2 Hz, O-CO-N), 166.06 d + 166.71 d (*J*(C,P) = 4.2 + 4.4 Hz, O-CO-).$^{31}$P NMR (202.4 MHz, $CDCl_3$; T = 25 °C): 40.19, 39.87. IR $v_{max}$/$cm^{-1}$ (KBr): 3211 s (NH); 2912 vs, 2856 s ($CH_2$); 1734 vvs (C=O) ester; 1710 vs, 1700 vs (C=O) carbamate; 1546 s (amide II); 1238 s (P=O); 998 s, 979 s (P-O-C); 3082 w, 3062 w, 3035 w, 1586 w, 1499 m, 1454 s, 1028 m, 739 s, 696 s (rings). HRMS (ESI) [$M+Na^+$] for $C_{32}H_{42}NO_6NaP$ calc: 590.26420; found: 590.26447.

[((*R*,*S*)-1-Aminopentyl)(adamantan-1-yloxy)phosphoryl]acetic acid **53**

Ester **52** (7.2 g, 12.7 mmol) was dissolved in methanol (100 mL). To this solution, 10% Pd/C (300 mg) was added, and the mixture was subjected to hydrogenolysis in an autoclave under a hydrogen pressure of 15 psi with vigorous stirring. After 12 h, the catalyst was removed by filtration through Celite, and the pad was washed with methanol (150 mL). Evaporation of the combined filtrates under reduced pressure afforded a paleyellow foam, which was sonicated for 5 min in hot acetonitrile (100 mL, 80 °C). The resulting suspension was placed in freezer overnight at −20 °C and subsequently filtered through a Büchner funnel. Yield 3.8 g (86%). Colorless solid, m. p. 154 – 156 °C. $R_f$ = 0.46 (DCM/methanol/conc. $NH_4OH$ 75 : 22 : 3). Two diastereoisomers observed in the ratio ca 2 : 1. Only signals of major isomer are given below.

$^{1}$H NMR (600 MHz, DMSO; T = 25 °C): 0.88 t (*J* = 7 Hz, -$CH_3$), 1.27 m + 1.33 m (>$CH_2$), 1.48 m (>$CH_2$), 1.83 m (>$CH_2$), 1.57 m (3x >$CH_2$, Ada), 1.98 m + 2.03 m (3x >$CH_2$, Ada), 2.12 m (3x >CH-, Ada), 2.75 dd, *J* = 18.8 and 15.4 Hz + 3.19 dd (*J* = 16.2 and 15.4 Hz, P-$CH_2$-), 3.14 m (P-CH-N). $^{13}$C NMR (150.9 MHz, DMSO; T = 25 °C): 13.94 ($CH_3$), 21.91 (>$CH_2$), 27.29 (>$CH_2$), 27.52 d (*J* = 10.5 Hz, >$CH_2$), 30.65 (3x >CH-, Ada), 35.46 (3x >$CH_2$, Ada), 43.69 d (*J* = 3.4 Hz, 3x >$CH_2$, Ada), 50.90 d (*J* = 104.8 Hz, >C<, Ada), 81.36 d (*J* = 9.6 Hz, >C<, Ada), 167.64 d (*J* =

2.1 Hz, COOH).[31]P NMR (202.4 MHz, $CDCl_3$; T = 25 °C): 39.62. IR $v_{max}/cm^{-1}$ (KBr): 2914 vs, 2854 s ($CH_2$); 1652 s, 1632 s ($NH_3^+$); 1592 s, 1420 s ($COO^-$); 1225 s (P=O); 1051 vs, 998 vs, 983 s (P-O-C). HRMS (ESI) [M+H$^+$] for $C_{17}H_{31}NO_4P$ calc: 344.19852; found 344.19871.

[((*R*,*S*)-1-{[(9H-Fluoren-9-ylmethoxycarbonyl)amino]}pentyl)(adamantan-1-yloxy)phosphoryl]acetic acid **IV**

In a 250 mL round-bottom flask, acid **53** (3.6 g, 10.5 mmol) and $Na_2CO_3$ (1.1 g, 10.5 mmol) were dissolved in water (50 mL). The resulting solution was cooled in an ice bath, and a solution of Fmoc-OSu (3.6 g, 10.5 mmol) in dioxane (50 mL) was added dropwise over 30 min with continuous stirring. The reaction mixture was maintained at 0 °C for 1 h, then allowed to warm to room temperature and stirred overnight. After completion of reaction, the mixture was then recooled in an ice bath and acidified to pH 2 with 0.1 M hydrochloric acid under vigorous stirring as monitored by pH-indicator paper. Water (50 mL) was added, and the aqueous phase was extracted with ethyl acetate (4 × 50 mL). The combined organic extracts were washed with brine (2 × 50 mL), dried over anhydrous sodium sulfate, filtered, and concentrated under reduced pressure. The crude residue was purified by column chromatography using a linear gradient of ethyl acetate in cyclohexane. The purified material was dissolved in dioxane (100 mL), frozen, and lyophilized to afford the product. Yield 3.1 g (53%). Colorless lyophilizate. $R_f$ = 0.67 ($SiO_2$; ethyl acetate/aceton/methanol/water 6 : 1 : 1 : 0.5). Two diastereoisomers in the ratio ca 2 : 1 were obtained. Only the signals for the major isomer are given below.

[1]H NMR (600 MHz, DMSO; T = 25 °C): 0.85 t (*J* = 7 Hz, $CH_3$), 1.23 m + 1.31 m (>$CH_2$), 1.23 m + 1.33 m (>$CH_2$), 1.48 – 1.56 m (3x >$CH_2$, Ada), 1.58 m + 1.73 m (>$CH_2$), 1.95 – 2.05 m (3x >$CH_2$, Ada), 2.09 m (3x >CH-, Ada), 3.79 m (N-CH-P), 4.23 bt (*J* = 7.0 Hz, >CH-), 4.31 dd + 4.37 dd (*J* = 10.6 and 7.0 Hz, O-$CH_2$-), 7.32 m + 7.41 m + 7.72 m + 7.89 m (8x Ar-CH=, Fmoc), 7.52 d (*J* = 10.8 Hz, (>NH), 12.45 br (-COOH).[13]C NMR (150.9 MHz, DMSO; T = 25 °C): 13.94 (-$CH_3$), 21.67, 27.26 and 27.85 (3x >$CH_2$), 30.68 (3x >CH-, Ada), 35.49 (3x >$CH_2$, Ada), 36.39 d (*J* = 76.0 Hz, P-$CH_2$-), 43.60 (3x >$CH_2$, Ada), 46.91 (>CH-, Fmoc), 51.63 d (*J* = 112.1 Hz, P-CH-N), 65.76 (O-$CH_2$-), 82.03 d (*J* = 9.8 Hz, >C<, Ada), 120.58 (2x Ar-CH=, Fmoc), 125.37 + 125.49 (2x Ar-CH=, Fmoc), 127.13 + 127.22 (2x Ar-CH=, Ada), 127.79 (2x Ar-CH=, Ada), 140.90 (2x Ar-C=, Ada), 143.83 + 144.06 (2x Ar-C=, Ada), 156.46 d (*J* = 5.0 Hz, O-CO-N), 167.76 d (*J* = 6.3 Hz, COOH). [31]P NMR (202.4 MHz, $CDCl_3$; T = 25 °C): 41.84. IR $v_{max}/cm^{-1}$ (KBr): 3293 m (NH); 2914 vs, 2856 s ($CH_2$); 1725 vvs (C=O) acid; 1703 vvs (C=O) carbamate; 1531 s (amide II); 1244 s (P=O); 1005 vs, 985 s (P-O-C); 3065 w, 3041 w, 1610 w, 1581 w, 1478 m, 1451 m, 759 m, 741 s (fluorene). HRMS (ESI) [M+Na$^+$] for $C_{32}H_{40}NO_6NaP$ calc: 588.24855; found: 588.24878.

#### 1.3: Synthesis of peptides and pseudopeptides

Peptides were synthetized by solid phase synthesis using the Fmoc/HBTU/DIPEA or Fmoc/DIC/HOBT strategy on a Spyder Mark IV Multiple Peptide Synthesizer (European Patent application EP17206537.7), using Fmoc-Tyr(tBu)-Wang resin. Pseudo-dipeptide precursors and modified amino acids were coupled manually. All peptide-based compounds were purified by reversed-phase high-performance liquid chromatography (RP-HPLC); their purity was higher than 95%. The identities of peptides and pseudopeptides were confirmed by high-resolution mass spectrometry (HR-MS) **(Figures S4-S6**).

##### 1.3.1: Quenched fluorescence substrates Q-BP5 and FS-6, peptide 1 and pseudopeptides 2-8

Quenched fluorescence substrates Q-BP5 and FS-6 were synthetized by standard solid phase synthesis procedure and the non-standard amino acids were coupled manually. Pseudopeptides **2**–**8** were synthesized by solid-phase peptide synthesis (SPPS) using the Fmoc/*t*Bu strategy. The synthesis was initiated by removal of the Fmoc protecting group from commercially available Fmoc-Tyr(*t*Bu)-Wang resin. Subsequently, amino acids 2–7 were automatically coupled for inhibitors **2**–**4**, whereas amino acids 2–8 were coupled for inhibitors **5**–**8**. Each coupling reaction was performed twice using two different coupling mixtures: (i) Fmoc-AA-OH (4 equiv.), DIC (4 equiv.), and HOBt (4 equiv.) in DMF; and (ii) Fmoc-AA-OH (4 equiv.), HBTU (4 equiv.), and DIPEA (7 equiv.) in DMF. Each coupling step was allowed to proceed for 1.5 h. Fmoc deprotection was carried out using 20% piperidine in DMF for 5 and 20 min.

Phosphinate building blocks **I** and **II** bearing a free P–OH group were coupled manually using a mixture of **I** or **II** (1.1 equiv.), EDC·HCl (5 equiv.), HOBt (1.1 equiv.), and DIPEA (2.1 equiv.) in DMF/DCM. The coupling reaction was performed twice, each for 1 h. Completion of the reaction was confirmed by a negative Kaiser test. The remaining amino acids 11–34 were subsequently coupled automatically for inhibitors **2** and **3** using a mixture of Fmoc-AA-OH (4 equiv.), BOP (4 equiv.), and DIPEA (7 equiv.) in DMF. Each coupling step was performed twice for 1.5 h.

Phosphinate building blocks **III** and **IV** bearing a protected P–O-Ad group were coupled manually using a mixture of **III** or **IV** (5 equiv.), HOBt (5 equiv.), and DIC (5 equiv.) in DMF. The coupling reaction was performed twice, each for 18 h. Completion of the reaction was confirmed by a negative Kaiser test. The remaining amino acids 11–34 (inhibitors **4** and **5**), 11–25 (inhibitor **6**), 11–20 (inhibitor **7**), and 11 (inhibitor **8**) were subsequently coupled automatically. Coupling reactions were carried out using either (i) Fmoc-AA-OH (4 equiv.), DIC (4 equiv.), and HOBt (4 equiv.) in DMF or (ii) Fmoc-AA-OH (4 equiv.), HBTU (4 equiv.), and DIPEA (7 equiv.) in DMF. Each coupling step was performed for 1.5 h.

After completion of peptide assembly, the resin was washed with DMF and DCM and dried overnight under high vacuum in a desiccator. Peptides were cleaved from the resin using a TFA/TIPS/$H_2O$ (95:2.5:2.5, v/v/v) cleavage cocktail for 2 h and precipitated with ice-cold diethyl ether (−80 °C). The crude peptides were collected by centrifugation and purified by preparative RP-HPLC on a Nucleosil 100-5 C8 column (250 × 10 mm, Macherey–Nagel) at a flow rate of 4 mL.min$^{-1}$ using a linear gradient from 10% B at t = 0 min to 100% B at t = 30 min on an ECOM ECS21 HPLC system (A = 0.1% TFA in water; B = 0.1% TFA in 80% $CH_3CN$). Elution was monitored at 218 and 254 nm.

The purity of the final compounds was assessed by analytical RP-HPLC using a Nucleosil 120-5 C8 column (250 × 4.6 mm, Macherey–Nagel) at a flow rate of 1 mL.min$^{-1}$ on a Watrex HPLC system equipped with a DeltaChrom Pump and LC100 UV Detector. Compound identities were confirmed by mass spectrometry.

#### 1.3.2: Pseudo-dipeptide 9

Two synthetic approaches were evaluated for the preparation of inhibitor **9** (**Scheme 7**).

The first followed the strategy previously developed for the synthesis of building block **II**. Acrylate precursor **55** was prepared from Meldrum's acid, while the Ac-Nleψ[PO(OH)H] fragment was synthesized from phosphinic acid **27** by treatment with acetic anhydride. A phospha-Michael addition between these two components afforded intermediate **56**, which, upon hydrogenolytic removal of the protecting groups, yielded the target inhibitor **9**.

Alternatively, phosphinic precursor **II** (48 mg, 78 μmol) was dissolved in 10 mL of 20% piperidine in a 1:1 mixture of DCM and DMF. After stirring for 20 min at room temperature, the reaction mixture was concentrated under reduced pressure. Acetic anhydride (1 mL) was then added to the residue, and after 10 min the reaction mixture was again evaporated under reduced pressure. TFA (5 mL) was added to the resulting residue, and after 30 min the mixture was concentrated under reduced pressure. Cold diethyl ether was added to the residue, resulting in precipitation of the product. The precipitated product was purified and purity confirmed by RP-HPLC and identity verified by HR-MS (Figure S5E). A pale yellow lyophilizate was obtained. Yield 4 mg (15%).

##### 1.3.2.1: Synthetic protocols for compounds in Scheme S7

2,2-Dimethyl-5-[4-(benzyloxycarbonylamino)butyl]-1,3-dioxane-4,6-dione **54**

Intermediate **54** was prepared by reaction of Cbz-GABA-OH (14.2 g, 60 mmol), Meldrum's acid **39** (8.6 g, 60 mmol), DMAP (11 g, 90 mmol), DIC (10.3 mL, 66 mmol) and (4.8 g ; 127.9 mmol) $NaBH_4$ by using protocol for synthesis compound **40**. Yield 12.8 g (62 % over two steps). Colorless solid, m. p. 96 -98 °C. $R_f$ = 0.62 (ethyl acetate/toluen 50 : 50). $^1$H NMR (600 MHz, $CDCl_3$; T = 25 °C): 1.49 m (2H, >$CH_2$), 1.57 m (2H, >$CH_2$), 1.75 s and 1.78 s (6H, 2x -$CH_3$), 2.10 m (2H, >$CH_2$), 3.22 br q (2H, -$CH_2$-N), 3.52 t (1H, *J* = 5.0 Hz, >CH-), 4.88 br (1H, >NH), 5.09 s (2H, O-$CH_2$-), 7.30 – 7.36 m (5H, -$C_6H_5$). $^{13}$C NMR (150.9 MHz, $CDCl_3$; T = 25 °C): 25.51 (>$CH_2$), 25.78 (>$CH_2$), 26.75 and 28.40 (2x -$CH_3$), 29.68 (>$CH_2$), 40.44 (2H, -$CH_2$-N), 46.04 (>CH-), 66.58 ( O-$CH_2$-), 104.90 (O-C-O), 128.04 (3x Ar-CH=), 128.48 (2x Ar-CH=), 136.57 (Ar-C=), 156.37 (O-CO-N), 165.37 (2x O-CO-). IR $v_{max}/cm^{-1}$ (KBr): 3345 s (NH); 2949 m, 2874 m, 1393 m ($CH_3$); 2932 m, 2853 m ($CH_2$); 1799 s, 1745vs (C=O) ester; 1689 vs (C=O) carbamate; 1541 s (amide II); 3092 w, 3063 w, 3036 w, 1026 m, 973 m, 753 m (phenyl). HRMS (ESI): $[M-H]^-$ for $C_{18}H_{22}NO_6$ calc: 348.14526; found: 348.14492.

Benzyl 6-[(benzyloxycarbonyl)amino]-2-methylenehexanoate **55**

Intermediate **55** was prepared by reaction of dioxane **54** (12.4 g, 35.5 mmol), benzyl alcohol (7.7 g, 71 mmol), paraformaldehyde (1.1 g, 35.5 mmol) and piperidine (350 μl, 3.55 mmol) by using protocol for synthesis compound **41**. Yield 9.2 g (71%). Slight yellow oil. $R_f$ = 0.68 (ethyl acetate/toluen 20 : 80). $^1$H NMR (600 MHz, $CDCl_3$; T = 25 °C): 1.50 m (2H, >$CH_2$), 1.52 m (2H, >$CH_2$), 2.34 m (2H, >$CH_2$), 3.20 br q (2H, -$CH_2$-N), 4.79 br (1H, >N-H), 5.09 s (2H, O-$CH_2$-), 5.19 s (2H, O-$CH_2$-), 5.56 br and 6.20 d (2H, =$CH_2$), 7.30 – 7.37 m (10H, 2x -$C_6H_5$).$^{13}$C NMR (150.9 MHz, $CDCl_3$; T = 25 °C): 25.50 (>$CH_2$), 29.41 (>$CH_2$), 31.42 (>$CH_2$), 40.72 (2H, -$CH_2$-N), 66.37 (O-$CH_2$-), 66.55 (O-$CH_2$-), 125.33 (>$CH_2$), 127.97 (2x Ar-CH=), 128.04 (2x Ar-CH=), 128.13 (2x Ar-CH=), 128.46 (2x Ar-CH=), 128.51 (2x Ar-CH=), 135.99 (Ar-C=), 136.57 (Ar-C=), 140.13 (>C=), 156.34 (O-CO-N), 166.89 (O-CO-). IR $v_{max}/cm^{-1}$ ($CHCl_3$): 3452 m (NH), 2943 m, 2866 m ($CH_2$); 1630 m (C=$CH_2$); 1714 vvs (C=O) ester + carbamate; 1515 s (amide II); 3092 w, 3068 w, 1609 w, 1455 m, 1087 w, 698 m (phenyl). HRMS (ESI): $[M+Na]^+$ for $C_{22}H_{25}NNaO_4$ calc: 390.16758; found: 390.16758.

Benzyl (*R*,*S*)-2-[((*R*,*S*)-1-{acetylamino}pentyl)(hydroxyphosphoryl)methyl]-6-[(benzyloxycarbonyl)amino]hexanoate **56**

Intermediate **56** was prepared by reaction of ester **55** (1.4 g, 3.84 mmol), phosphinic acid **57** (0.63 g, 3.2 mmol), and HMDS (3.35 ml, 16 mmol) by using protocol for synthesis compound **42**. Yield 1.3 g (72 %). Light yellow oil. $R_f$ = 0.53 (ethyl acetate/aceton/methanol/water 6 : 1 : 1 : 0.5).

Mixture of two stereoisomers in the ratio ca 55:45. NMR data for major isomer are given only. $^1$H NMR (600 MHz, DMSO; T = 25 °C): 0.84 t (3H, *J* = 7.0 Hz, -$CH_3$), 1.16 m (2H, >$CH_2$), 1.23 m (2H, >$CH_2$), 1.33 m (2H, >$CH_2$), 1.71 m (2H, >$CH_2$), 1.84 s (3H, -$CH_3$), 1.91 m (2H, P-$CH_2$-), 2.70 m (1H, >CH-), 2.92 m (2H, N-$CH_2$-), 3.92 m (N-CH-P), 4.99 s (2H, O-$CH_2$-), 5.05 d and 5.09 d (2H, *J* = 12.5 Hz, O-$CH_2$-), 7.29 – 7.37 m (10H, 2x -$C_6H_5$), 8.02 d (1H, *J* = 9.2 Hz, >N-H).$^{13}$C NMR (150.9 MHz, DMSO; T = 25 °C): 14.00 (-$CH_3$), 21.86 (>$CH_2$), 22.58 (-$CH_3$), 26.72 >$CH_2$), 27.96 d (*J* (C,P) = 10.8 Hz, >$CH_2$), 28.87 d (*J* (C,P) = 88.3 Hz, P-$CH_2$-), 29.41 (>$CH_2$), 33.04 d (*J* (C,P) = 8.2 Hz, >$CH_2$), 38.88 (>CH-), 48.21 d (*J* (C,P) = 105.5 Hz, P-$CH_2$-), 65.28 (O-$CH_2$-), 65.77 (O-$CH_2$-), 127.89 - 137.46 (10 x Ar-CH= and 2x Ar-C=), 156.25 (O-CO-N), 169.50 (-CO-N), 174.52 (O-CO-). $^{31}$P NMR (202.4 MHz, DMSO; T = 25 °C): 54.64. IR $v_{max}$/$cm^{-1}$ (film): 3303 m (NH); 2954 s ($CH_3$); 2934 m, 2862 m ($CH_2$); 1729 vs (C=O) ester, 1704 vs (C=O) carbamate; 1661 s (C=O) acetyl; 1537 s (amide II); 1172 s (P=O); 969 m (P-O-H); 3094 w, 3064 w, 3034 w, 1586 m, 1499 s, 1455 s, 1083 m, 1027 m, 751 m, 738 m, 698 s, 698 m (rings). HRMS (ESI) $[M+Na]^+$ for $C_{29}H_{41}N_2NaO_7P$ calc: 583.25436; found: 583.25384.

(*R,S*)-2-[((*R,S*)-1-{acetylamino}pentyl)(hydroxyphosphoryl)methyl]-6-amino hexanoic acid **9**
Phosphinic acid **56** (0.6 g, 1.1 mmol) was dissolved in methanol (40 mL). A 10% Pd/C (100 mg) was added, and the reaction mixture was subjected to hydrogenolysis in an autoclave under a hydrogen atmosphere (15 psi) with vigorous stirring. After 12 h, the catalyst was removed by filtration through a Celite pad, which was subsequently washed with methanol (40 mL). The combined filtrates were concentrated under reduced pressure to afford a residue, which was purified by using preparative RP-HPLC and lyophilizied from water. Yield 304 mg (90 %). Colorless lyophilizate. Mixture of two stereoisomers in the ratio ca 60 : 40. NMR data for major isomer are given only. $^1$H NMR (600 MHz, DMSO; T = 25 °C): 0.84 t (3H, *J* = 7.0 Hz, -$CH_3$), 1.17 m and 1.33 m (2H, >$CH_2$), 1.20 m and 1.29 m (2H, >$CH_2$), 1.28 m (2H, >$CH_2$), 1.44 and 1.72 m (2H, >$CH_2$), 1.52 m (2H, >$CH_2$), 1.55 m and 1.63 m (2H, >$CH_2$), 1.62 m and 1.88 m (2H, >$CH_2$), 1.85 s (3H, -$CH_3$), 2.59 m (2H, >$CH_2$), 2.77 m (2H, N-$CH_2$-), 3.93 m (1H, N-CH-P), 7.74 br (2H, -$NH_2$), 7.98 d (1H, *J* = 9.2 Hz, >N-H). $^{13}$C NMR (150.9 MHz, DMSO; T = 25 °C): 14.03 (-$CH_3$), 21.87 (-$CH_2$-), 22.63 (-$CH_3$), 23.31 (-$CH_2$-), 26.78 (-$CH_2$-), 26.98 (-$CH_2$-), 28.00 (-$CH_2$-), 28.45 d (*J* (C,P) = 87.5 Hz, P-$CH_2$-), 32.05 d (*J* (C,P) = 6.6 Hz, >$CH_2$), 38.57 (>CH-), 38.74 (N-$CH_2$-), 48.19 d (*J* (C,P) = 105.4 Hz, N-CH-P), 169.50 d, (*J* (C,P) = 4.0 Hz, -CO-N), 176.08 d (*J* (C,P) = 10.0 Hz, -COOH). $^{31}$P NMR (202.4 MHz, DMSO; T = 25 °C): 45.88. IR $v_{max}$/$cm^{-1}$ (KBr): 3272 w; 2960 s, 2874 m, 1376 m ($CH_3$); 2940 s, 2862 m ($CH_2$); 1697 s (C=O) acid; 1656 s (C=O) acetyl; 1545 m

(amide II); 1203 vs, 1182 vs (P=O); 957 m (P-O-H). HRMS (ESI) $[M-H]^-$ for $C_{14}H_{28}N_2O_5P$ calc: 335.17413; found: 335.17408.

((*R*,*S*) -1-[Acetylamino]pentyl)phosphinic acid **57**

Phosphinic acid **27** (1.00 g, 6.61 mmol) was dissolved in an aqueous solution of $NaHCO_3$ (1.11 g, 13.2 mmol) in water (10 mL). The resulting clear solution was cooled to 0 °C, and acetic anhydride (687 µL, 7.2 mmol) in dioxane (2 mL) was added dropwise under continuous stirring. The reaction mixture was stirred at 0 °C for 1 h and then allowed to warm to room temperature. After 18 h, trifluoroacetic acid was added until the pH value reached approximately 1. The mixture was frozen at −78 °C and lyophilized. Purification of the crude product by RP-HPLC and lyofilization from water afforded the desired product. Yield (0.65 g, 51%). Colourless lyophilizate. $^1H$ NMR (600 MHz, DMSO; T = 25 °C): 0.86 t (3H, $J$ = 7.0 Hz, $-CH_3$), 1.23 m and 1.29 m (2H, $>CH_2$), 1.23 m and 1.35 m (2H, $>CH_2$), 1.46 m and 1.66 m (2H, $>CH_2$), 1.86 s (3H, $-CO-CH_3$), 3.86 m (1H, >CH-N), 6.73 d (1H, $J$ = 531.8 Hz, -O-P(=O)-H), 8.08 d (1H, $J$ = 8.9 Hz, >N-H). $^{13}C$ NMR (150.9 MHz, $CDCl_3$; T = 25 °C): 13.97 ($-CH_3$), 21.89 ($>CH_2$), 22.53 ($-CH_3$), 25.85 d ($J$ (C,P) = 2.3 Hz, $>CH_2$), 27.85 d ($J$(C,P) = 11.6 Hz, $>CH_2$), 48.38 d ($J$(C,P) = 104.8 Hz, N-CH-P), 169.62 d ($J$(C,P) = 3.6 Hz, N-CO-). IR $v_{max}/cm^{-1}$ (KBr): 3203 m (NH); 2974 m, 2955 m, 2874 m, 1375 m ($CH_3$); 2936 m, 2861 m, 1456 s ($CH_2$); 1652 m (C=O); 1539 s (amide II); 1180 vs (P=O); 966 vs (P-O-H). HRMS (ESI) $[M+Na]^+$ for $C_7H_{16}NNaO_3P$ calc: 216.07600; found: 216.07626.

### 2. Cell culture

Lung cancer cell line A549 (ATCC; CCL-185) and human embryonic kidney HEK293T cell line (ATCC; CRL-3216) were grown in DMEM- high glucose 5 mM (Biosera) supplemented with 10 % fetal bovine serum (FBS), 2 mM of L-glutamine and 100 units/ml penicillin/streptomycin in humidified air with 5 % $CO_2$ at 37 °C.

Cell lines used for cytotoxicity assay were: CCRF-CEM (ATCC; CCL-119), HepG2 (ATCC: HB-8065), Hela S3 (ATCC: CCL-2.2), HL-60 (ATCC: CCL-240) and non-tumor human dermal fibroblasts NHDF (ATCC: PCS-201-012). The cells were maintained in RPMI-1640 (CCRF-CEM, HL60 and Hela), MEM (HepG2) or DMEM (NHDF) culture medium containing 10% FBS and 1% GlutaMax without antibiotics and were grown in humidified air with 5 % $CO_2$ at 37 °C.

### 3. Recombinant PAPP-A preparation

Recombinant PAPP-A was expressed in HEK293T cells. After 24 h of seeding, the cells ($3 \times 10^6$ cells per 150 $cm^2$ cultivation flask) were transiently transfected with 2 µg/mL DNA of WT PAPP-

A in pcDNA3.1-C-(k)DYK vector (Genscript (cloneID: OHu20096) using linear polyethylenimine (PEI). DNA was preincubated with PEI for 20 min in a ratio of 1:6.5 in 1 mL of Opti-MEM™ medium and added to the cells., Valproic acid 3 mM was added 24 h post-transfection. The cells were grown for 5 days in total.

PAPP-A was isolated from conditioned media of transiently transfected HEK293T cells. The conditioned medium was centrifuged for 5 min at 240 x g and the supernatant was centrifuged for another 5 min at 2701 x g. The supernatant was filtered and concentrated using 100 kDa MWCO concentrator columns (Millipore UFC9100) to 20 ml. PAPP-A was purified by FLAG affinity chromatography (Genscript Anti-DYKDDDDK G1 Affinity resin) and eluted with 0.2 mg/ml 3×-DYKDDDDK peptide (Genscript) in TBS according to manufacturer's standard procedure. The protein was aliquoted and stored in -80 °C for further use. Concentration of the PAPP-A dimer was estimated from absorbance at 280 nm with $\varepsilon = 468170\ cm^{-1} \times M^{-1}$ based on amino acid sequence. The effectiveness of the transfection was checked by Western Blotting, using monoclonal anti-FLAG antibody conjugated with HRP (Sigma-Aldrich, A8592).

**4. Quenched fluorescence assay**

The inhibitory potency of the compounds was measured by quenched fluorescence assay. We designed and synthesized the substrate Q-BP5 to measure the activity of recombinant PAPP-A. The measurements with PAPP-A were performed in reaction buffer 50 mM Tris-HCl pH 8.0 with 1 mM $CaCl_2$ and 0.01% Triton X-100. We also tested commercial PAPP-A2 (Sino Biological, cat.no. 10528-H08H).

We synthesized an established substrate, FS-6 to measure the activity of commercial MMP-1 (Sino Biological, cat.no. 10532-H08H) and MMP-7 (a kind gift from Prof. Sagi, WIS, Israel). MMP-1 and MMP-7 were activated by 1 mM *p*-Aminophenylmercuric Acetate, APMA (Sigma Aldrich, cat.no. 164610) prior to testing. The measurements with MMP-1 and MMP-7 were performed in reaction buffer 100 mM Tris-HCl pH 7.5, 100 mM NaCl, 10 mM $CaCl_2$ with 0.01% Triton and 1% DMSO.

The substrates were serially diluted to yield final concentrations of 0.78 – 100 µM to assess $K_m$ values of the enzymes. Concentrations of enzymes were 4 nM of PAPP-A dimer, 10 nM MMP-1 and 2 nM MMP-7. Optimal concentration of the enzymes was checked in preliminary experiments using dilutions of the enzyme with substrate. For the $IC_{50}$ measurement, the substrate Q-BP5 concentration was set at 10 µM and the inhibitors were serially diluted in the reaction buffer using BRAVO pipetting station (Agilent, USA) (6 orders of magnitude from the concentration range $10^{-11}$ to $10^{-3}$ M were included depending on inhibitor's potency). In the case of MMP-1 and MMP-7, the FS-6 substrate concentration was 5 µM and the inhibitors were tested at a single concentration of 100 µM. Batimastat (Calbiochem, cat.no. 196440) was tested in the concentration range $10^{-11}$ - $10^{-4}$ M with MMPs and at 100 µM with PAPP-A. STC-1 (Sino Biological, 15411-H08H) was tested only with PAPP-A at 100 nM.

Optimization reactions and $K_m$ measurements were pipetted manually. For the PAPP-A inhibition assessment, the inhibitors were pre-spotted onto black 384-well low-volume U-bottom plates (Corning, USA) using an ECHO 650 liquid handler (Beckman Coulter, USA) to obtain desired final concentrations. The substrate Q-BP5 and PAPP-A were added equally using a CERTUS FLEX liquid dispenser (Fritz Gryger AG, Switzerland) to a final volume of 5 µl. The plate was temporarily sealed with Duct tape (Hampton Research, USA) and shaken for 15 s at 800 rpm. The seal was removed, and the fluorescence was measured using Tecan Spark plate reader (Tecan, Switzerland) with excitation at 310 nm and emission at 420 nm for 1 h at room temperature.) All samples were run as triplicates, experiments were repeated at least twice.

The initial slope data (from 1-11 min) were analyzed using GraphPad Prism 8 (GraphPad, USA), applying the Michaelis-Menten non-linear regression model for $K_m$ measurement. $IC_{50}$ values were assessed by non-linear fitting of log(inhibitor) *vs*. response variable slope. To estimate the inhibition constant (*K*i), we recalculated the $IC_{50}$ data using Cheng-Prussoff equation ($K_i = IC_{50}/(1+[S]/K_m)$).

**5. Assessment of cleavage of IGFBP-4 and IGFBP-5 by PAPP-A using Western Blot**

The IGFBP-4 (Sino Biological, 10967-H08H) and IGFBP-5 (Sino Biological, 10206-H08B) cleavage by PAPP-A was followed using Western blot. The cleavage reactions were performed in total reaction volume 15 µL in reaction buffer (50 mM Tris-HCl pH 8,0; 1 mM $CaCl_2$). IGFBP-4 at final concentration 333 nM was pre-incubated with IGF-1 (Tercica, Inc.) at final concentration 1 µM (ratio 1:3) at 37 °C for 45 min. IGFBP-5 was used at the same concentration as IGFBP-4 but without IGF-1 addition. IGFBPs were incubated with PAPP-A (about 2 nM of dimer) for 1 h at 37 °C in absence or presence of serially diluted inhibitors (0.04 µM to 10 µM). The reaction was stopped by addition of 1 × Laemmli sample buffer (Bio-Rad) with 10 % 2-mercaptoethanol.

Time profile (1 h - overnight) of the reaction was performed in the same conditions with inhibitors **3**, **5** and peptide **1** at final concentration 3.3 µM.

The IGFBP-4 and IGFBP-5 cleavage was visualized using SDS-PAGE with 12 % acrylamide gel and subsequent Western blot analysis with an anti-IGFBP-4 primary antibody (Abcam, cat. no. AB205581; dilution 1:1000) and anti-6 × His tag primary antibody (Invitrogen, cat. no. MA1-135; dilution 1:1000) for IGFBP-5. The experiment was repeated 3 times.

**6. RT-qPCR for mRNA expression along STC-PAPP-A-IGFBP-IGF axis**

The *PAPP-A*, *PAPP-A2*, *STC1/2*, *IGFBP-2/4/5* and *IGF-1/2* mRNA expressions in the cell line A549 were assessed using standard RT-qPCR procedures. The expressions under normal growth conditions, in starved cells and after stimulation with 10nM IGF-1 for 2 or 4 hours were determined. Used primers are listed in Table S3.

Briefly, for RNA isolation, cells were lysed in TRI Reagent (Sigma Aldrich). Samples were incubated for 5 min at room temperature (RT), centrifuged (12,000 x g, 10 min, 4 °C), and the supernatant was transferred to a new tube. Chloroform in ratio 1:5 was added, and the samples were vortexed for 15 s, incubated for 5 min at RT, and centrifuged (12,000 x g, 15 min, 4 °C). The aqueous phase was transferred to new tubes, 100% ethanol in ratio 1:1 was added, and subsequently, the Direct-zol RNA Miniprep Kit (Zymo Research) was used according to the manufacturer's instructions. For subsequent cDNA synthesis, LunaScript RT SuperMix Kit (New England Biolabs) was used according to the manufacturer's instructions. The qPCR was performed using LightCycler 480 SYBR Green I Master (Roche) according to the manufacturer's instructions on LightCycler 480 II (Roche). qPCR analyses were performed on samples from six independent biological replicates across two experimental batches. First batch (n = 3) included control, starvation, and 4 h treatment conditions and expression of *PAPP-A*, *PAPP-A2*, *STC1/2, IGFBP-2/4/5* was analyzed. The second batch (n = 3) included an additional 2 h treatment condition and additional *IGF-1/2* genes, while maintaining the same experimental setup. Data were combined after normalization.

Samples were analyzed in technical triplicate. Replicates with Ct values differing by more than 0.5 cycle were excluded from the analysis. Relative mRNA expression levels were normalized to *GAPDH* and calculated using the $2^{-\Delta\Delta Ct}$ method relative to the control.

Statistical significance was calculated using ΔCt values and one-way ANOVA with Tukey post-hoc test, *$p < 0.05$, **$p < 0.01$, ***$p < 0.001$, ****$p < 0.0001$.

**7. Inhibition of IGF-1 signaling in a cell-based assay**

Lung cancer cell line A549 was shown to secrete PAPP-A[7]. We showed PAPP-A expression in the A549 cell lysates on protein level using western blot. Cell lysates of control cells and after overnight starving were compared. The cells were lysed in RIPA buffer (Sigma Aldrich) with protease inhibitor cocktail (Sigma Aldrich). Protein concentration was measured by Bradford Assay. Cell lysates (10-15 μg) were separated by SDS-PAGE (8% acrylamide gel) and visualized on Western blot using anti-PAPP-A antibody (Abcam, cat. no.: EPR12897, dilution 1:1000) and anti-GAPDH antibody (Cell Signaling, cat. no.: 97166S, dilution 1:1000).

Next, we used A549 cell line to assess the ability of inhibitors to attenuate IGF-1 signaling through inhibition of its liberation from IGFBPs. Confluent cells in 24 well plates were placed to the growth media without serum and incubated overnight to allow for PAPP-A accumulation. IGF-1 was preincubated with IGFBP-4 for 30 min in a ratio 1:2, final concentration was 10 nM IGF-1 and 20 nM IGFBP-4 and added to the cells for two hours. The cells were pre-treated with the inhibitors (1 – 1000 nM) for 1 h. The potency of synthetic inhibitors was compared with 0.1 - 10 nM STC1 (Sino Biological, 15411-H08H). The cells were lysed in electrophoresis SDS sample buffer (100 μL/well of 62.5 mM Tris/HCl (pH 6.8), 2% SDS (w/v), 10% glycerol (v/v), 0.01% Bromophenol Blue (w/v), 0.1M DTT (w/v), 50 mM NaF, 1 mM $Na_3VO_4$ and 0.5% protease inhibitor cocktail (Sigma-Aldrich)). The samples were analyzed using standard Western blotting. Autophosphorylation of the receptor was detected using phospho-IGF-1Rβ (Tyr1135/1136)/IRβ (Tyr1150/1151) (19H7), Cell Signaling, and phosphorylation of Akt using phospho-Akt (Thr308) (C31E5E), Cell Signaling. Anti-actin (20-33), Sigma-Aldrich, was used as a loading control.

#### 8. Inhibition of IGF signaling in zebrafish *in vivo*

Two zebrafish models, Tg(igfbp5a:GFP) and *stc1a*(+17)$^{+/-}$;Tg(igfbp5a:GFP), were used to determine whether the pseudopeptides inhibit Papp-aa activity *in vivo*. Zebrafish were raised, maintained, crossed, and staged according to standard zebrafish husbandry guidelines[8]. Embryos and larvae were maintained at approximately 28°C in standard E3 embryo medium.

The compositions of the control embryo medium, containing 0.2 mM $Ca^{2+}$, and the induction embryo medium, containing 0.001 mM $Ca^{2+}$, have been reported previously[9]. To quantify changes in the number of $Na^+$/$K^+$-ATPase-rich ionocytes (NaR cells), zebrafish larvae were anesthetized in embryo media containing 0.168 mg/mL tricaine, mounted in 0.3% low-melting-point agarose, and immersed in 1 mL of medium. Images were acquired using a Zeiss LSM 880 confocal microscope (Carl Zeiss Microscopy GmbH, Jena, Germany) and analyzed using Fiji.

Because *stc1a*$^{-/-}$ fish do not survive beyond 2 weeks, the line was maintained as *stc1a*$^{+/-}$;Tg(igfbp5a:GFP). In each experiment, adult *stc1a*$^{+/-}$;Tg(igfbp5a:GFP) fish were intercrossed. The resulting embryos, consisting of a mixture of *stc1a*$^{+/+}$ wild-type, *stc1a*$^{+/-}$ heterozygous, and *stc1a*$^{-/-}$ homozygous mutant genotypes, were raised and used.

### 9. Cytotoxicity assay

Pseudopeptides **3**, **5**, **6**, **7**, **8** and **9** were tested for cytotoxicity by standard procedure employing CellTiter-Glo® 2.0 assay (Promega, Madison, USA). Four cancer cell lines (CCRF-CEM, HepG2, Hela S3, HL-60) and non-tumor human dermal fibroblasts (NHDF) were used. The A549 cell line was tested with pseudopeptides **3**, **5** and **8**. Compounds were spotted to 384-well white plates (Thermo Fisher Scientific, Waltham, USA) by ECHO550 Acoustic Liquid Handler (Beckman). The cells were incubated with the compounds 0.125-100 μM at 37°C, 5% $CO_2$ for 72 hours. Luminescence was measured by a multimode plate reader. The signal of the compound-treated cells was related to the value of untreated control which was arbitrarily set to 100% viability. $IC_{50}$ values were calculated from dose-response curves using non-linear regression method using GraphPad Prism software.

### 10. Stability in plasma

Pseudopeptide **5** (20 μM, a variant without C-terminal Y, Mw 3854.5) was incubated in 25 μL of human plasma (Biowest) or mouse plasma (Innovative Research) for 0, 15, 60 and 120 min at 37°C. The proteins were precipitated with 25 μL of 90% acetonitrile, 0.1% TFA for 1 hour and separated by centrifugation (5 min at 14,000 x g). Sediment was discarded and compounds contained in supernatant were after lyophilization resuspended in 20 μL of deionized water and analyzed using MALDI-TOF. The MALDI-TOF spectra were measured on rapifleX MALDI-TOF/TOF mass spectrometer (Bruker Daltonics, Germany) with Smartbeam 3D™ laser with up to 10 kHz repetitions. The measurements were done in positive reflectron mode technique, with the mass range 0.8-6 kDa. The accelerating voltage was set at 20 kV. Typically, spectrum was obtained by accumulating 3000 shots. Samples were prepared by dried droplet method. Sample was diluted in 50% ACN ,0.1% TFA (10 pmol/ul). A saturated matrix solution DHB (50%ACN ,0.1% TFA) was prepared. This matrix solution was mixed in equal volumes with the sample solution. The mixture was pipetted on the target (1 µl) and dried at ambient temperature. Internal standard (Mw 3765) was added to enable quantitation. Each measurement was repeated 6-times, peak area of the peptide was related to the internal standard and mean ± SD values were plotted. The experiment was repeated twice.

To monitor plasma enzymatic activity, propantheline (Sigma Aldrich) was used as a positive control. Plasma proteins were precipitated with a two-fold excess of methanol (1:2 v/v). For Echo-MS analysis (Triple Q 6500+,Sciex) 10 μL of the supernatant was further diluted with 40 μL of 30% MeOH in a 384-well Echo-MS compatible plate to ensure optimal acoustic ejection. This dualplatform approach enabled both sensitive peptide detection and rapid verification of the assay's performance.

### 11. Molecular modeling

The AlphaFold3 (Ref.[10]) predicted structure of the IGFBP-5$^{139\text{-}171\text{+}Y}$ substrate (peptide **1**) (pTM = 0.72, including c-terminal Tyr added for experimental reasons) was aligned with the experimental structure of the complex (PDB ID: 7UFG) to obtain complete peptide **1**:PAPP-A model. To reduce computational cost while preserving biological relevance, the PAPP-A was trimmed to retain only the regions involved in substrate binding. This reduced the original 1,627-residue enzyme to a 386-residue fragment, which included all functionally important areas highlighted in the sequence.

<u>Enzyme -sequence</u>:

```
MRLWSWVLHLGLLSAALGCGLAERPRRARRDPRAGRPPRPAAGPATCATRAARGRRASPPPPPPPGGAWEA
VRVPRRRQQREARGATEEPSPPSRALYFSGRGEQLRLRADLELPRDAFTLQVWLRAEGGQRSPAVITGLYD
KCSYISRDRGWVVGIHTISDQDNKDPRYFFSLKTDRARQVTTINAHRSYLPGQWVYLAATYDGQFMKLYVN
GAQVATSGEQVGGIFSPLTQKCKVLMLGGSALNHNYRGYIEHFSLWKVARTQREILSDMETHGAHTALPQL
LLQENWDNVKHAWSPMKDGSSPKVEFSNAHGFLLDTSLEPPLCGQTLCDNTEVIASYNQLSSFRQPKVVRY
RVVNLYEDDHKNPTVTREQVDFQHHQLAEAFKQYNISWELDVLEVSNSSLRRRLILANCDISKIGDENCDP
ECNHTLTGHDGGDCRHLRHPAFVKKQHNGVCDMDCNYERFNFDGGECCDPEITNVTQTCFDPDSPHRAYLD
VNELKNILKLDGSTHLNIFFAKSSEEELAGVATWPWDKEALMHLGGIVLNPSFYGMPGHTHTMIHEIGHSL
GLYHVFRGISEIQSCSDPCMETEPSFETGDLCNDTNPAPKHKSCGDPGPGNDTCGFHSFFNTPYNNFMSYA
DDDCTDSFTPNQVARMHCYLDLVYQGWQPSRKPAPVALAPQVLGHTTDSVTLEWFPPIDGHFFERELGSAC
HLCLEGRILVQYASNASSPMPCSPSGHWSPREAEGHPDVEQPCKSSVRTWSPNSAVNPHTVPPACPEPQGC
YLELEFLYPLVPESLTIWVTFVSTDWDSSGAVNDIKLLAVSGKNISLGPQNVFCDVPLTIRLWDVGEEVYG
IQIYTLDEHLEIDAAMLTSTADTPLCLQCKPLKYKVVRDPPLQMDVASILHLNRKFVDMDLNLGSVYQYWV
ITISGTEESEPSPAVTYIHGSGYCGDGIIQKDQGEQCDDMNKINGDGCSLFCRQEVSFNCIDEPSRCYFHD
GDGVCEEFEQKTSIKDCGVYTPQGFLDQWASNASVSHQDQQCPGWVIIGQPAASQVCRTKVIDLSEGISQH
AWYPCTISYPYSQLAQTTFWLRAYFSQPMVAAAVIVHLVTDGTYYGDQKQETISVQLLDTKDQSHDLGLHV
LSCRNNPLIIPVVHDLSQPFYHSQAVRVSFSSPLVAISGVALRSFDNFDPVTLSSCQRGETYSPAEQSCVH
FACEKTDCPELAVENASLNCSSSDRYHGAQCTVSCRTGYVLQIRRDDELIKSQTGPSVTVTCTEGKWNKQV
ACEPVDCSIPDHHQVYAASFSCPEGTTFGSQCSFQCRHPAQLKGNNSLLTCMEDGLWSFPEALCELMCLAP
PPVPNADLQTARCRENKHKVGSFCKYKCKPGYHVPGSSRKSKKRAFKTQCTQDGSWQEGACVPVTCDPPPP
KFHGLYQCTNGFQFNSECRIKCEDSDASQGLGSNVIHCRKDGTWNGSFHVCQEMQGQCSVPNELNSNLKLQ
CPDGYAIGSECATSCLDHNSESIILPMNVTVRDIPHWLNPTRVERVVCTAGLKWYPHPALIHCVKGCEPFM
GDNYCDAINNRAFCNYDGGDCCTSTVKTKKVTPFPMSCDLQGDCACRDPQAQEHSRKDLRGYSHG
```

The final simulated system thus consisted of the truncated PAPP-A fragment (386 residues), the peptide **1** (34 residues), and a zinc ion ($Zn^{2+}$).

The complex was prepared using the Protein Preparation Workflow[11]. The system was solvated in an orthorhombic simulation box, with at least 15 Å of water between the protein surface and each edge of the box, using the TIP3P water model[12]. To neutralize the overall charge, 15 sodium ions were added. Harmonic position restraints (10 kcal/mol/Å²) were applied to the enzyme backbone atoms to prevent unfolding or non-physiological behavior during the simulation.

The molecular dynamics (MD) simulations were performed using the OPLS4 force field[13]. Prior to the production run, the system was relaxed using the default five-step relaxation protocol in Desmond, Schrödinger[14]. The first step consisted of a 100 ps Brownian dynamics simulation under the NVT ensemble at 10 K, with harmonic restraints of 50 kcal/mol/Å² applied to all heavy atoms. This was followed by a 12 ps equilibration under the NVT ensemble, also at 10 K, using the same restraints and employing Langevin dynamics for temperature regulation. The third step was a 12 ps equilibration under the NPT ensemble at 10 K and 1 atm, introducing pressure control while maintaining the same temperature. In the fourth step, the temperature was gradually increased from 10 K to 300 K over a 12 ps NPT equilibration, again using Langevin dynamics for temperature control. The final step in the relaxation protocol was a 24 ps NPT simulation at 300 K and 1 atm, during which all restraints on the solute heavy atoms were removed. This allowed the protein to equilibrate naturally in the solvent environment under physiological temperature and pressure conditions.

The production simulations were carried out for a total of 500 ns. The temperature was maintained at 300 K using the Nosé–Hoover thermostat[15,16] with a coupling constant of 1.0 ps, and the pressure was held at 1.01325 atm using the Martyna–Tobias–Klein barostat[17] with a coupling constant of 2.0 ps. The integration time step was set to 2 fs for bonded and short-range non-bonded interactions, while long-range non-bonded interactions were updated every 6 fs using the particle mesh Ewald method. System details were saved every 100 ps, resulting in a total of 5,000 frames per simulation. To ensure statistical robustness, four independent simulations were performed, each initiated with randomized initial velocities. The resulting trajectories were analyzed to assess structural stability and identify key enzyme–substrate interactions. RMSD (Root Mean Square Deviation) metrics is summarized in **Table S1**.

Trajectory analysis was performed using the Simulation Interaction Diagram tool in Maestro 14.1 (*Schrödinger Release 2025-2: Maestro, Schrödinger, LLC, New York, NY, 2025.*, b.r.). Structural stability was evaluated by two root-mean-square deviation (RMSD) metrics calculated over the course of the simulation. The first, S-RMSD, measured the RMSD of the backbone atoms of the peptide **1**, providing insight into its conformational stability. The second, C-RMSD, tracked the RMSD of the entire complex (enzyme and substrate) aligned together, capturing the dynamic fit of the two components over time. Trajectories were sampled at 10 ps intervals.

Non-covalent interactions between the PAPP-A fragment and peptide **1** were identified using the Protein Interaction Analysis tool in Maestro 14.1 (*Schrödinger Release 2025-2: Maestro, Schrödinger, LLC, New York, NY, 2025.*, b.r.), allowing for the detection of key interacting residues at the binding interface. All reported values represent averages over four independently performed simulation replicates. Several residues (numbering refers to the IGFBP-5 sequence UniProt P24593 including the signal 20 amino-acid peptide) $R^{143}$, $K^{148}$, $K^{154}$, $R^{156}$, $K^{158}$, $Q^{162}$ and $K^{164}$ were identified as key anchoring points within the interface in agreement with cryo-EM structures **Figure S13**. Our MD simulation reviled a bend of the peptide in the active site that was secured by $Q^{162}$ and $K^{164}$ interactions that probably oriented the cleaved bond into a catalytically competent position. Raw data are available via the Open Science Framework repository: 10.17605/OSF.IO/7CZ2B.

### 12. Cryo-EM

#### 12.1: Cryo-EM sample preparation and data collection

UltrAuFoil R1.2/1.3 grids (Quantifoil) were plasma cleaned using a Henniker Plasma HPT-200 operated at 50% power for 3 min. 4 uL of samples were applied to grids in a VitroBot Mark IV (TFS) at 4 °C and 90% humidity at blot force 0 for 7 s and vitrified in liquid ethane. Data were collected on a Titan Krios G4 (ThermoFisher Scientific) equipped with a Gatan K3 camera and a Quantum-GIF energy filter. EPU 3 software was used to automate data collection. Imaging was performed in nanoprobe energy filtered zero loss mode using a 10 eV slit width. A nominal magnification of 105,000× was used, giving a calibrated pixel size of 0.833 Å. A C2 condenser aperture of 50 μm and objective aperture of 100 μm were used and the K3 camera was operated in correlated double sampling mode at a dose rate of 10.8 e−.pixel−1 s−1. Movies were collected using a 3.32 s exposure time fractionated into 50 sub-frames resulting in a total accumulated dose of 51.9 e− Å−2 per movie. 13,693 movies were collected in aberration-free image shift (AFIS) mode at a defocus range of -0.4 μm to -1.8 μm from two grids.

#### 12.2: Cryo-EM refinement

All processing was performed within cryoSPARC v4.7.1 (*76*). The movies were patch-motion corrected and patch-CTF estimation was performed. Micrographs containing thick or contaminating ice, CTF fit resolution > 4 Å or significant motion were removed, leaving 10,838 good micrographs. Particles were picked using templates generated from a low-resolution structure from a screening dataset and extracted, binned 6x (box size 100 px). Particles were 2D classified, with obvious non-protein particles and duplicates removed, giving 7.6M particles. *Ab initio* reconstruction with 4 classes was performed from 200k of the selected particles and an additional decoy map generated from the rejected particles. Heterogeneous refinement using the most complete *ab initio* map against the decoy was performed and particles corresponding to the desired 3D class were re-extracted with a pixel size of 1.66 Å (300 px box). Iterative heterogeneous refinement against two copies of the resulting high quality reconstructions was then performed, with 500k high-quality particles eventually retained for the final refinements.

These particles were re-extracted with a pixel size of 1.14 Å (440 px box) and refined using the NU-refine job to give a consensus 3 Å map. CTF refinement and reference-based motion correction were performed iteratively until resolution and map quality failed to improve the consensus. Local refinement with a mask covering one half of the complex gave the final 2.7 Å map.

#### 12.3: Model building

Modelling was initiated from the published PAPP-A IGFBP5 peptide complex (pdb: 7UFG). ChimeraX v1.10 (Ref.[18]) was used to rigid body fit the model to the final map and morph it to fit the density with adaptive restraints to the initial models using ISOLDE v1.10 (Ref.[19]). Models and restraints for pseudopeptides 3 and 4 were generated by building from scratch within ChimeraX, and restraints generated with ELBOW within phenix 2.1. The IGFBP5 peptide was mutated to the pseudopeptide sequences within coot 0.9.8.6 and further refinement of the model was completed by iterating between coot and ISOLDE. Final refinement statistics were generated using real space refinement within phenix. Atomic model and map have been deposited at the PDB and EMDB with the accession codes 45DF and EMD-83190, respectively.

#### Supporting information references


1. Dingwalla, J. G., Ehrenfreund, J. & Hall, R. G. Diethoxymethylphosphonites and phosphinates. Intermediates for thesynthesis of α,β- and X aminoalkylphosphonous acids. *Tetrahedron* **45**, 3787-3808 (1989).
2. Pícha, J. *et al.*, Efficient synthesis of phosphonodepsipeptides derived from norleucine. *Tetrahedron* **65**, 6090-6103 (2009).
3. Shin, I. *et al.*, Synthesis of optically active phthaloyl D-aminooxy acids from L-amino acids or L-hydroxy acids as building blocks for the preparation of aminooxy peptides. *J. Org. Chem.* **65**, 7667-7675 (2000).
4. Pícha, J., Buděšínský, M., Macháčková, K., Collinsová, M. & Jiráček, J. Optimized syntheses of Fmoc azido amino acids for the preparation of azidopeptides. *J. Pept. Sci.* **23**, 202-214 (2017).
5. Yiotakis, A., Vassiliou, S., Jiráček, J. & Dive, V. Protection of the Hydroxyphosphinyl Function of Phosphinic Dipeptides by Adamantyl. Application to the Solid-Phase Synthesis of Phosphinic Peptides. *J. Org.Chem.* **61**, 6601-6605 (1996).
6. Campagne, J.-M., Coste, J., Guillou, L., Heitz, A. & Jouin, P. Solid phase synthesis of phosphinic peptides. *Tetrahedron Lett.* **34**, 4181-4184 (1993).
7. Mikkelsen, J. H. *et al.*, Inhibition of the proteolytic activity of pregnancy-associated plasma protein-A by targeting substrate exosite binding. *J. Biol. Chem.* **283**, 16772-16780 (2008).
8. Westerfield, M. *The Zebrafish Book: A Guide for the Laboratory Use of Zebrafish (Danio Rerio)* (University of Oregon Press, 2000).

9. Dai, W. *et al.*, Calcium deficiency-induced and TRP channel-regulated IGF1R-PI3K-Akt signaling regulates abnormal epithelial cell proliferation. *Cell Death Differ.* **21**, 568-581 (2014).
10. Abramson, J. *et al.*, Accurate structure prediction of biomolecular interactions with AlphaFold 3. *Nature* **630**, 493-500 (2024).
11. Sastry, G. M., Adzhigirey, M., Day, T., Annabhimoju, R. & Sherman, W. Protein and ligand preparation: parameters, protocols, and influence on virtual screening enrichments. *J Comput Aided Mol Des* **27**, 221-234 (2013).
12. Jorgensen, W. L., Chandrasekhar, J., Madura, J. D., Impey, R. W., Klein, M. L. Comparison of simple potential functions for simulating liquid water. *J. Chem. Phys.***79**, 926-935 (1983).
13. Lu, C. *et al.*, OPLS4: Improving force field accuracy on challenging regimes of chemical space. *J. Chem. Theor. Comput.* **17**, 4291-4300 (2021).
14. Bowers, K. J. *et al.*, Scalable algorithms for molecular dynamics simulations on commodity clusters. *ACM/IEEE SC 2006 Conference (SC'06)*, 43-43 (2006).
15. Nosé, S. A unified formulation of the constant temperature molecular dynamics methods. *J. Chem. Phys.s* **81**, 511-519 (1984).
16. Hoover, W. G. Canonical dynamics: Equilibrium phase-space distributions. *Phys. Rev. A* **31**, 1695-1697 (1985).
17. Martyna, G. J., Tobias, D. J., Klein, M. L. Constant pressure molecular dynamics algorithms. *J. Chem. Phys.* **101**, 4177-4189 (1994).
18. Meng, E. C. *et al.*, UCSF ChimeraX: Tools for structure building and analysis. *Protein Sci.* **32**, e4792 (2023).
19. Croll, T. ISOLDE: a physically realistic environment for model building into low-resolution electron-density maps. *Acta Crystallogr. D* **74**, 519-530 (2018).
